# Beyond EEG Onset Transients: Sensitisation and Habituation of Hyper-excitation to Constant Presentation and Offset of Pattern-Glare Stimuli

**DOI:** 10.1101/2024.09.13.612622

**Authors:** Tom Jefferis, Cihan Dogan, Claire E. Miller, Maria Karathanou, Austyn J. Tempesta, Andrew J. Schofield, Howard Bowman

**Author notes:** **Correspondence** Professor Howard Bowman, School of Psychology and Computer Science, University of Birmingham, Edgbaston, Birmingham, B15 2TT, UK.

## Abstract

Pattern-glare, characterised by visual distortions, discomfort, and stress when viewing striped patterns, has been associated with cortical hyperexcitability, particularly in individuals with visually induced epilepsy, migraines, and visual stress. While previous studies have explored the onset transients of such stimuli, this research investigates the sensitisation and habituation effect to the constant presentation and offset of pattern-glare stimuli. We analysed the temporal characteristics of the Event Related Potentials (ERPs) in healthy participants to find correlates of cortical hyperexcitability over two time granularities: a fine granularity (over seconds) and a coarser granularity (across the time-course of the entire experiment). We looked for habituation and sensitization effects across these time periods. Our results suggest that brain responses to pattern-glare stimuli are correlated to participants’ sensitivity to visual discomfort with statistically significant effects observed for this factor and for its interaction with changes over time. This study improves our understanding of how the brain adapts to persistent visual stimuli and provides insights that may inform treatments for conditions like migraine and epilepsy.

## 1 Introduction

**Pattern-glare** is characterised by symptoms of perceptual distortions, discomfort and visual stress when viewing striped patterns (Evans and Stevenson, 2008). Since striped patterns rarely occur in nature (particularly at the spatial frequencies that aggravate the brain), it is thought that such stimuli take the brain beyond the processing regime for which it has evolved, and the observed hyper-excitation response to such stimuli reflects this evolutionary aberrance (Fernandez and Wilkins, 2008; Monger et al., 2015). Some individuals are more affected by these patterns, particularly those who suffer from visually induced epilepsy, migraines or visual stress. As early as 1935, researchers documented the effects of bright lights or patterns on people who suffer from migraines, finding a small number of individuals who had episodes that could be triggered by these patterns or lights (Turville, 1935).

Thus, pattern-glare is a well-established phenomenon with sustained research dating from the 1980’s on the effects of striped patterns on the visual system of humans. The pattern-glare test was formalised in 2001 to allow practitioners to assess individual’s susceptibility to pattern-glare (Wilkins and Evans, 2001). The test is designed to elicit visual distortions and discomfort. Individuals report the level of distortion and discomfort on a questionnaire with a set of yes/no questions. For an individual to be clinically diagnosed with pattern-glare, they must score in or above the 95^th^ percentile in the test. However, there is no objective measure to assess the level of distortion an individual is experiencing; answers to questionnaires are subjective and susceptible to response bias.

More recently, researchers have shown that visual gratings can induce pattern-glare, the brain correlates of which can be detected at the scalp level using EEG, with patterns close to 3 cycles per degree (c/deg)^1^ eliciting the greatest response (Adjamian et al., 2004; Aurora and Wilkinson, 2007). More recently there have been studies linking electrophysiological correlates of hyper-excitation, with Fong et al. finding differences between the responses of migraine sufferers and controls in the time-domain. The group of migraineurs had an enhanced N2 deflection that could be driven by hyperexcitation (Fong et al., 2020). Migraineurs sensitivity was corroborated with findings from Harle et al. who found that migraine sufferers saw significantly more illusions when viewing striped patters and would be more likely to select a coloured filter to aid with visual comfort (Harle et al., 2006). In a review of literature, Aurora and Wilkinson suggested that repetitive patterns, particularly those closely related to the high contrast striped or checkerboard patterns were linked to visual triggers of migraines (Aurora and Wilkinson, 2007). Additionally, we provided evidence that susceptibility to headaches predicts the absence of the N1 ERP (Event Related Potential) component, i.e. participants with increased headache susceptibility exhibited a smaller N1 and thus a more positive going response to the stimulus at the aggravating spatial frequency (Tempesta et al., 2021). However, Tempesta et al. did not consider how the ERP response changes through time, thus neither sensitisation or habituation effects were identified, patterns of change that are likely to be important in understanding migraine, epilepsy and visual stress.

### Change through time

Many studies have found a dysfunctional habituation system for migraine and epilepsy conditions, disorders that have been linked to a susceptibility to pattern glare (Brazzo et al., 2011; Coppola et al., 2009). As discussed, the possibility of a dysfunctional habituation system was not explored in Tempesta; an aim of this paper is to address this shortcoming. We will do this by analysing how the ERP response to pattern-glare stimuli changes through both their (short-term) repetition and across the time-course of the whole experiment.

Importantly, treatments for visually-induced migraine and epilepsy could be informed by studying how the brain habituates to pattern-glare stimuli, where that habituation could be over short or long time frames. Accordingly, our experiment has a two-by-two structure, with two types of change through time: habituation (exponentially decreasing response through time) and sensitisation (exponentially increasing response through time), and two granularities of time: fine/short-term (through trains of stimulus presentations) and coarse/long-term (through the course of the entire experiment). We also investigate how these types and granularities of change through time are modulated by participants’ state and trait sensitivities to relevant conditions, such as headache, visual stress and discomfort induced by viewing pattern-glare stimuli. These three sensitivities will be called *factors* in this work, since they emerged from a factor analysis. Although, in this paper, our main focus will be on the discomfort factor, since it gives us our strongest effects. This may be because it is largely a state measure, reflecting experience during the experiment, rather than a trait measure, reflecting participants’ (more subjective) view of their long-term susceptibility to headaches, visual hallucinations, etc. This said, we do report a small number of effects on the visual stress and headache factors.

### DC-shift and Offset

The vast majority of human electrophysiology experiments consider the transients associated with the onset of a new stimulus (Luck, 2014). However, this focus ignores two other aspects of the brain’s processing of stimuli: 1) the stationary or evolving response to a continuously-presented stimulus (because of the pattern of responding, we will call this the *DC-shift* period) and 2) the transients associated with the brain’s return to stasis when a stimulus is turned off. In the context of studying hyper-excitation, these two aspects of brain processing can be revealing. In particular, they could indicate the inhibitory mechanisms employed in the brain, since these would be engaged during a period of constant stimulation and may become unopposed at stimulus offset, enabling electrophysiological correlates of inhibition to be observed (Thompson and Burr, 2009).

*1) DC-shift:* consistent with the focus on onset transients, most ERP studies only present the stimulus for a short period of time, e.g. a few hundreds of milliseconds. However, a stimulus that stays on for longer will continue to drive the brain, perhaps particularly a brain susceptible to hyper-excitation. To observe this brain response, we presented stimuli for three seconds. A basic feature we observe is that the electrical response does reach a somewhat stationary state with constant stimulation, but with a baseline shift relative to the pre-stimulus period; see Figure 10C. In electrical engineering terms, this new baseline could be considered a direct-current effect, hence the term DC-shift.
*2) Offset-transients:* we also seek to understand the return to stability from the pattern-glare stimulus. Thus, we analyse the offset to stimuli, ERP effects that very few researchers have previously considered (Bendixen et al., 2006; Marmoy et al., 2021; O’reilly, 2019; Takahashi et al., 2004) and particularly not the offset transients to pattern-glare stimuli.

The effects we identify could serve as biomarkers of hyperexcitation and visual stress in the DC-shift and offset period of the pattern-glare experiment.

## 2 Methods

The three stimuli were made with the Psychophysics Toolbox in MATLAB (Brainard, 1997; Kleiner et al., 2007; Pelli, 1997) and were based on the stimuli from the pattern-glare test (Wilkins and Evans, 2001). The three stimuli were made from horizontal square-wave gratings at three different spatial frequencies (SF) (thin = 12 c/deg, medium = 3 c/deg, thick = 0.37 c/deg) with 75% contrast as illustrated in Figure 1. The stimuli were displayed within a circle with diameter 15.2 deg on a Samsung 932BF LCD monitor (Resolution = 1280 x 1024 pixels) at a viewing distance of 86 cm. Pattern 1 (thick) is meant to be a control for low SF and is not supposed to trigger distortions in most participants. However, it is useful in detecting ‘which patients may be highly suggestible and may respond yes to any question about visual perception distortions’ (Evans and Stevenson, 2008). Pattern 2 (medium) is the only relevant clinical stimulus falling between SF’s 1-4, which are known to elicit migraines and epileptic seizures (Braithwaite et al., 2013; Wilkins, 2016). Pattern 3 (thin) is a control for poor convergence and accommodation. Those with poor convergence and/or accommodation will see distortions in this stimulus reflecting optical rather than neurological factors (Conlon et al., 2001). Therefore, for any effect that simply reflects spatial frequency, rather than hyper-excitation, the medium stimulus should have a response close to the mean of the thin and thick stimuli.

**Figure 1:**
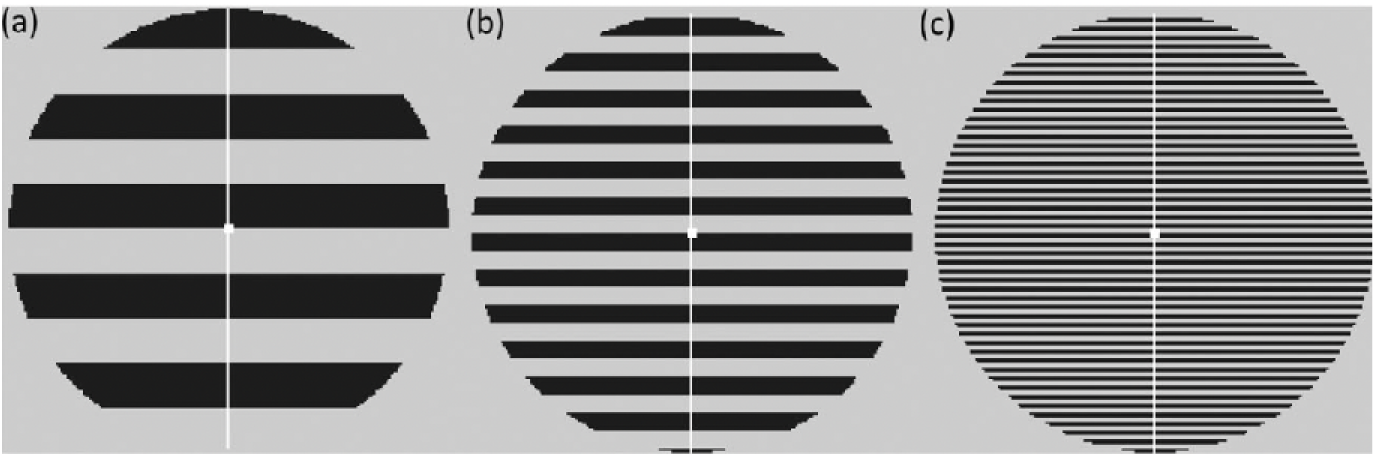
Illustration of the three stimuli used in the pattern-glare test and our experiment. a) First control pattern, 0.5 c/deg (thick); b) clinically-relevant pattern, 3c/deg (medium); and c) third control pattern, 12c/deg (thin). Here the stripes have been scaled so as to avoid distortions in print but are representative of the stimuli shown to the participants.

### 2.1 Data Collection

Forty participants were recruited at the University of Birmingham, all giving consent and compensated with £24 for participating. None of the participants had prior history of neurological, psychiatric, or psychological conditions as well as no history of unconsciousness, convulsions, or epilepsy. Two of the participants were excluded before pre-processing: one withdrew from the experiment before completion, while an equipment failure meant that only a partial dataset was recorded for the other.

### 2.2 Questionnaires

Multiple questionnaires were used to assess the participants’ headache histories and proneness to suffer from visual stress. These questionnaires included the cortical hyperexcitability index (CHi), used to asses participants’ visual discomfort and visual aura (Braithwaite et al., 2015), and the visual discomfort scale (VDS), which was used for assessing participants’ visual discomfort and side effects to the pattern-glare experiment (Conlon et al., 1999). For headache symptoms, we selected questions from a more general, headache and general health questionnaire (HGHQ). The headache criteria specified by the International Headache Society (Arnold, 2018) were not used as these are criteria for a clinical diagnosis and do not provide scale measures of headache proneness. The criteria heavily rely on factors such as: headache intensity, nature, duration, and frequency. All these factors were recorded by the HGHQ.

### 2.3 Procedure

Following EEG cap setup, there was a five-minute resting period before the main experiment began. The main experiment consisted of three blocks (which we call partitions), with six trials per stimulus type (thin, medium, thick) for a total of eighteen trials per stimulus type across the whole experiment. Every trial began with a fixation cross for four seconds, followed by seven to nine onsets of the same stimulus, which were presented on grey backgrounds with the same space-averaged luminance, each of which stayed on the screen for three seconds. This was followed by another fixation cross for between 1 to 1.4 seconds; see Figure 2. Following each trial, the participants rated their degree of visual discomfort on a five-point scale (1= comfortable, 5 = extreme discomfort) and recorded how many times they believed the stimulus was shown to assess their attentiveness. At the end of each block (partition), there was another five-minute resting period where the participant was instructed to rest and close their eyes.

**Figure 2:**
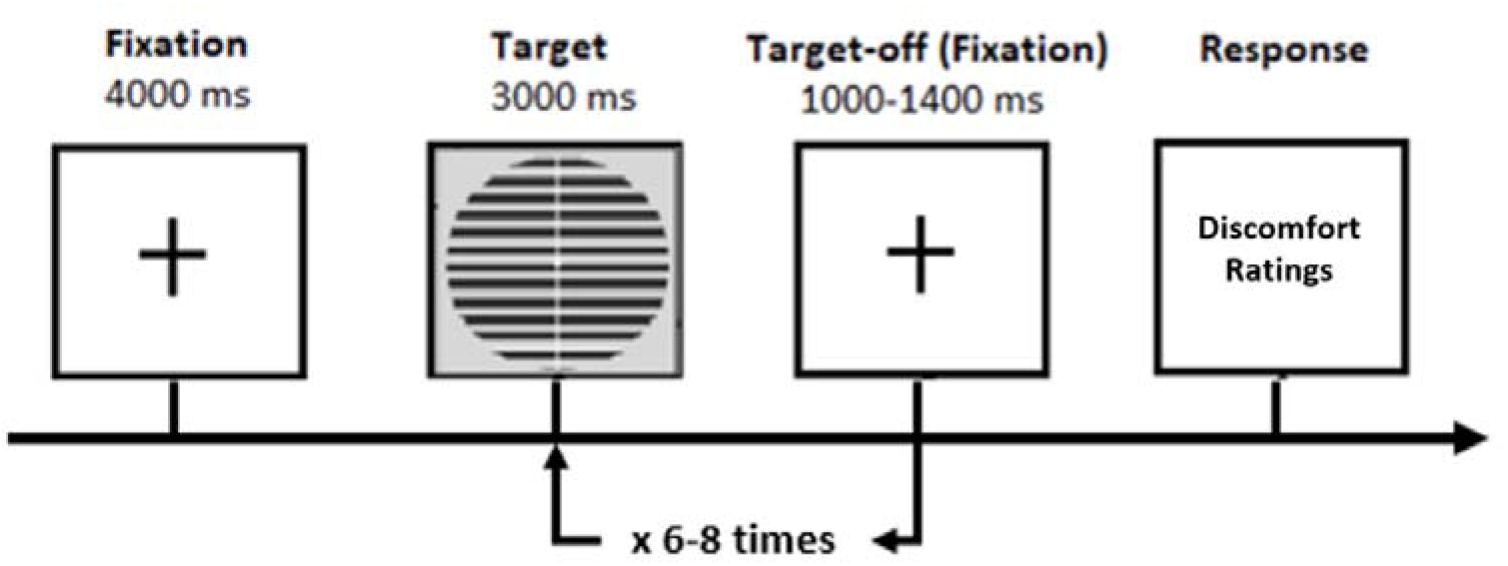
Schematic representation of a single trial. This sequence was repeated 6 times per stimulus type to complete one block (partition) of experiments.

### 2.4 Factor Analysis

Working with the 38 participants who completed the study, we computed the mean discomfort ratings for each participant and stimulus type across the three blocks (partitions). As discomfort ratings tend to co-vary across the stimulus types, the discomfort index for each participant was computed by subtracting the mean discomfort ratings for the thin and thick stimuli from the scores of the medium stimuli. By doing this for each participant, we would be able to identify participants who found the medium stimulus more uncomfortable relative to the two control stimuli. The overall scores for the CHi and VDS were computed according to their instructions. Lastly, the factors from the HGHQ were extracted including headache frequency, intensity and duration, and the experience of sensory aura. There was a total of seven measures, which had very different ranges, so they were standardised before entering them to the factor analysis stage. This step identified three factors based on a Scree plot analysis. Following a Varimax rotation, three factors were identified as, visual stress, headache, and discomfort (see appendix section: 9.1.1 for full analysis on the identified factors). This paper focuses on the discomfort factor, since it gave us our most substantial effects. Although, we also present a small number of weaker effects associated with visual stress and headache. We mainly focus on effects associated with orthogonalized regressors, although we do also summarise the same effects without orthogonalization. The factors were orthogonalized in the order visual stress, headache, and discomfort using the Gram-Schmidt method (GSM) (Arfken, 1985); this was because visual stress was highest in the Scree plot (highest Eigenvalue), headache was next and discomfort third.

### 2.5 Data Pre-processing

The EEG data was first decimated from the recording frequency of 2048Hz to 512Hz using the Biosemi toolbox and EEGLAB (Delorme and Makeig, 2004). Eye blink artifacts were removed using independent component analysis (ICA) and the dataset was then recompiled.

The Fieldtrip toolbox (version 20210921) (Oostenveld et al., 2010) was then used for the rest of the pre-processing stages. The signals were bandpass filtered using an FIR filter with a range of 0.1 to 30Hz using a Hanning window. The data for each onset were epoched between -200ms and 4000ms relative to stimulus onset, and referenced to the average of all electrodes.

The two different windows that we analysed, DC-shift and Offset, employed different baselining windows. The DC-shift period was baselined relative to the -200 - 0ms time window just before stimulus onset. Our analysis of the DC-shift period seeks to identify stationary as well as non-stationary differences between the three stimuli during this period, Consequently, it is appropriate to baseline correct to the nearest time-period without transients before the DC-shift period, certainly rather than at the start of the DC-shift period itself, where there are significant transients.

In contrast, baseline correction for the Offset period was carried out in the period 2800ms – 3000ms after stimulus onset. This period was chosen because it immediately preceded stimulus offset, and, due to the smoothness of the (DC-shift) period, suggesting that the time-series were close to stationary at this point. Additionally, had we used the -200ms – 0ms window, as we did for the DC-shift period, the signal could have drifted substantially by the time of the offset. Further, the DC-Shift effect itself meant that waveforms had not returned to their original baseline and this deviation from baseline was itself correlated with our factors and would thus have confounded a separate analysis of the offset transient. Thus this choice of offset baselining enables us to directly compare the return to stasis of the three stimulus types: thick, medium and thin.

To confirm that this baselining window should not itself confound the results, we fit a line to the grand average ERPs in the baselining windows. Figure 3, illustrates these grand averaged ERPs for all three stimulus types at electrode Oz (A23 on the biosemi 128 EEG cap). We also fit a line to the pattern glare index (PGI; see Equation 2) of each participant for the standard baselining window of -200ms – 0ms and 2800ms – 3000ms. This index (explained in full later) is a measure of the relative response to the clinically relevant medium spatial frequency. We then ran a two-sample t-test on the gradients for both baseline periods using the PGI. The p value of this test was p = 1.000, meaning there was no significant difference in gradients between the two windows.

**Figure 3:**
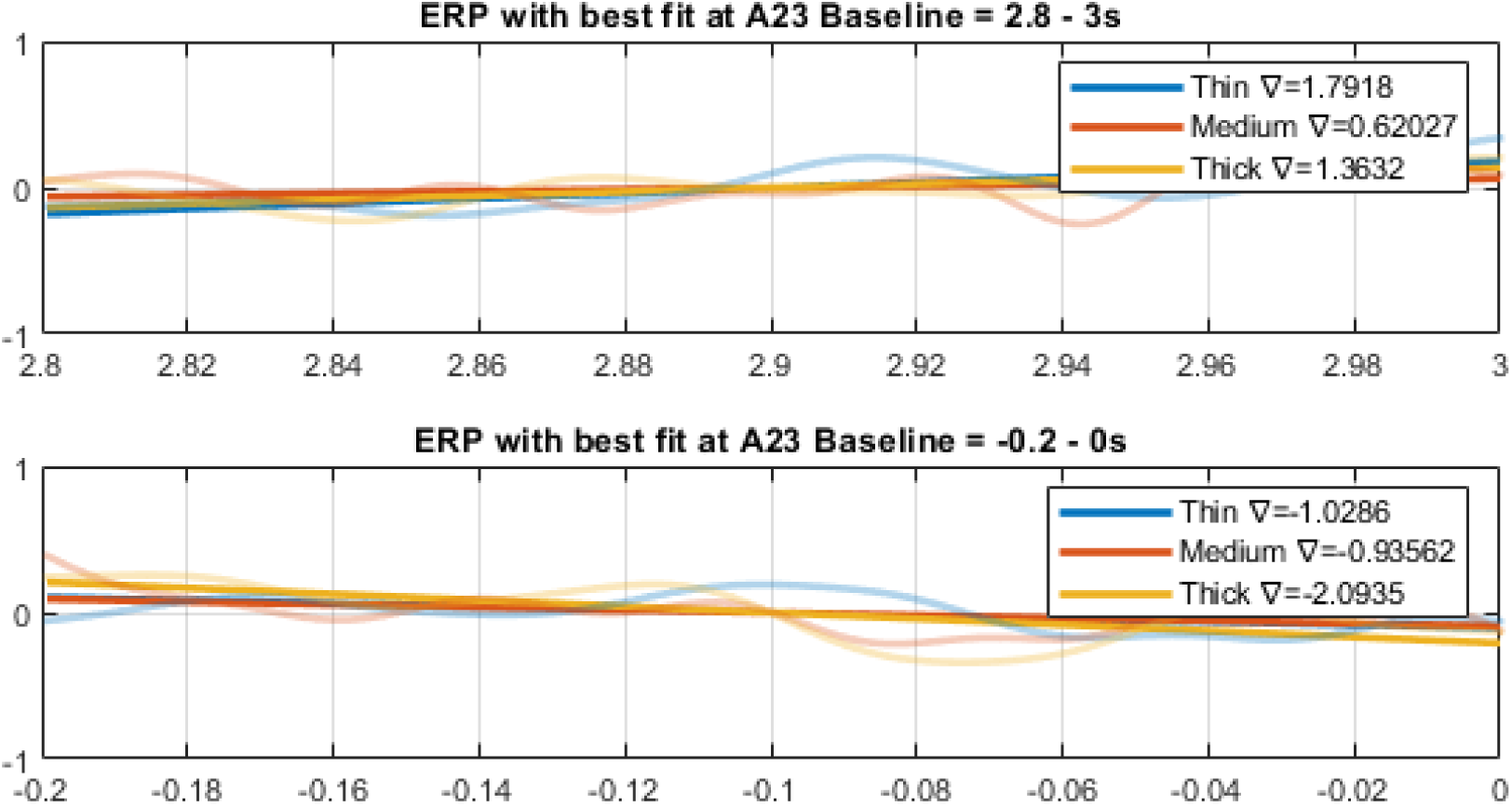
Baselining windows used in analyses. The top plot is the baselining window used for this offset analysis and the bottom plot is the baselining window used for the DC-shift analysis.

Thresholding was then applied with a ± 100µV threshold to remove any remaining large artifacts in the data. Participants were then either accepted or rejected based on having a minimum of 20% useable trials for each condition after thresholding, in line with suggestions by Luck (Luck, 2014), leaving us with 34 participants for the basic average of onsets analysis, 32 for the coarse and fine time granularity analysis and 31 for the three way analysis (types of analyses are outlined in section 2.7).

A number of different regressions were performed. These fell into four types, according to the form of the dependent variable employed. In the first type, the dependent-variable was the average of the onsets 2-8; onset 1 was excluded from the average as it could contain an effect of surprise (all three stimuli were equally likely at onset one, but the same stimulus was presented for the remaining onsets). Onset 9 was also excluded as it was infrequently shown to participants, and was thus considered too noisy. This first type gave us the mean/intercept and Discomfort independent variable regressors discussed in section 2.7.1. In the second type, the dependent-variable was the coarse time granularity, in which the experiment was split into three partitions in line with the experimental blocks, with onsets averaged in each partition. This gives us an idea of how the participants’ brain response changed throughout the duration of the whole experiment. This second type gave us the coarse change through time independent variable regressors discussed in section 2.7.2. In the third type, the dependent-variable was the fine time granularity, which compares onsets. Pairs of onsets were averaged together, onsets 2 and 3; onsets 4 and 5; and onsets 6 and 7. These averages were then compared to give us an idea of how the participants’ brain response changed within each trial. This third type gave us the fine change through time independent variable regressors discussed in section 2.7.2. In the final type, the dependent-variable enabled us to perform the three-way interaction. This had a similar form to that of the partitions dependent-variable however, in this case, each partition was split into three onset-pairs (2:3, 4:5 and 6:7). In this way, we analyse the participants’ progression through both the coarse and fine time granularities. This fourth type gave us the three-way interaction independent variable regressors discussed in section 2.7.2.1.

### 2.6 Temporal Region of Interest

The offset analysis window was determined through a data driven approach, using the fully flattened average (Bowman et al., 2020; Brooks et al., 2017) in combination with the global field power (GFP) (Murray et al., 2008; Skrandies, 1990). The method involved calculating the fully flattened average for all electrodes, and then the global field power using the equation:

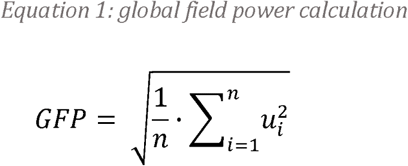

Thus, the GFP is the root mean squared across all electrodes (of which there are *n* in equation 1) at a point in time. This gives us an indicator of the global power (i.e. squared distance from zero) at any time point, which allows us to take regions of analysis based on the periods with the most activity. Unconstrained window selection runs the risk of inflating the false positive rate, however, since the fully flattened average does not discriminate across conditions, it does not cause such an inflation (Bowman et al., 2020; Brooks et al., 2017).

One reason for subdividing the offset time period is because it contains peaks with very different smoothness. Cluster inference procedures can show reduced sensitivity to clusters in rougher regions of the volume, because the (null hypothesis) permutation distribution is dominated by clusters found in smoother regions, which will typically be bigger.

There were three temporal windows in the offset. These were identified by taking the N largest peaks (N = 10) over the mean GFP across the whole period, then for each peak, the previous trough, which fell below the GFP mean, was found. This point would be the start of the window. This was repeated for the following trough after the peak (for the end of the window), and then for every peak found. The overlapping windows were then either removed or cut to end as the next window started, windows less than 10% of the length of the signal were removed. The windows found were: 3.09 – 3.18s, 3.18 - 3.45s, 3.45 - 3.83s shown in Figure 4. In addition to these three windows, the time period 3.09 – 3.99s was included to be able to see any potential effects that span multiple windows.

**Figure 4:**
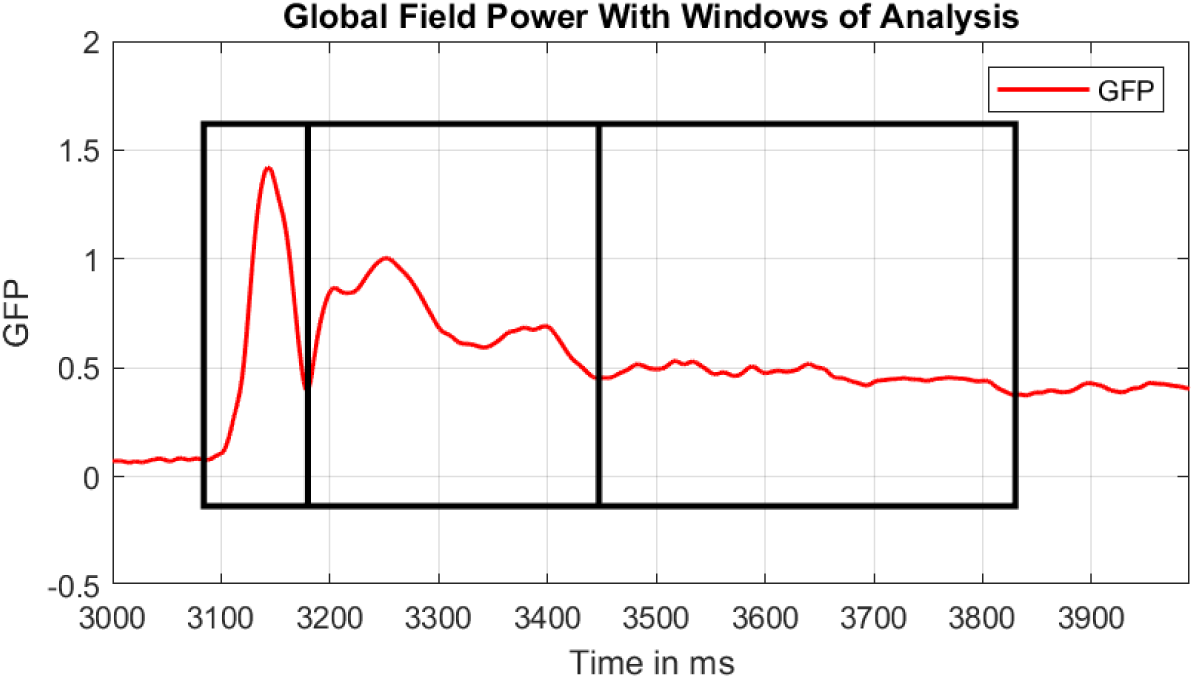
Offset global field power with windows of interest, X axis represents time in ms and Y axis represents GFP. The three windows of analysis are shown in black boxes.

### 2.7 Mass univariate analysis

A mass univariate analysis (MUA) is the analysis of a large number of simultaneously measured dependent variables (e.g. voxels or samples) via the performance of the same univariate hypothesis tests (e.g., t-tests) across all of those dependent variables. This method provides a powerful correction for multiple comparisons. An MUA was conducted on the participants’ ERPs in FieldTrip, using FieldTrip’s cluster-based permutation test method. The significance probabilities of the permutation tests were calculated using the Monte Carlo method, using a significance threshold for FWE-correction of 0.05, with cluster forming threshold of 0.025. All tests were run for 10,000 permutations, but then effects with p values close to 0.05 (i.e. where passing the 0.05 threshold is uncertain) were run again with 25,000 permutations to obtain a more accurate p-value estimate. The MUA was conducted on what we call the Pattern Glare Index (PGI). This index enables us to focus on parts of the data volume where the medium stimulus exhibited more extreme responses than the two control stimuli. The equation used for the PGI calculation is:

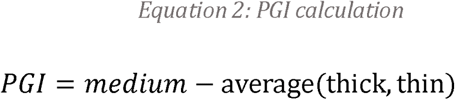

#### 2.7.1 Basic (non-temporal) analyses

For the first dependent-variable-type, we ran a MUA, with a regression with intercept^2^ and factor scores; see Figure 5 and Figure 6 (shown for discomfort). Since all the factor regressors are mean centred, the intercept becomes the mean of the basic stimulus effect on the PGI. This analysis identifies points in the data volume in which medium is more extreme than the two control stimuli (thin and thick), without considering a participant’s proneness to visual stress, headache or discomfort, or also their sensitivity to change through time.

**Figure 5:**
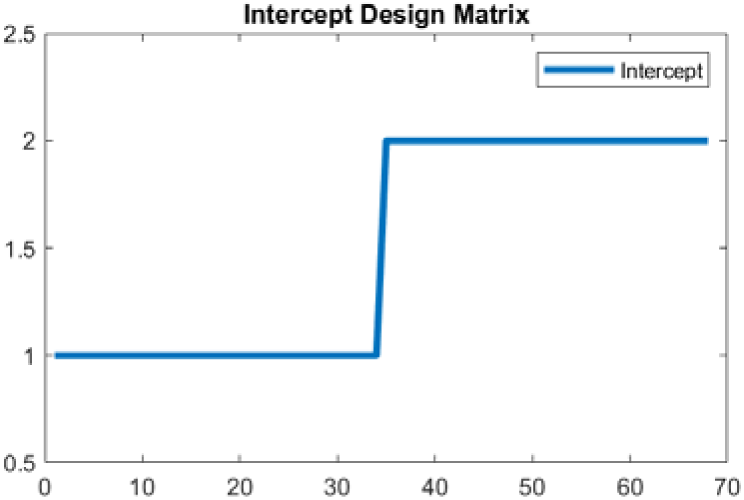
Intercept regressor for the average of onsets 2-8. X axis represents participants, Y axis represents the scores entered in the regressor for each participant. Participants labelled 35 and above are null data, with Y variable set as 2 for the MUA to run.

**Figure 6:**
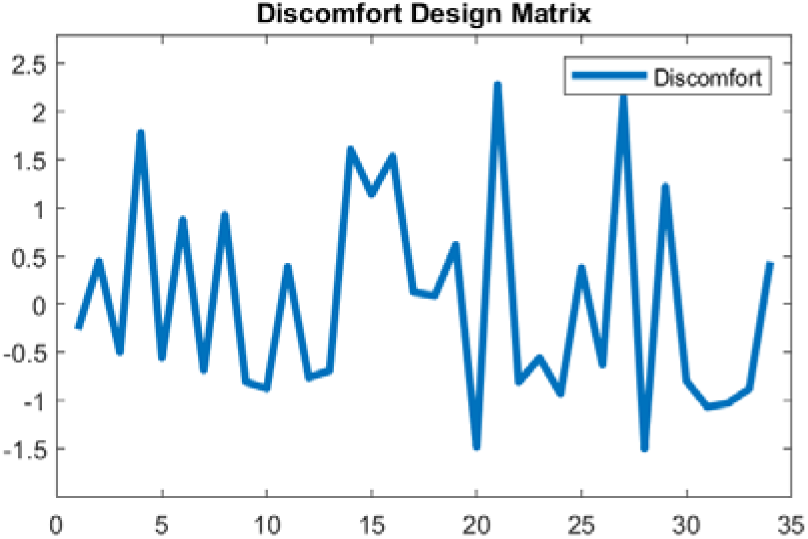
Discomfort regressor for the average of onsets 2-8. X axis represents participants, Y axis represents the scores entered in the regressor for each participant.

#### 2.7.2 Change through time

Two additional steps were taken to generate the factors x time granularity regressors; i.e. formulate the interaction between factor and change through time. Firstly, all the factor scores were shifted such that all the scores were positive or zero, this was done by subtracting the minimum from all the scores. These scores were then duplicated three times (one for each time period, i.e. block or onset-pair) and then multiplied by an exponential change pattern, which we explain formally in the appendix (See Figure 25 and section 9.1). Lastly, the regressor was mean centred.

The resulting two-way interaction regressors are illustrated in Figure 7, with a decreasing pattern, characteristic of habituation, through the partitions on the left and an increasing pattern, characteristic of sensitisation, through the partitions on the right. In particular, the exponential change through time periods should be evident. For example, in the decreasing pattern on the left, the mean across participants in partition 1, i.e. across the blue line, is substantially higher than in partition 2, i.e. across the red line, which is somewhat higher than in partition 3, i.e. across the orange line. Additionally, the extent to which the variability in the discomfort factor is exhibited also varies with partitions. That is, in partition one, we are looking for brain responses that vary very substantially across participants, with those high on the discomfort factor exhibiting substantially higher responses than those low on the factor. Although still present, this variability is substantially reduced in partition 2 and completely absent in partition 3, i.e. by the end of the experiment, there is full habituation, with the hyper-excitation (which was differential across the discomfort factor), effectively, gone. These factor by decrease and factor by increase interaction regressors will also be used unchanged to explore the pattern of habituation and sensitisation across onset pairs, our finer granularity of change through time.

**Figure 7:**
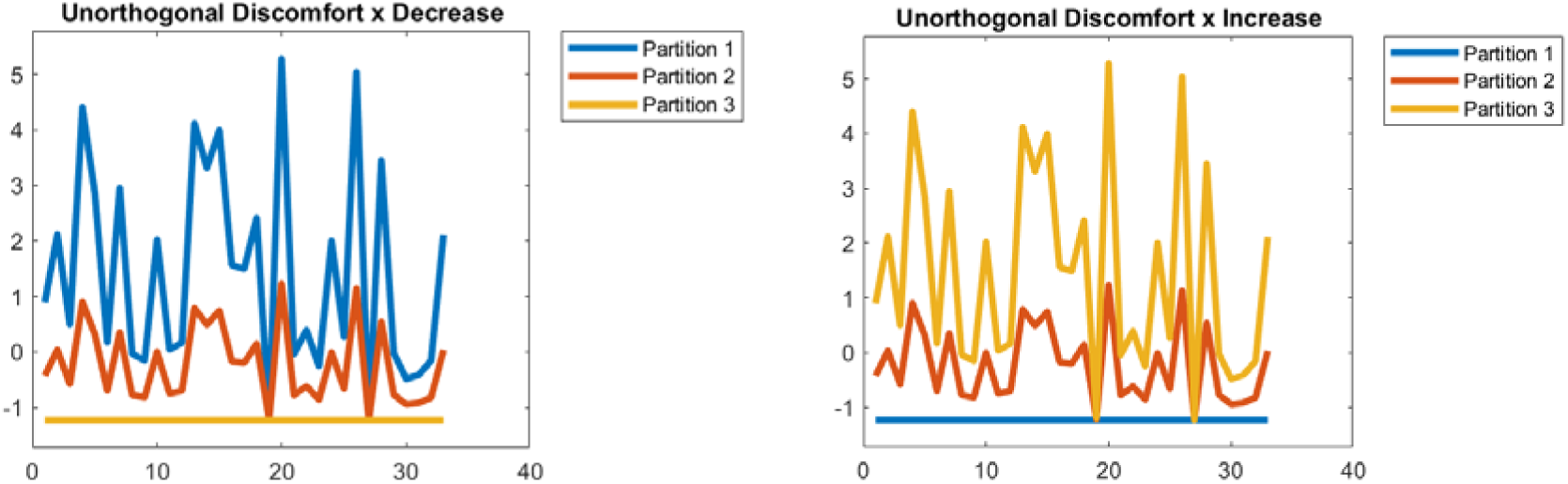
Left: regressor for discomfort by decrease across the blocks/partitions before orthogonalization. X axis represents participants, Y axis represents the scores of each participant. Right: regressor for discomfort by increase across the blocks/partitions before orthogonalizatio n. X axis represents participants, Y axis represents the scores of each participant.

The second step was orthogonalizing the three-resulting interaction regressors (one for each factor, visual stress, headache, discomfort), which was again performed using the Gram-Schmidt algorithm; see Figure 8. The other two interaction regressors were orthogonalized with respect to the Visual stress by Decrease interaction regressor and themselves, using the Gram–Schmidt process (Arfken, 1985). This sequence of orthogonalizations was chosen because Visual Stress was the factor that obtained the strongest loading in our factor analysis and thus would naturally be preserved unchanged by the orthogonalization. We ran another analysis where the interaction regressors were not orthogonalized and these are presented as exploratory results in the appendix (see section 9.2).

**Figure 8:**
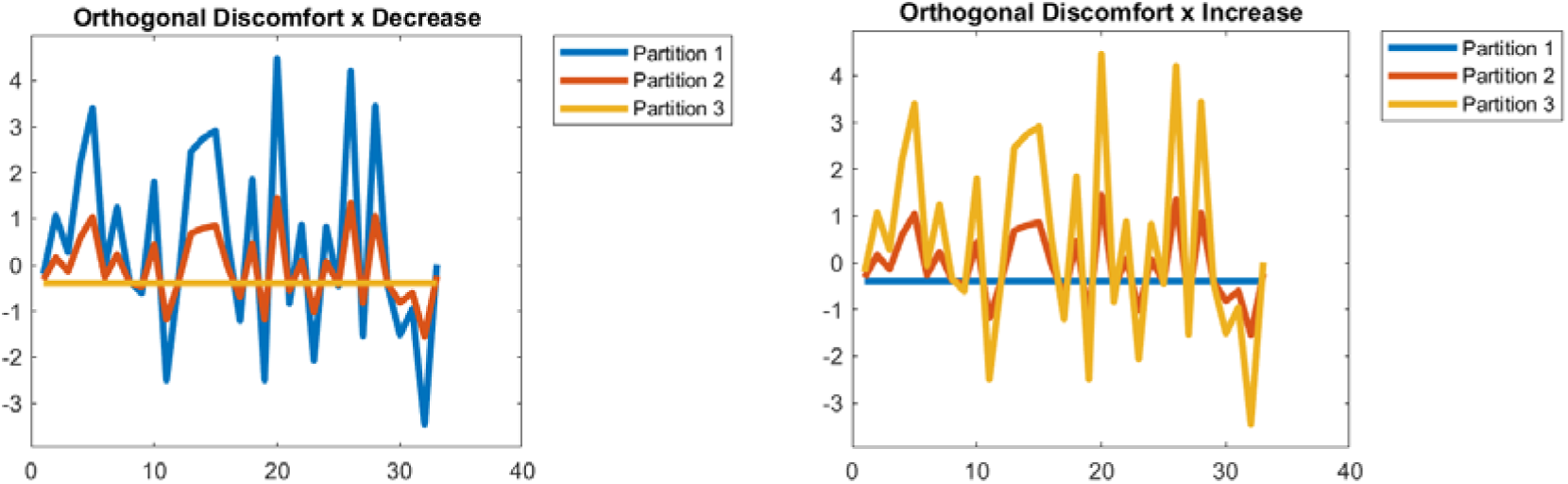
Left: regressor for discomfort by decrease a cross the blocks/partitions after orthogonalization. X axis represents participants, Y axis represents the scores of each participant. Right: regressor for discomfort by sensitisation across the blocks/partitions after orthogonalization. X axis represents participants, Y axis represents the scores of each participant.

##### 2.7.2.1 Three-way interaction

Each three-way interaction regressor (one for each factor) was constructed by taking the factor regressor from the basic analysis and duplicating this three times (one for each onset-pair); these three were multiplied by the exponential change to form the first change through time effect. Then, these three resulting regressors were appended to form one regressor, which were again duplicated three times (one for each partition). Each of these three (factor by change through onset-pairs) regressors were multiplied by the exponential change to form the second change through time effect. The final step of this process was orthogonalizing the three-resulting interaction regressors (one for each factor, visual stress, headache, discomfort), which was again performed using the Gram-Schmidt algorithm. Figure 9 shows an example (orthogonalized) three-way interaction regressor (note this is not the regressor we present results for, since the three-way that came out exhibited a decrease across onsets and an increase across partitions). We ran another analysis where the interaction regressors were not orthogonalized however, no clusters were identified in the MUA and they are thus not presented.

**Figure 9:**
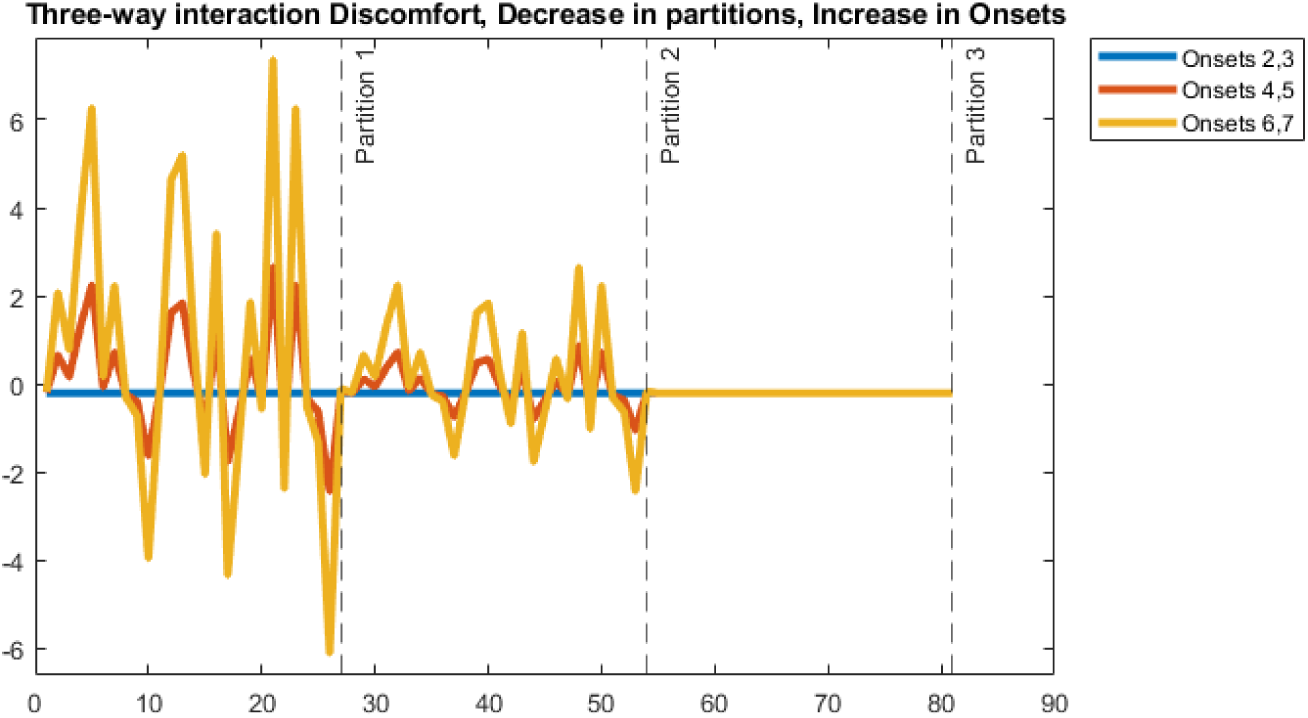
Discomfort by increase in the onsets by decrease in the partitions. X axis represents participants in each partition with dashed vertical line to mark partition boundaries, Y axis represents the scores of each participant, Onset-pair, Partition that is entered into the regressor.

### 2.8 Data Visualisation

The MUA analyses space-time maps for statistically significant correlations with each regressor and then returns a space-time map with a mask indicating the identified clusters. This approach does not visualise these significant clusters. Therefore, we create three different visualisations of the significant effects.

The first is a time-volume plot, which shows the duration of any identified clusters within the analysis window and the percentage of the volume occupied by the effects at any time-point. These are created from the returned statistics matrix from the MUA. The second is a series of topographic maps, which visualise the location of the most significant cluster on the scalp and how it changes through time. The final visualisation is the grand-average plots, which are created from the electrode in a significant cluster exhibiting the largest effect through time (i.e. with the most timepoints above the significance threshold in the cluster). This electrode is then plotted for the analysis window for each stimulus (thin, medium, thick) and PGI. For factors (not intercept), a median split is performed to visualise the differences between those high and low on a factor.

Grand averages are plotted with bootstrapped 95% confidence intervals (CI). To begin, ERPs from N participants are sampled with replacement, where N corresponds to the total number of participants. The surrogate grand average of these bootstrapped ERPs is then computed and saved for each condition. This entire process is repeated 5,000 times to create a distribution of bootstrapped (surrogate) grand-averaged across participants. Finally, confidence intervals around grand-averages are generated by calculating the 2.5% and 97.5% percentiles for each timepoint in the time series based on the generated distribution.

## 3 Results

The results are split into three main sections: 1) the effects during the DC shift period and 2) the effects during the offset period. These first two sections focus on the Discomfort factor, since the vast majority of our effects came out on this factor. The third main section presents the few results for VS and Headache that are of interest.

The first two (Discomfort-focussed) sections are further split into four sections each: Average of Onsets 2-8 (i.e. mean-intercept), Partitions (coarse time granularity), Onsets 2,3 vs 4,5 vs 6,7 (fine time granularity) and the three-way interaction (coarse vs fine time granularity). Only the three-way interaction results for the DC shift period are presented here as there were no significant or close to significant effects in the offset period.

### 3.1 DC Shift Results

#### 3.1.1 Average of Onsets 2-8

##### 3.1.1.1 Mean/Intercept Effect

In this section, we present the results for the mean/intercept effect across the collapsed average of onsets 2-8. From Table 1, we can see both the positive and negative tail yield significant clusters in the DC shift period for the mean/intercept effect. The positive cluster spans the majority of the DC shift period, whereas the negative cluster only becomes significant towards the end of the window. These positive and negative clusters are likely to reflect the same generator seen from opposite sides of the electrical dipole. Panel b in Figure 10 and Figure 11 are consistent with this interpretation. The stronger of the two is the positive going cluster, with a p value of 0.0002; this effect is large and spans the entirety of the DC shift period. Figure 10a, shows the volume of the cluster over time, the cluster occupies just over 15% of the volume at its maximum point. The negative going cluster has a p value of 0.0318, only being present at the end of the analysed window; it occupies over 25% of the volume at its maximum (Figure 11a).

**Figure 10:**
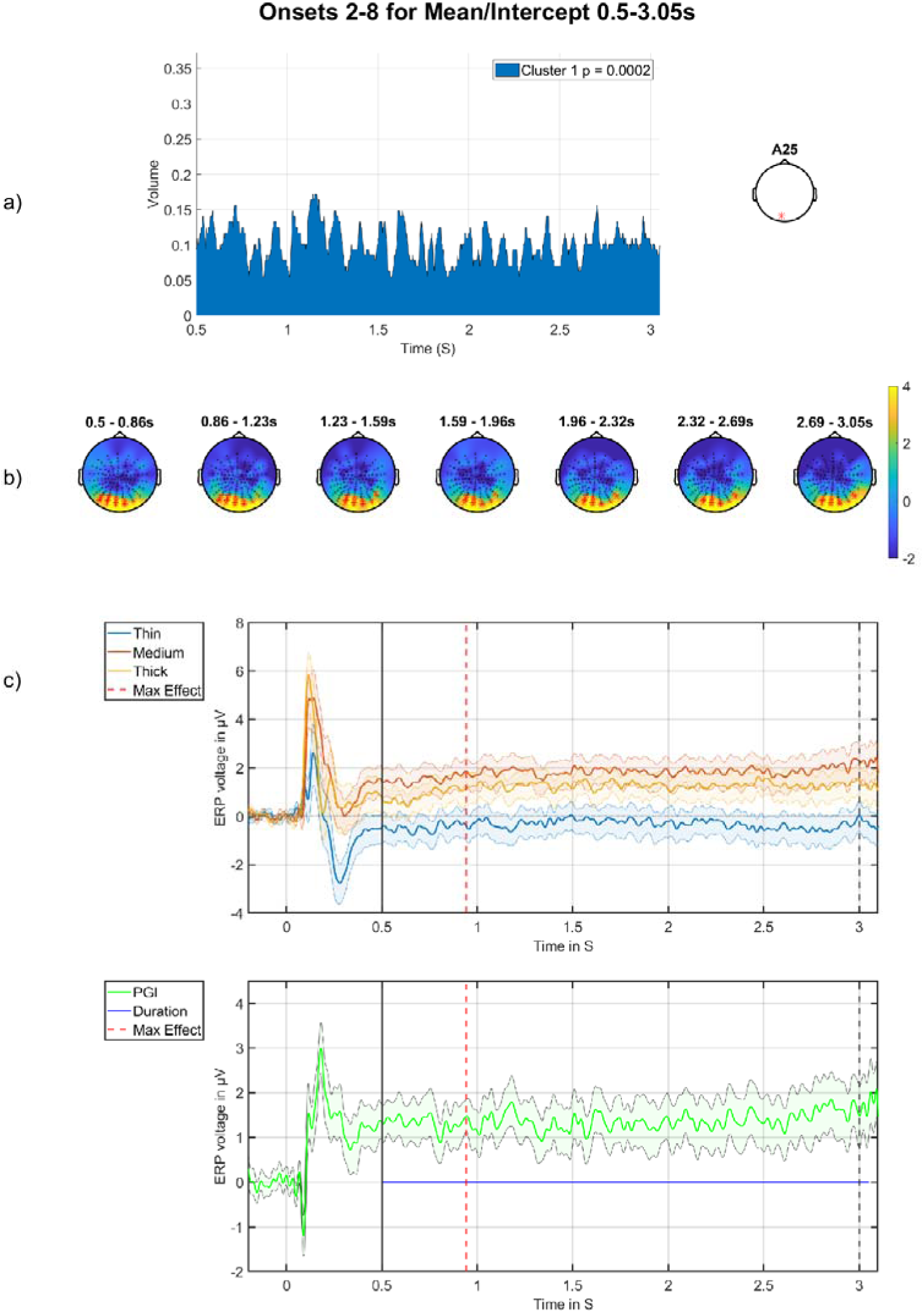
Mean/intercept onsets 2-8 positive cluster in the DC-shift period. a) cluster volume as a percentage of the entire scalp, the electrode used for plotting is displayed on the right. b) Topographic maps through time for the whole period, with red crosses indicating the significant cluster, which corresponds to the blue region in a). c) Grand-averages at electrode indicated on right in panel (a), with time of maximum effect marked with red vertical line, window start marked with a solid bl ack line, and stimulus offset marked with a dashed black line. Top are the grand-averages for thick, medium and thin, with stimulus onset marked at zero; bottom is the time-series of the PGI, with the blue horizontal line indicating the duration of the effect.

**Figure 11:**
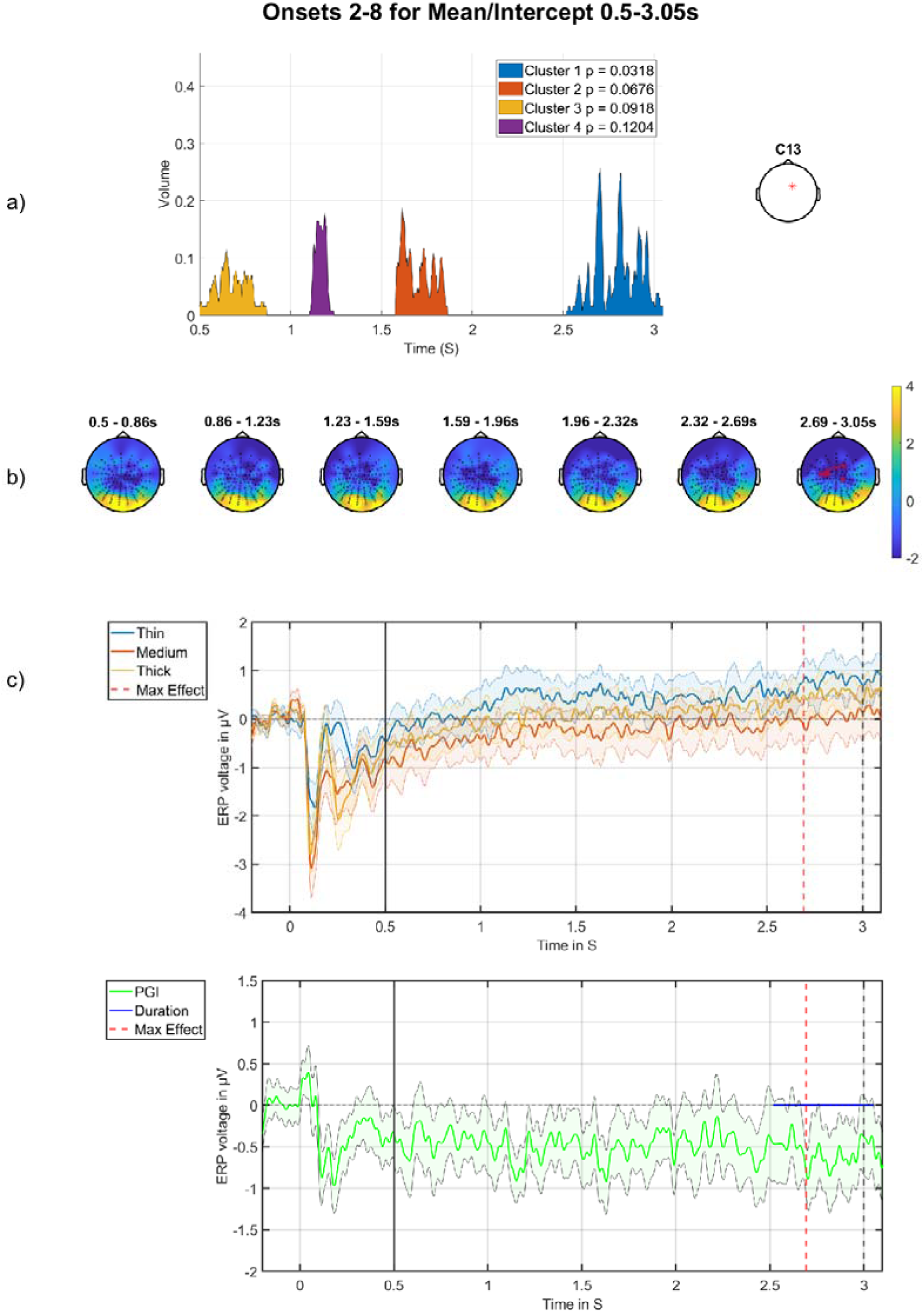
Mean/intercept onsets 2-8 negative cluster in the DC-shift period. a) cluster volume as a percentage of the entire scalp, the electrode used for plotting displayed on the right. b) Topographic maps through time for the whole period, with red crosses indicating the significant cluster, which corresponds to the blue region in a). c) Grand-averages at electrode indicated on right in panel (a), with time of maximum effect marked with red vertical line, window start marked with a solid black line, and stimulus offset marked with a dashed black line. Top are the grand-averages for thick, medium and thin, with stimulus onset marked at zero; bottom is the time-series of the PGI, with the blue horizontal line indicating the duration of the effect.

**Table 1:** MUA results for mean/intercept effect for the average of all onsets. Only results for clusters containing significant effects (FWE-corrected) or borderlin e effects smaller than a p-value of 0.1 are shown, both positive and negative tails.

| Effect | Tail (+1) | Tail (-1) |
| --- | --- | --- |
| Mean/intercept<br>0.5-3.0s | 1<sup>st</sup> cluster p-value: 0.0002 | 1<sup>st</sup> cluster p-value: 0.0318 |
|  | 1<sup>st</sup> electrode: A25 | 1<sup>st</sup> electrode: C13 |
|  | 1<sup>st</sup> peak time: 0.9414 | 1<sup>st</sup> peak time: 2.6914 |
|  | 1<sup>st</sup> r correlation effect size: 0.79 | 1<sup>st</sup> r correlation effect size: -0.65 |
|  | 1<sup>st</sup> Cohens d effect size: 2.59 | 1<sup>st</sup> Cohens d effect size: -1.69 |

The most important effect we see is a failure to return to stasis for the medium stimulus (Figure 10c), which is more positive going throughout the DC shift period compared to the thick and thin stimuli. This effect is seen across a large portion of the posterior region (Figure 10b), with the maximum effect happening at 0.9414s after stimulus onset. This suggests sustained hyper-excitation to pattern-glare, which remains, essentially, stable, with some signs of an ongoing increase, throughout a period of continuous and constant (driving) visual stimulation.

##### 3.1.1.2 Factors

There were no effects for the Discomfort factor on its own, which is a form of main-effect of our analysis (i.e. not crossed with any other variable). A non-significant effect is reported in the appendix (see section 9.2.1.1).

#### 3.1.2 Partitions

##### 3.1.2.1 Orthogonalized results

Since there were no significant effects for the pure change through time effects (i.e. basic increase or decrease effects, which would again be types of main-effect), we now present the discomfort factor effects through the coarse time granularity (partitions) of the experiment (i.e. an interaction of discomfort with time).

###### Discomfort-by-Decrease across Partitions

There was a significant negative going cluster for the discomfort by decrease effect across the partitions. The effect lasts over a second and is in the right posterior region of the scalp (Figure 12b), occupying around 10 % of the volume at its maximum point (Figure 12a), with a p value of 0.0192 (Table 2, FWE-corrected at the cluster-level). The main feature that drives this interaction is a change in the high group from partition 1 to partitions 2/ 3 (see top-row, right-side of panel c). This is a *negative*-going effect on a *decrease* across partitions. A negative decrease represents an effective increase, which is what we observe: the response *increases* from partition 1 (red) to partitions 2/3 (green/blue); i.e. over the course of the experiment. However, the signal starts negative in Partition 1 then increases towards zero in subsequent partitions. Accordingly, we view this as an habituation effect, i.e. a progression through time towards the absence of an effect: a PGI of zero (i.e. when there is no difference between Thin, Medium and Thick). In the first two panels in Figure 12c, you can see the high group (right) exhibiting this negative decrease effect, the red line indicates partition 1, green partition 2 and blue partition 3. As just discussed, for the high group, the red line (partition 1) starts as the most negative going and for the green line (partition 2), the response becomes substantially more positive, arriving at a value around zero. This effect is seen in both the PGI plot (top row) and also for the medium stimulus, suggesting that the effect is driven by a change in the brain response to the clinically-relevant (medium) stimulus, rather than to thick and thin. This in turn suggests that there is a reduction in a hyperexcitation pattern, which we are observing, at this position on the scalp (which is anterior to visual cortex), as negative-going, which starts around the middle of the DC-shift period and is only present in partition one.

**Figure 12:**
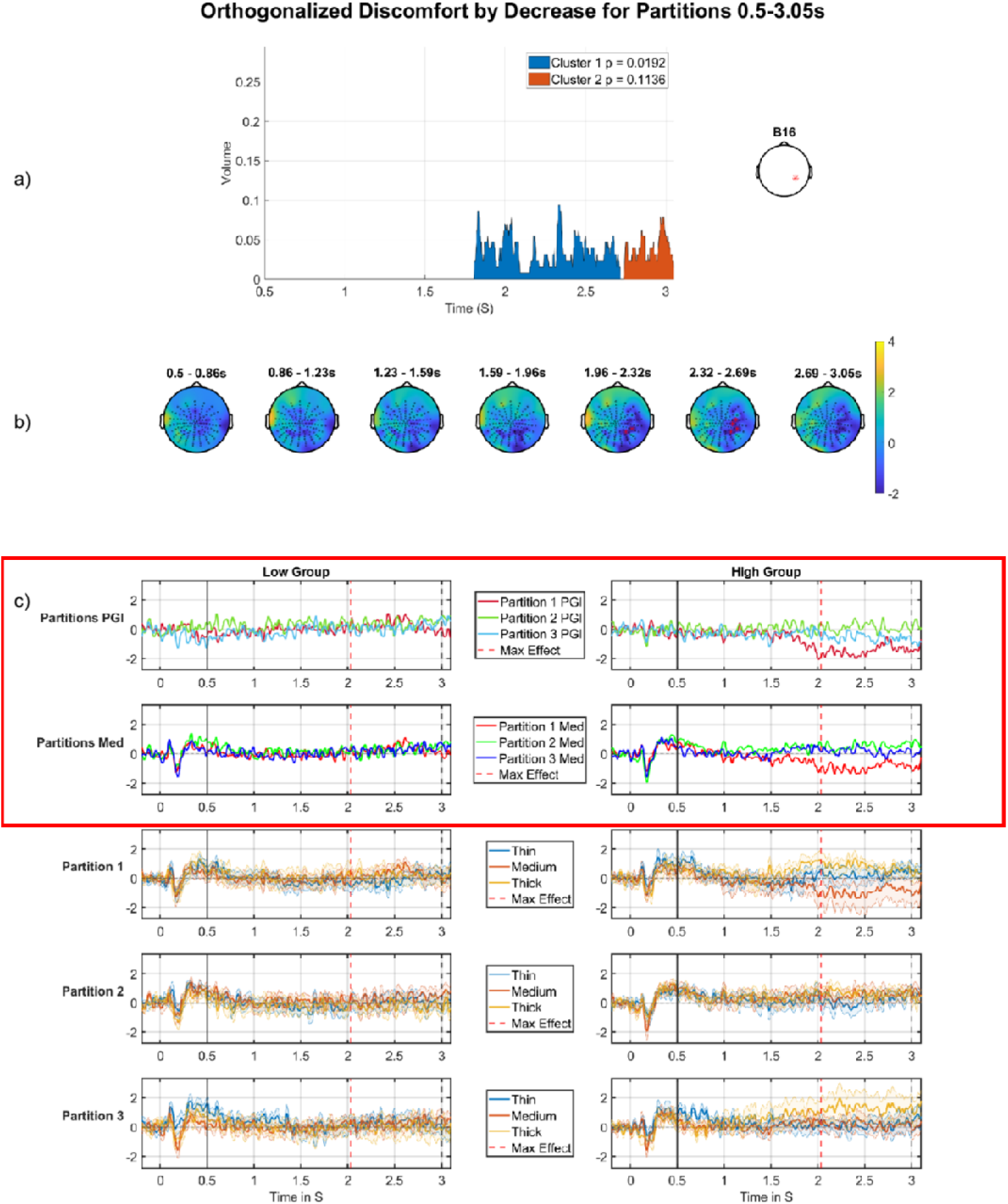
Discomfort by decrease across partitions, negative cluster, DC shift period. a) Cluster volume as a percentage of the entire scalp, the electrode used for plotting displayed on the right. b) Topographic maps through time for the whole period, with red crosses indicating the significant cluster, which corresponds to the blue region in a). c) Median split (on Discomfort) grand-averages at electrode indicated on right in panel (a), the left column of grand-averages is for the low group, right column of grand-averages is for the high group. Top is the time-series for the PGI for each partition, red (partition 1), green (partition 2), blue (partition 3) with maximum effect marked with a red vertical line, window start marked with a solid black line, and stimulus offset marked with a dashed black line; second row is the grand-averages for the medium stimulus for each partition, red (partition 1), green (partition 2), blue (partition 3); third, fourth and fifth rows present grand-averages for partitions 1, 2 and 3 (respectively), each showing thin, medium and thick. The main feature that drives this interaction is a change in the high group from partition 1 to partitions 2/ 3. This comes out as a negative effect on adecrease across partitions. A negative decrease is an increase, which is what we observe: the response increases from partition 1 to partitions 2/3. Additionally, since this increase is from negative towards zero, we can functionally view this as an habituation effect, i.e. an extreme negative-going response is tending towards what we take as stasis, which here is zero. The most important time series comparisons are the PGI and medium stimuli ERPs with the red outline, this makes the effects easy to see with the high group on both outlined rows displaying the habituation effect and the low group not showing any clear pattern.

**Table 2:** MUA results for discomfort by decrease through the partitions with orthogonalized regressor. Only results for clusters containing significant effects (FWE-corrected) or borderline effects smaller than a p-value of 0.1 are shown, both positive and negative tails.

| Effect | Tail (+1) | Tail (-1) |
| --- | --- | --- |
| Discomfort<br>by decrease<br>0.5-3.0s | No significant Cluster | 1<sup>st</sup> cluster p-value: 0.0192<br>1<sup>st</sup> electrode: B16<br>1<sup>st</sup> peak time: 2.0332<br>1<sup>st</sup> r correlation effect size: -0.48<br>1<sup>st</sup> Cohens d effect size: -1.1 |

###### Discomfort-by-Increase across Partitions

In the analysis seen in Table 3, there were two significant clusters in a similar location on the scalp with the same direction (tail -1). From the plots in Figure 13, we can say this is likely the same effect that fades briefly and then returns, thus being classified as two clusters. Setting a lower threshold for clustering (or smoothing more) could yield a single stronger cluster. The most significant cluster starts just before 1.5s after stimulus on and ends just before 2.5s. The second cluster starts at just before 2.5s after stimulus on and ends at 3s and looks like it is being cut short by the end of the analysis window (Figure 13a). The main effect we see in the grand-averages (Figure 13b) is most clearly seen in the high group (right column of Figure 13c), where we see the PGI becoming more negative (see top row of Figure 13c) as we progress through the partitions. However, there is little evidence of this pattern in the right panel of the second row of Figure 13c, i.e. for the medium stimulus, all three Partitions sit on top of each other. This in turn suggests that this effect is largely caused by changes across partitions in Thick and Thin, rather than in Medium, meaning that this effect is not a particularly clear hyper-excitation effect.

**Figure 13:**
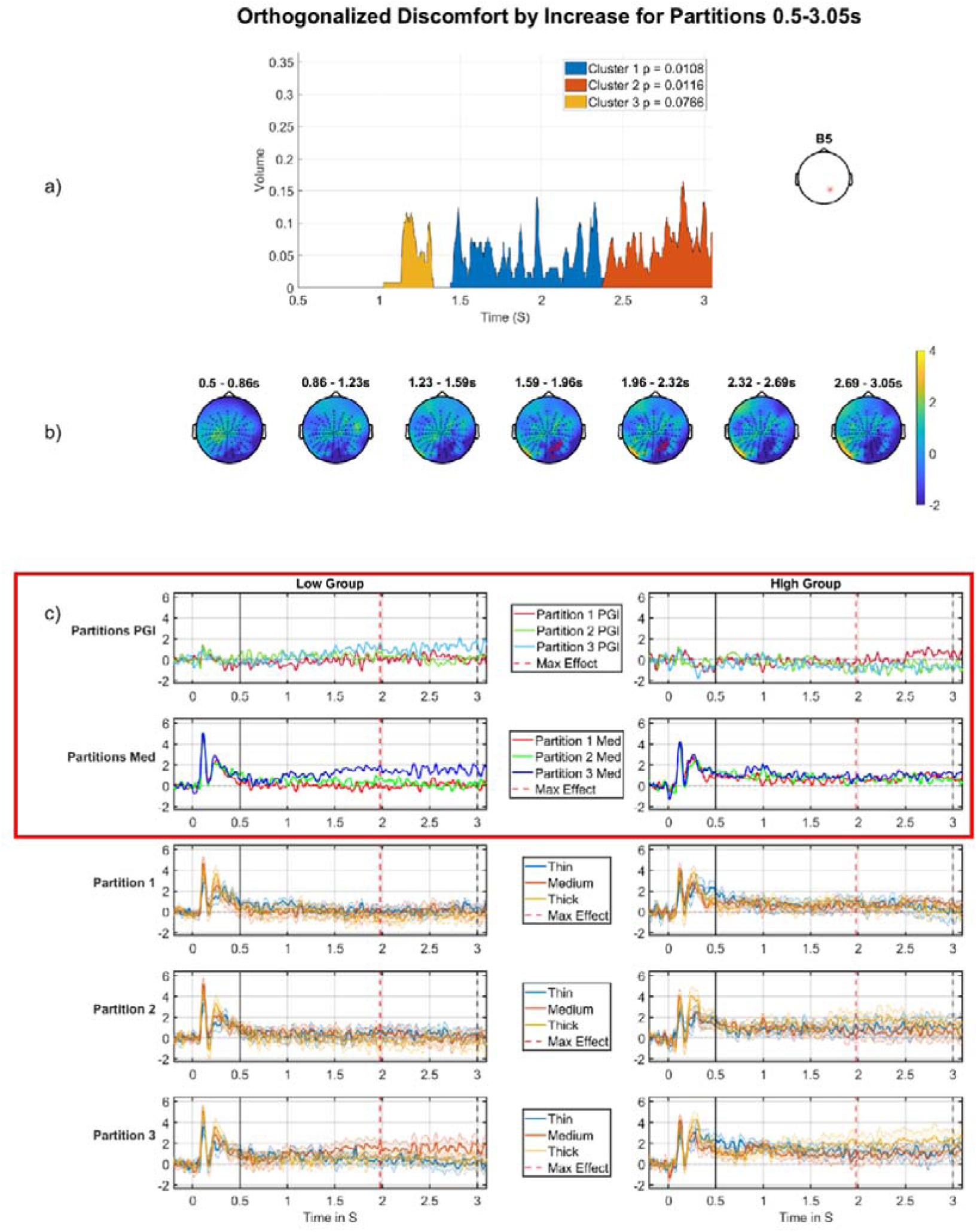
Discomfort by increase across the partitions, negative cluster, DC shift period. a) Cluster volume as a percentage of the entire scalp, the electrode used for plotting displayed on the right. b) Topographic map through time for the whole period, with red crosses indicating the significant cluster, which corresponds to the blue region in a). c) Median split (on Discomfort) grand-averages at electrode indicated on right in panel (a), the left column of grand-averages is for the low group, right column for the high group. Top is the grand-average for the PGI for each partition, red (partition 1), green (partition 2), blue (partition 3), with maximum effect marked with a red vertical line, window start marked with a solid black line, and stimulus offset marked with a dashed black line; second row is the grand-average for the medium stimulus for each partition, red (partition 1), green (partition 2), blue (partition 3); third, fourth and fifth rows present grand-averages for partitions 1, 2 and 3 (respectively), each showing thin, medium and thick; Although not directly plotted, the second significant cluster, orange in panel (a), is at an adjacent electrode to that plotted in panel (c), i.e. A31 is next to B5. Accordingly, this second cluster is effectively an extension of the one depicted here and ca n be interpreted from the plots in panel (c). The most important time series comparisons are the PGI and medium stimuli ERPs with the red outline, this makes the effects easy to see with the low group on both outlined rows displaying the sensitisation (increasing) effect and the high group not showing any clear pattern.

**Table 3:** MUA results for the discomfort by increase through the partitions with orthogonalized regressor. Only results for clusters containing significant effects (FWE-corrected) are shown, both positive and negative tails.

| Effect | Tail (+1) | Tail (-1) |
| --- | --- | --- |
| Discomfort by increase 0.5-3.0s | No significant Cluster | 1<sup>st</sup> cluster p-value: 0.0108<br>1<sup>st</sup> electrode: B5<br>1<sup>st</sup> peak time: 1.9766<br>1<sup>st</sup> r correlation effect size: -0.45<br>1<sup>st</sup> Cohens d effect size: -1.0<br>2<sup>nd</sup> cluster p-value: 0.0116 |
|  |  | 2<sup>nd</sup> electrode: A31<br>2<sup>nd</sup> peak time: 2.8809<br>2<sup>nd</sup> r correlation effect size: -0.40<br>2<sup>nd</sup> Cohens d effect size: -0.88 |

#### 3.1.3 Onsets 2,3 vs 4,5 vs 6,7

##### 3.1.3.1 Orthogonalized regressor

In this section, we present the results of discomfort through the short time granularity (onsets).

###### Discomfort-by-decrease

There was a significant cluster for the discomfort by decrease effect. At any single time-point, this cluster is relatively small in size, only reaching a maximum of 5% of the volume however, it is a strong effect, since it lasts over a second with a p value of 0.0004.

Figure 14 shows the effect, the nature of which is most clearly seen in panel c). Specifically, the top row of panel c) shows the basic effect, which starts just before 1.5 seconds and continues to the end of the segment. We see a large positive-going change from Onset-pair 4,5 to Onset-pair 6,7, for the high group (right side), but a negative-going change from Onset-pair 2,3 to Onset-pair 4,5, for the low group (left side). Additionally, in the same time-period, a similar, although weaker, effect can be observed for the medium stimulus for the high group [2nd row of panel c), right hand side], suggesting that the pattern for the high group is not just driven by changes in the response for thick and thin. Although, the low group shows little difference between mediums [2nd row of panel c), left hand side] during the period of the cluster, suggesting that the effect in the low group is driven by changes in thick and thin.

**Figure 14:**
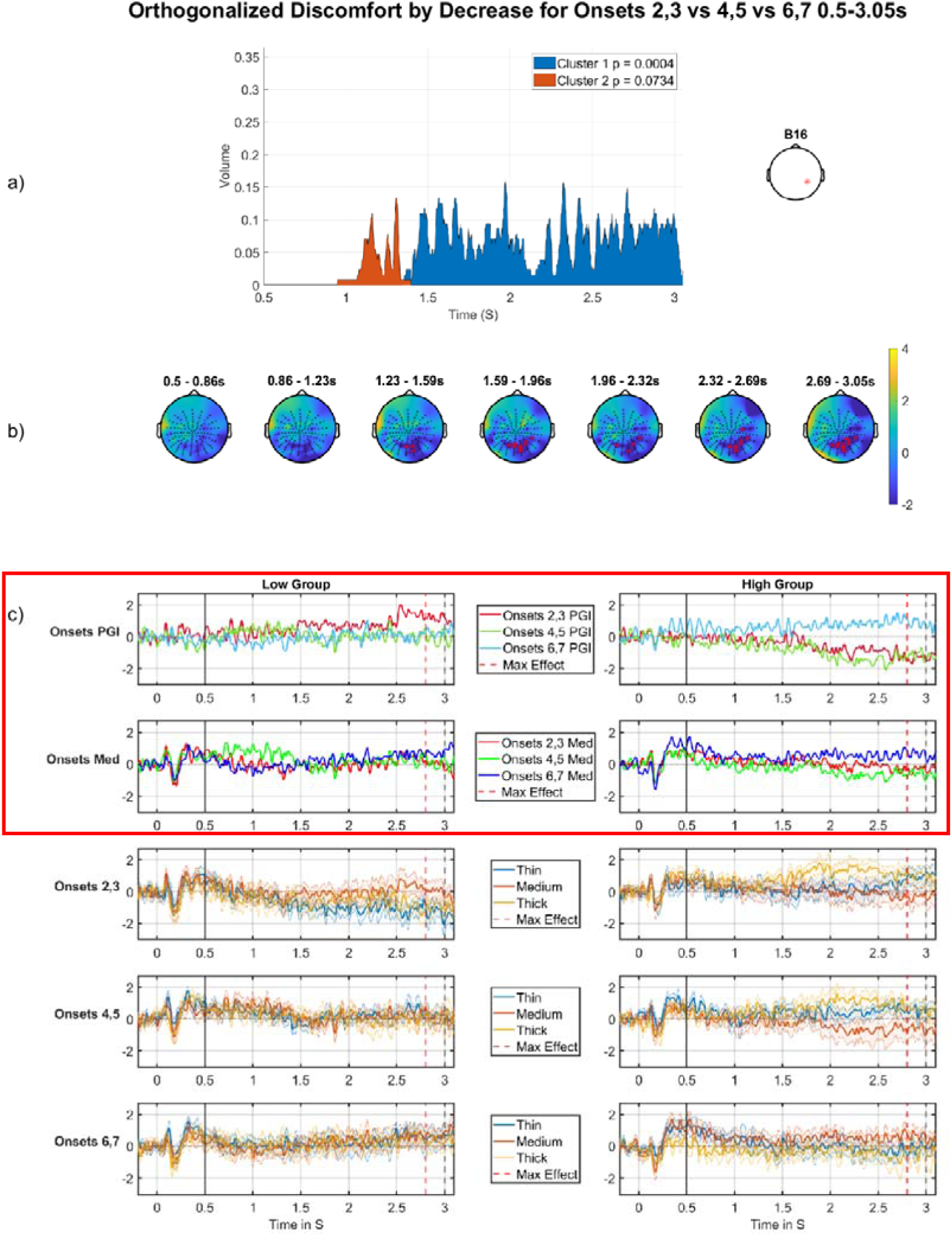
Discomfort by decrease across the onsets, negative cluster, DC shift period. a) Cluster volume as a percentage of the entire scalp, the electrode used for plotting displayed on the right. b) Topographic maps through time for the whole period, with red crosses indicating the significant cluster, which corresponds to the blue region in a). c) Median split (on Discomfort) grand-averages at electrode indicated on right in panel (a); the left column of grand-averages is for the low group, right column of grand-averages is for the high group. Top are the grand-averages for the PGI for each partition, red (partition 1), green (partition 2), blue (partition 3) with maximum effect marked with a red vertical line, window start marked with a solid black line, and stimulus offset marked with a dashed black line; second row are the grand-averages for the medium stimulus for each partition, red (partition 1), green (partition 2), blue (partition 3); third, fourth and fifth rows present grand-averages for partitions 1, 2 and 3 (respectively), each showing thin, medium and thick. The top row of panel c) shows the basic effect, which starts just before 1.5 seconds. We see a large positive-going change from onset-pair 4,5 to onset-pair 6,7 for the high group, but a negative-going change from onset-pair 2,3 to onset-pair 4,5, for the low group. Additionally, a similar, although weaker, effect can be observed for the medium stimulus for the high group [2nd row of panel c), right hand side], suggesting that the pattern for the high group is not just driven by changes in response for thick and thin. Although, the low group shows little difference between mediums [2nd r ow of panel c), left hand side], potentially indicating that the effect in the low group is driven by changes in thick and thin. The most important time series comparisons are the PGI and medium stimuli ERPs with the red outline, this makes the effects easy to see with the high group on both outlined rows displaying the increasing effect and the low group almost showing a habituation (decreasing) pattern.

Thus, the main phenomenon we observe could be interpreted as an increase in hyper-excitation in the final Onset-pair for those high on Discomfort (blue line, Figure 14c, right column, top row) and a decrease in hyperexcitation from the first Onset-pair to the second for those low on Discomfort (red line, Figure 14c, left column, top row), although, as just discussed, this latter phenomenon may not be carried by the medium stimulus (Figure 14c, second row). Notably, although we are looking at the Discomfort by *Decrease* interaction, since we are observing a negative effect, the pattern may best be interpreted as Discomfort by *Increase* ^3^, i.e. an increase through the Onsets for the high-group, and, if anything, a decrease through the Onsets for the low-group. Thus, particularly motivated by the pattern for the Medium stimulus in the high-group (right side, second row of Figure 14c), we interpret this finding “functionally” as a (differential) sensitisation effect through the onsets for those high on discomfort.

Additionally, these effects seem to be increasing as one approaches the offset of the stimulus, which occurs at three seconds. Thus, there may be a sense to which the hyperexcitation is building through the period that the stimulus is on and “driving” the visual system. That is, for those suffering discomfort during the experiment, there is sensitisation at the finest temporal scale, i.e. the three seconds that the stimulus is on for, and also sensitisation at the next temporal scale, i.e. through the sequence of onsets.

Finally, this effect comes out almost as strongly for the equivalent unorthogonalized regressor (see appendix, section 9.2.1.2). This indicates that the effect we are observing in this section is not carried by some quirk of the orthogonalization process, giving greater credence to the effect’s reliability.

#### 3.1.4 Three-way Interaction

##### Discomfort by increase across the partitions and decrease across the onsets

Although not quite significant, the three-way interaction is qualitatively present in the grand-averages (Figure 15c). The first row (partition 1) shows a striking sensitisation effect for the high group (right side), with a substantially higher response in the final Onset-pair (6,7). This pattern is absent, and potentially reversed into an habituation pattern for the corresponding low group grand-averages (left side of first row). Additionally, this sensitisation across Onsets for high and weak habituation for low in partition 1 is also present when we plot the medium alone (2nd row), suggesting the elevated Onset-pair 6,7 effect for the high group truly reflects hyperexcitation. In contrast, the remaining 4 rows of panel c), which correspond to partitions 2 and 3, exhibit no, or certainly much weaker, patterns of change through the onsets.

**Figure 15:**
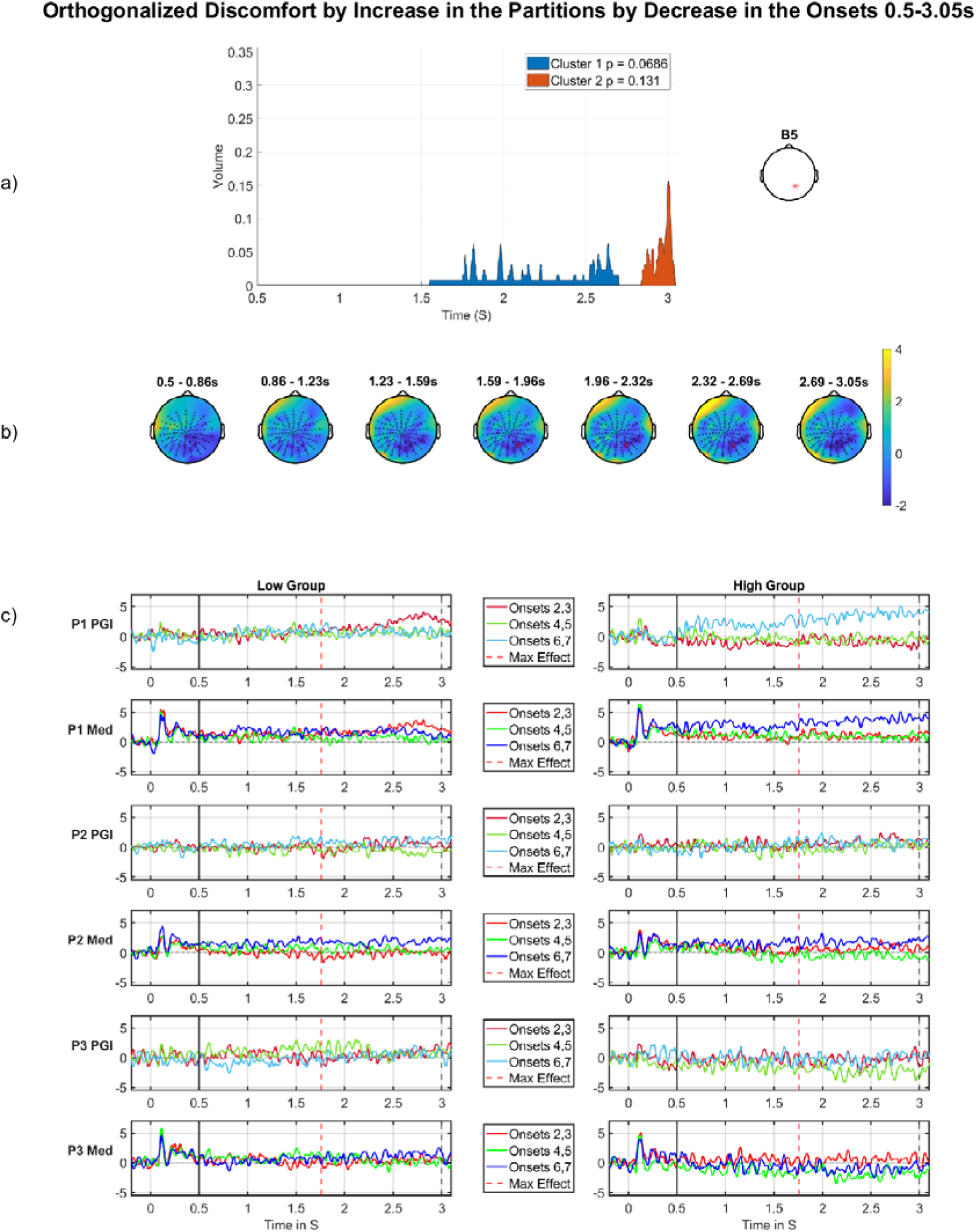
Discomfort by increase (through the partitions) by decrease (through the onsets), DC shift period. a) Cluster volume as a percentage of the entire scalp, the electrode used for plotting displayed on the right. b) Topographic map through time for the whole period, wit h red crosses indicating the significant cluster, which corresponds to the blue region in a). c) Median split (on Discomfort) grand-averages at electrode indicated on right in panel (a); the left panel of grand-averages is for the low group, right panel of grand-averages is for the high group. Top row (partition 1), third row (partition 2) and fifth row (partition 3) show grand-averages for the PGI for each onset, red (onsets 2,3), green (onsets 4,5), blue (onsets 6,7) with maximum effect marked with a red vertical line, window start marked with a solid black line, and stimulus offset marked with a dashed black line. Second row (partition 1), fourth row (partition 2) and sixth row (partition 3) are the grand-averages for just the medium stimulus for each onset, red (onsets 2,3), green (onsets 4,5), blue (onsets 6,7). Panel c) shows what underlies this effect. The first row (partition 1) shows a striking sensitisation effect for the high group (right side), with a substantially higher response in the final Onset-pair (6,7). This pattern is absent, and potentially reversed into an decrease pattern for the corresponding low group grand-averages (left side of first row). Additionally, this increase across Onsets for high and weak decrease for low in partition 1 is also present when we plot the medium alone (2nd row), suggesting the elevated Onset-pair 6,7 effect for the h igh group truly reflects hyperexcitation. In contrast, the remaining 4 rows of panel c), which correspond to partitions 2 and 3, exhibit no or certainly much weaker patterns of change through the onsets.

Thus, the key phenomena are in Partition 1, with the high group showing a clear sensitisation effect, including very substantial hyper-excitation in the final Onset-pair. This suggests that the process of repeating the aggravating stimulus (over a short timeframe of 10s of seconds) increases the brain’s sensitivity to that stimulus.

When we look through the partitions, we see an habituation effect: the differential sensitisation effect that we observe in partition 1 (sensitisation through onsets for high group and not for low group) reduces from then on, effectively being absent in partitions 2 and 3. Thus, this effect suggests a selective hypersensitivity when those susceptible to pattern glare stimuli start the experiment, which their brains successfully quench through the course of the experiment.

Though not significant, this effect does occur in a region we have seen the discomfort effects for previous significant clusters. In particular, this three-way interaction is a decomposition of the two-way interaction we observed in Figure 14: it happens at the same position on the scalp, in a similar time interval.

This three-way interaction effect occupies only 5% of the volume at its maximum point but does last a significant portion of the analysis period. With regards to this effect failing to be significant, it is important to note that adding an extra factor to an interaction (going from two-way to three-way) will reduce statistical power, since the data needs to be distributed across more bins. Additionally, we are performing statistical inference in a large volume (the entire DC shift period), which reduces the statistical power of the FWE-correction we perform with Fieldtrip.

### 3.2 Offset Results

Now we present stimulus offset effects. These effects are presented the same way as the DC shift results: mean/intercept effects, followed by factor effects, pure change through time effects and then interactions between time and factors.

#### 3.2.1 Average of Onsets 2-8

##### 3.2.1.1 Mean/intercept Effect

For the mean/intercept in the offset period, we see negative going clusters across all windows of analysis and a positive going effect for the first window of analysis. The full set of these results, across all windows analysed, are presented in Table 11, in the appendix (see section 9.2.2.1).

From Table 11, we can see multiple significant clusters. First, the negative going cluster in the large window (3.09 – 3.99s), is large and spans much of the window, having a p-value of 0.0004 (FWE-corrected at the cluster-level). Figure 16a shows the extent of the cluster (blue), which occupies around 10% of the scalp’s volume at its maximum point. Figure 16b shows the location of the cluster, which sits over the centre of the occipital lobe. The cluster does not move substantially spatially over the course of the significant period. Figure 16c shows the grand-averages from the strongest electrode in the cluster: divided into thick, medium and thin (top) and PGI (bottom). The duration of the effect is shown in the bottom plot (blue horizontal solid line) and its point of maximum effect (red vertical dashed line) is shown in both panels. The effect suggests a differentially large (in a negative-going direction) return to stasis for the medium stimulus, which is consistent with Medium being elevated relative to Thin and, to some extent, Thick during the DC-shift period; see Figure 10. That is, Medium has to move further in a negative-going direction to return to stasis, than Thin and, to some extent, Thick.

**Figure 16:**
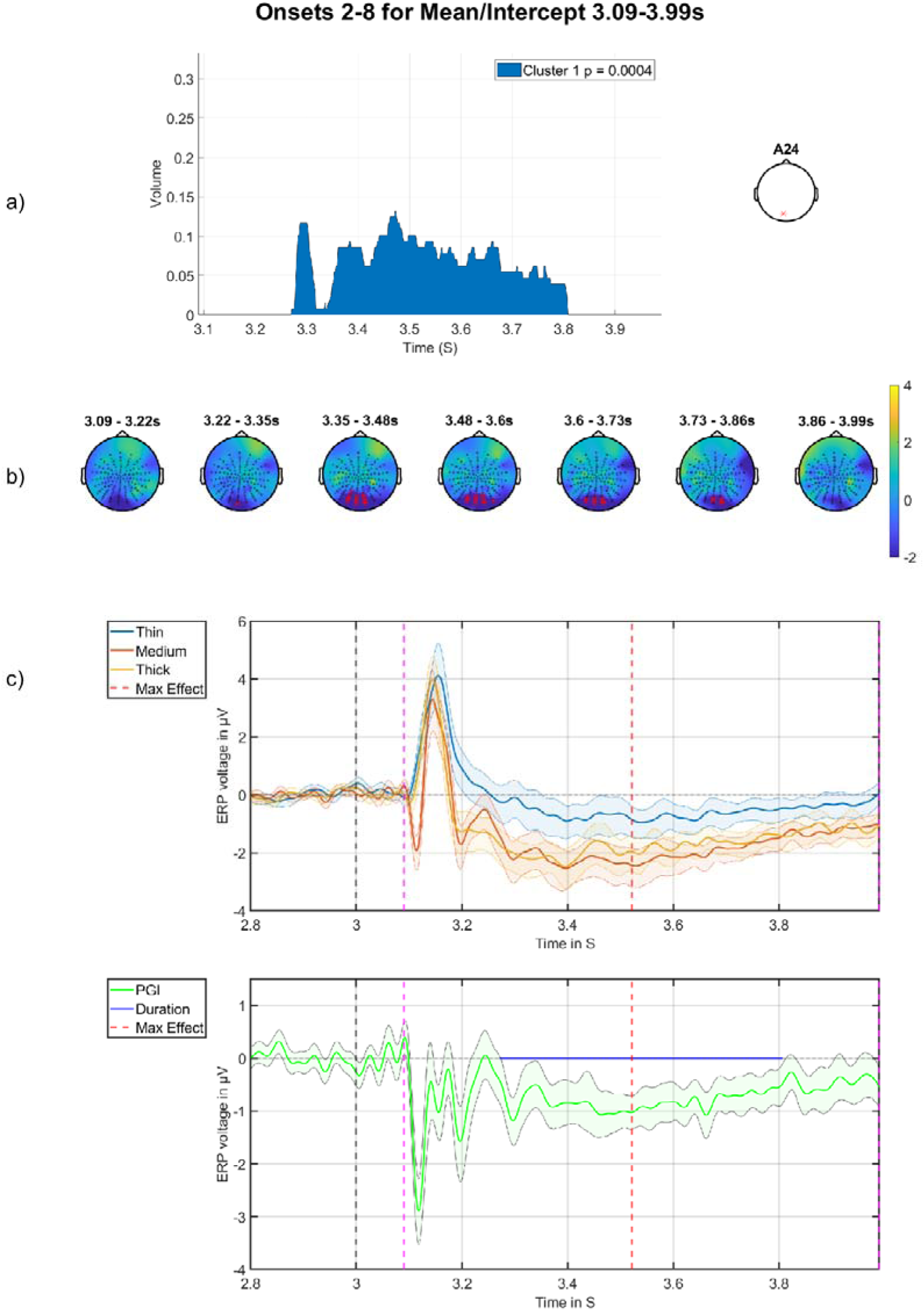
Mean/intercept of offset. a) Cluster volume as a percentage of the entire scalp, the electrode used for plotting displayed on the right. B) Topographic maps through time for the whole period, with red crosses indicating the significant cluster, which corresponds to the blue region in a). c) grand-averages at electrode indicated on right in panel (a), with time of maximum effect marked with red vertical dashed line, stimulus offset marked with black dashed line and window of analysis marked with pink dashed lines. Top are the grand-averages for thick, medium and thin, with stimulus offset marked at three seconds with a black dashed line; bottom is the grand-average of the PGI, with the blue horizontal line indicating the period of statistical significance. A highly significant negative-going cluster can be observed that sits over visual cortex and is extended in time, suggesting a differentially extreme (in a negative-going direction) return to stasis for the medium stimulus.

The most significant cluster in the first window (3.09 – 3.18s) shows an effect that is negative-going with a p value of 0.0076 (FWE-corrected at the cluster-level). This cluster is shown in Figure 17b (indicated with red crosses), where it sits over the occipital lobe and is short in time. This cluster is the first effect after the stimulus is turned off and the medium stimulus shows a qualitative difference from the two control stimuli, with medium exhibiting a negative deflection that is wholly absent for thick and thin. We have called this effect the *PGI Offset N120*. In Figure 17a, the cluster’s volume over time can be seen, it is very short, only lasting for around 30ms. This shortness in time could be why this effect was not significant in the full offset window analysis. Figure 17c shows the grand-averages for this time plotted at A26, the top grand-averages shows all three stimuli and at around 3.1s there is a sharp negativity for only the medium stimulus; this feature is even more apparent in the PGI grand-average (bottom). This short and sharp medium-specific negativity may suggest an inhibitory mechanism that is specific to the clinically relevant stimulus.

**Figure 17:**
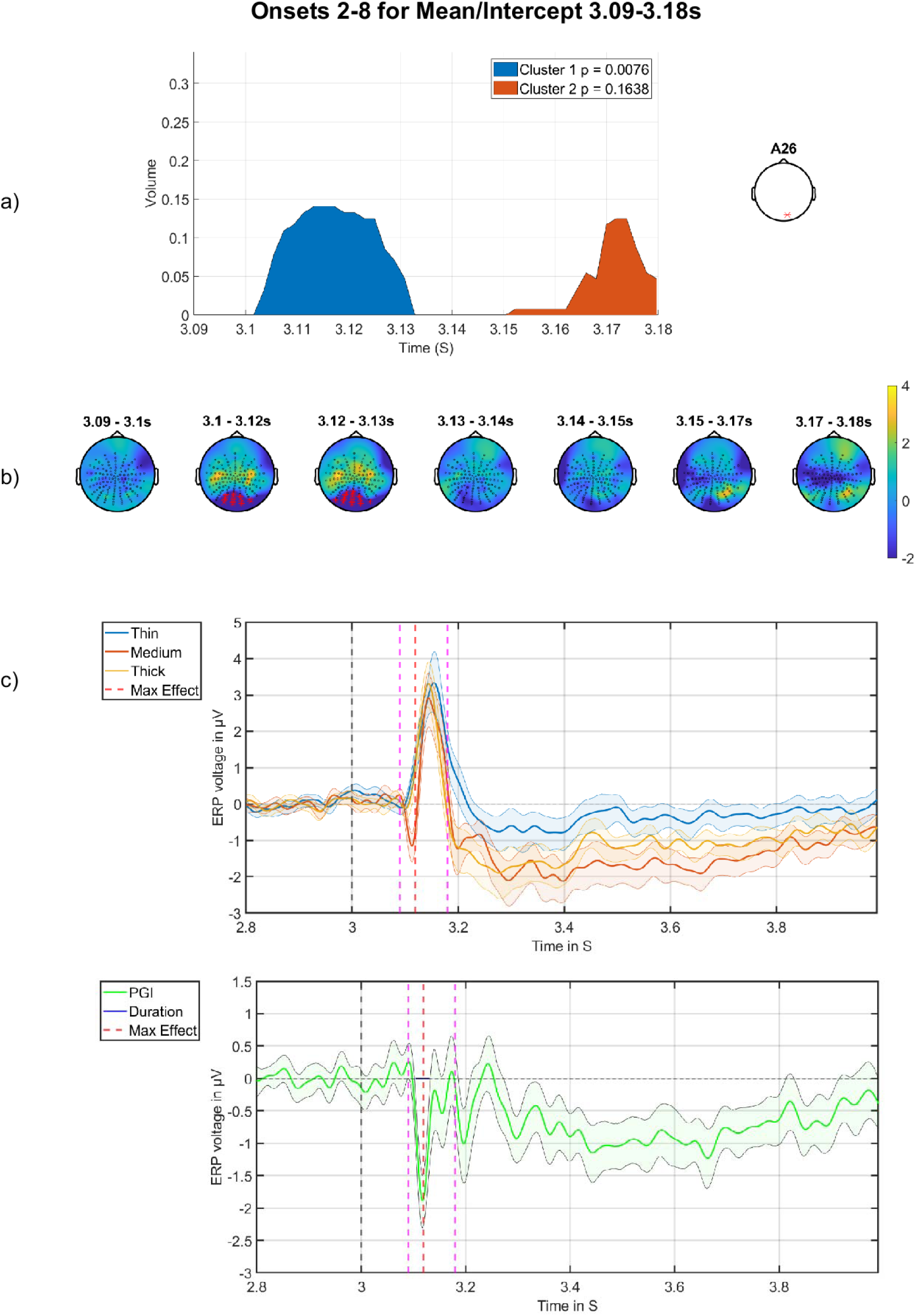
Negative-going mean/intercept effect in first analysis window. a) Cluster volume as a percentage of the entire scalp, with the electrode used for plotting displayed on the right. b) Topographic maps through time for the whole period, with red crosses indicating the significant cluster, which corresponds to the blue region in a). c) grand-averages at electrode indicated on right in panel (a), with time of maximum effect marked with red vertical dashed line, stimulus offset marked wi th black dashed line and window of analysis marked with pink dashed lines. Top are the grand-averages for thick, medium and thin, with stimulus offset at three seconds with a black dashed line; bottom is the grand-average of the PGI, with the blue horizontal line indicating the period of statistical significance. A short, but very high PGI, negative-going cluster can be observed that sits over visual cortex. Strikingly, we observe a qualitative difference between stimulus responses, with medium exhibiting a negative deflection, which is completely absent for thick and thin.

There is also a positive cluster for the first window. This cluster, shown in Figure 18, looks to be the opposite side of the dipole observed in the previous PGI offset N120 effect. This is particularly clear from Figure 18b. That is, in Figure 18b,c a short positive-going PGI cluster can be observed that is relatively central on the scalp. Again, we observe a qualitative difference between stimulus responses, with medium exhibiting a positive deflection around the red dashed vertical line, which is completely absent for thick and thin. The temporal coincidence of this effect to the negative deflection seen in Figure 17 suggests that this feature is the positive-going side of the same electrical dipole that generates the negative-going effect of Figure 17.

**Figure 18:**
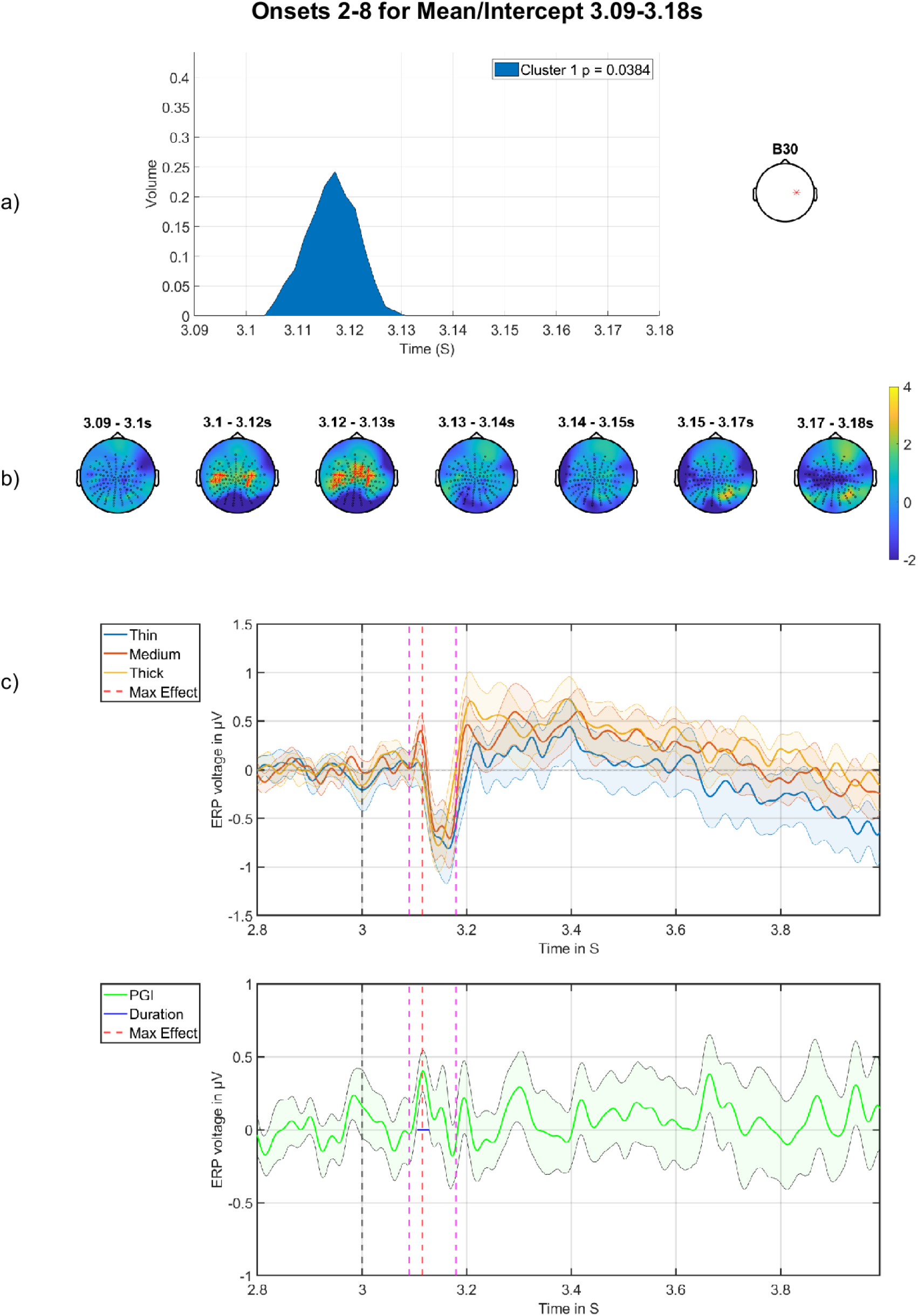
Positive-going mean/intercept effect in first analysis window. a) Cluster volume as a percentage of the entire scalp, with the electrode used for plotting displayed on the right. b) Topographic maps through time for the whole period, with red crosses indicating the significant cluster, which corresponds to the blue region in a). c) grand-averages at electrode indicated on right in panel (a), with time of maximum effect marked with red vertical dashed line, stimulus offset marked with black dashed line and window of analysis marked with pink dashed lines. Top are the grand-averages for thick, medium and thin, wit h stimulus offset at three seconds; bottom is the grand-average of the PGI, with blue horizontal line indicating the period of statistical significance. A short positive-going PGI cluster can be observed that is relatively central on the scalp. Again, we observe a qualitative difference between stimulus responses, with medium exhibiting a positive deflection around the red dashed vertical line, which is completely absent for thick and thin. The temporal coincidence of this effect to the negative deflection seen in figure 17 suggests that this feature is the positive-going side of the same electrical dipole that generates the negative-going effect of figure 17.

There are two further significant clusters, which are seen in the second and third windows of analysis for the mean/intercept. However, these are parts of the effect observed in Figure 16 split across two analysis windows. Further information can be found in the appendix (see Figure 28 and Figure 29 and the discussion in section 9.2.2.1).

##### 3.2.1.2 Orthogonalized factor results

###### Discomfort Factor

Now, we present the results for discomfort on the average of onsets 2-8. There was a close to significant effect for the Visual Stress factor which can be found in Figure 36 and Table 17.

In Table 6, only the significant analysis for the discomfort factor is presented, both positive and negative tails. Amongst the factor effects, only the positive tail for discomfort surpasses the second level statistic threshold for significance. This effect for discomfort is a strong positive effect with a p value of 0.018 (FWE-corrected at the cluster-level). It falls in the first window of analysis and could be related to the effect on the mean/intercept in a similar time-window (Figure 17); compare the blue region in Figure 17a and in Figure 19a.

**Figure 19:**
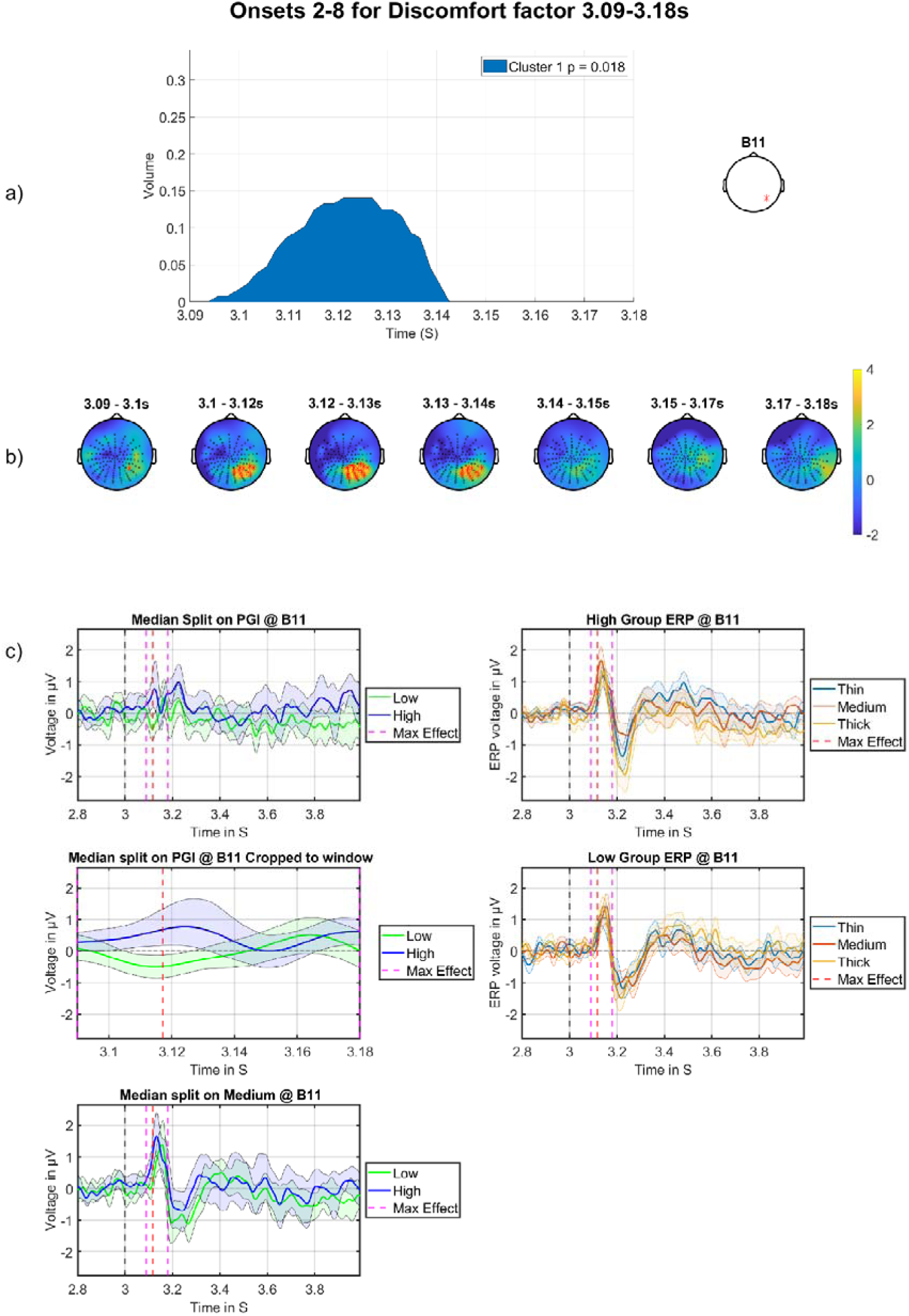
Offset effect for the discomfort factor on the average of the onsets. a) Cluster volume as a percentage of the entire scalp. The electrode used for plotting is displayed on the right. b) Topographic maps through time for the whole period, with red crosses indicating the significant cluster, which corresponds to the blue region in a). c) grand-averages at electrode indicated on right in panel (a), with maximum effect marked with the red dashed vertical line, stimulus offset marked with a black dashed line and analysis window indicated with dashed pink lines. Top left is the high vs low discomfort group for the PGI; middle left are the grand-averages for high vs low discomfort group for the PGI with window showing only the period of analysis; bottom left are the high vs low group for the medium stimulus; top right are the grand-average s for the high group showing thick, medium and thin; bottom right are the grand-averages for the low group showing thick, medium and thin. We observe a differential latency and amplitude change for the medium stimulus, such that the high group has a higher (positive-going) and earlier peak, accompanied by a shallower following negativity (this negativity is lowest around 3.25 s). This suggests a higher amplitude accelerated response associated with hyper-excitation.

**Table 4:** MUA results for the discomfort by decrease through the onsets with orthogonalized regressor. Only results for clusters containing significant effects (FWE-corrected) or borderline effects smaller than a p-value of 0.1 are shown, both positive and negative tails.

| Effect | Tail (+1) | Tail (-1) |
| --- | --- | --- |
| Discomfort<br>by decrease<br>0.5-3.0s | No significant Cluster | 1<sup>st</sup> cluster p-value: 0.0004<br>1<sup>st</sup> electrode: B16<br>1<sup>st</sup> peak time: 2.8037<br>1<sup>st</sup> r correlation effect size: -0.53<br>1<sup>st</sup> Cohens d effect size: -1.24 |

**Table 5:** MUA results for the discomfort by increase (through the partitions) by decrease (through the onsets). Only results for analysis windows containing si gnificant effects (FWE-corrected) are shown, both positive and negative tails.

| Effect | Tail (+1) | Tail (-1) |
| --- | --- | --- |
| Discomfort<br>by increase<br>in<br>partitions<br>and<br>decrease in<br>onsets 0.5-3.0s | No significant Cluster | 1<sup>st</sup> cluster p-value: 0.0686<br>1<sup>st</sup> electrode: B5<br>1<sup>st</sup> peak time: 1.7578<br>1<sup>st</sup> r correlation effect size: -0.28<br>1<sup>st</sup> Cohens d effect size: -0.58 |

**Table 6:**
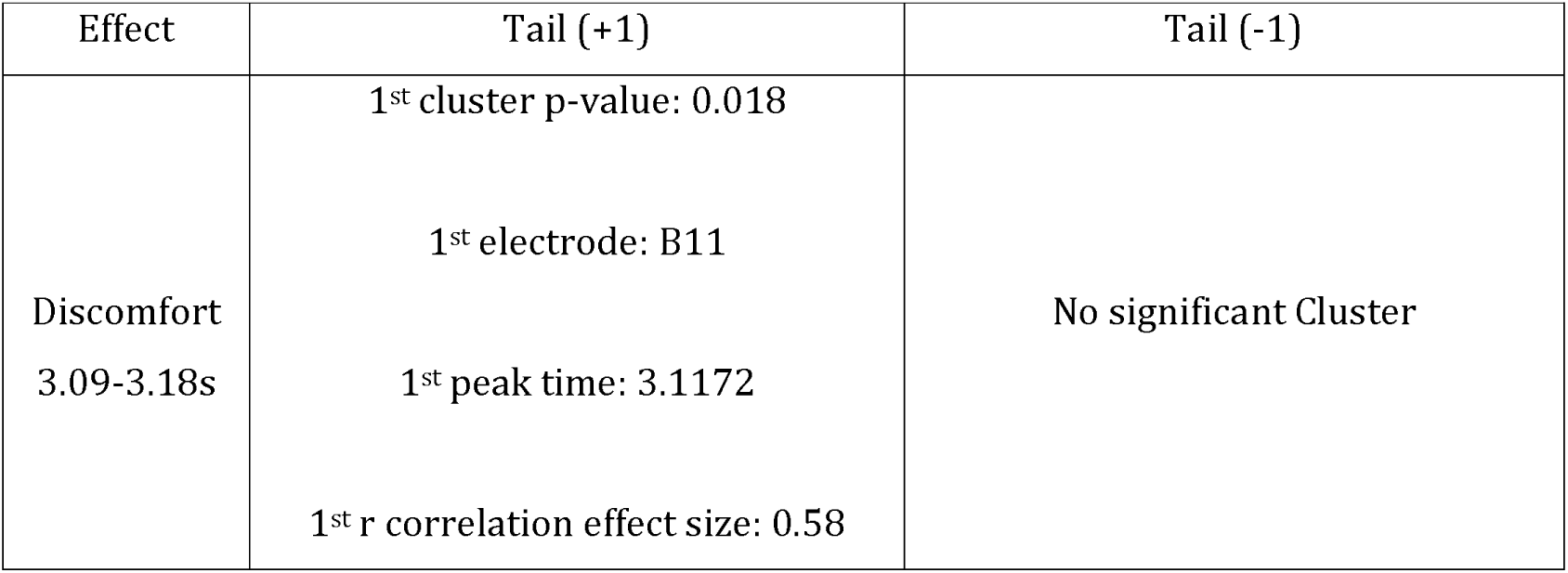
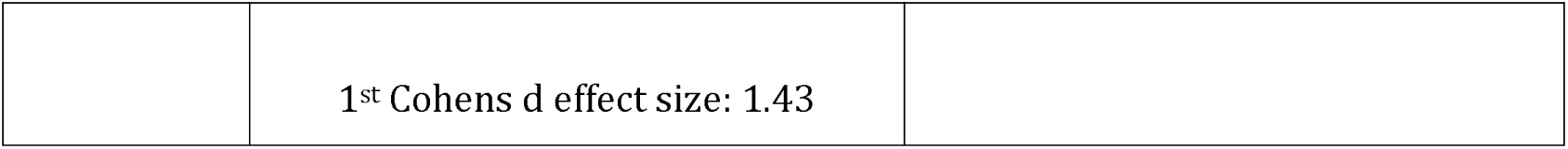
MUA results for discomfort on the average of onsets 2-8 orthogonalized regressor. Only results for clusters containing significant effects (FWE-corrected) or borderline effects smaller than a p-value of 0.1 are shown, both positive and negative tails.

**Table 7:** MUA results for the discomfort factor by decrease across the partitions for the time period 3.09 – 3.18s. Only results for clusters containing significant effects (FWE-corrected) or borderline effects smaller than a p-value of 0.1 are shown, both positive and negative tails.

| Effect | Tail (+1) | Tail (-1) |
| --- | --- | --- |
| Discomfort<br>by decrease<br>3.09-3.18s | 1<sup>st</sup> cluster p-value: 0.0262<br>1<sup>st</sup> electrode: B14<br>1<sup>st</sup> peak time: 3.1250<br>1<sup>st</sup> r correlation effect size: 0.37 | No significant Cluster |
|  | 1<sup>st</sup> Cohens d effect size: 0.79 |  |

Figure 19 shows the discomfort cluster. Figure 19a shows this cluster’s volume over time; the cluster lasts 40ms and occupies just under 15% of the whole volume at its maximum point. In Figure 19b, the location of the cluster over time is shown; it occupies a right posterior region on the scalp, which does not substantially overlap with the cluster for the mean/intercept PGI offset N120 effect. Figure 19c shows the grand-averages from the cluster’s strongest electrode: B11. There is a clear difference in the PGI for the median split on discomfort; see Figure 19c’s top two plots on the left, with the top plot showing the period 2.8s-4s, which indicates the difference of the two groups over the whole offset period. The second plot shows the analysis window (3.09 – 3.18s), in which a clearer difference can be seen, with the High group having a more positive going grand-average than the low group. The bottom plot on the left of Figure 19c’s grand-average plots show the median split on the medium stimulus, from which we can see the effect is primarily driven by the medium stimulus. This is consistent with the right column of grand-averages, which show that the high group has a higher (positive-going) and earlier peak, which is accompanied by a shallower following negativity (this negativity is lowest around 3.25 s). This suggests a higher amplitude accelerated electrophysiological response associated with hyper-excitation.

Figure 30d (see section 9.2.2.2 in the appendices) shows the grand-averages of the same median split on discomfort at the most significant electrode from the PGI offset N120 effect. This is included to determine if there was any effect of the discomfort factor on the N120. Although not significant, a difference between the two groups is shown, with the high group having a shallower negativity than the low group, which is the opposite pattern to what would be naturally expected of a hyper excitation explanation.

#### 3.2.2 Partitions

Next, we present the results for the MUA on the coarse time granularity (partitions) on the offset period. The same temporal windows are used as in the averaged onsets analysis.

##### 3.2.2.1 Pure Change through Time

There was a significant effect for the pure change through time in the offset period however, this effect was only present on fringe electrodes, which may be noisy, so it is included in the appendix (see section 9.2.2.3, Figure 31 and Table 12).

##### 3.2.2.2 Orthogonalized factor results

###### Discomfort-by-decrease

Next, we present the MUA results for discomfort by decrease (across the partitions) effect. For this analysis, only effects for the discomfort factor crossed the FWE-corrected significance threshold.

The effect seen for discomfort by decrease on the partitions is a relatively strong effect with a p value of 0.0262 (FWE-corrected at the cluster-level): it lasts most of the period of analysis and at its maximum point occupies just over 10% of the volume; see Figure 20a. This effect somewhat overlaps with the pure factors discomfort effect, both spatially and temporally, see Figure 19. From the grand-averages in Figure 20c, a difference in the two groups can be seen. Firstly, in the top row of grand-averages, the high group’s PGI through the partitions (right hand side), shows an habituation effect, where the first partition (red) is seen as the most positive going then partitions 2 (green) and then 3 (blue). In comparison, the grand-averages in the top row suggest that the low group does not have an habituation pattern; see left hand side. For the second row of the grand-averages, a less pronounced pattern for both the high and low groups can be seen suggesting this effect is at least partially driven by the two control stimuli.

**Figure 20:**
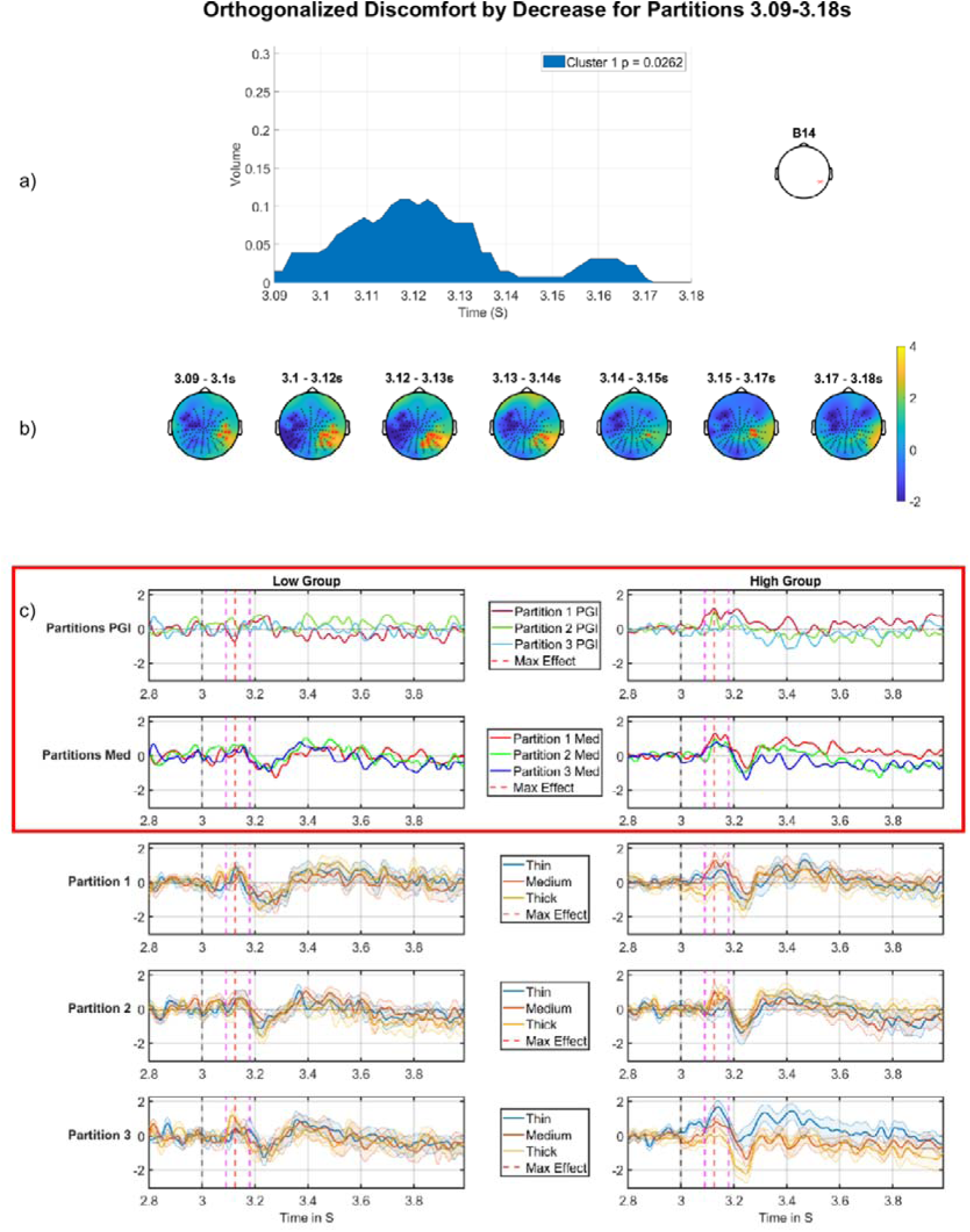
Discomfort by decrease regressor across the partitions on the offset period. a) Cluster volume as a percentage of the entire scalp; the electrode used for plotting displayed on the right. b) Topographic maps through time for the whole period, with red crosses indicating the significant cluster, which corresponds to the blue region in a). c) Median split (on Discomfort) grand-averages at electrode indicated on right in panel (a). The left panel of grand-averages is for the low group, right panel of grand-averages is for the high group. Top are the grand-averages for the PGI for each partition, red (partition 1), green (partition 2), blue (partition 3), with maximum effect marked with a red vertical dashed line, stimulus offset marked with a black dashed line and analysis window marked with pink dashed lines; second row are the grand-averages for the medium stimulus for each partition, red (partition 1), green (partition 2), blue (partition 3); third, fourth and fifth rows present grand-averages for partitions 1, 2 and 3 (respectively), each showing thin, medium and thick. There is a clear tendency throughout the time-period shown for partition one medium to be highest fo r the high group (right side), with much less evidence of this pattern for the low group (left side). This does then suggest a pattern of hyper excitation early in the experiment that is only strongly present for the high group and which reduces through the course of the experiment. The most important time series comparisons are the PGI and medium stimuli ERPs with the red outline, this makes the effects easy to see with the high group on both outlined rows displaying the habituation effect and the low group not showing any clear pattern.

We see another effect on a discomfort related contrast in the same right posterior region of the scalp. We have seen a number of such effects in the same region of the scalp in both the DC-shift time window and offset-transients time windows. The time point of maximum effect (see vertical red dashed line) shows a convincing pattern for the pattern-glare index (see top row of Figure 20). However, the pattern at that time point is not quite so clear cut when just looking at the medium stimulus (see second row of Figure 20). In fact, the pattern for medium looks most compelling from about 3.3 sec on to the end of the analysis segment.

Nonetheless, we see a pretty convincing habituation effect through the partitions for those high on the discomfort factor, which is not present and indeed for some time segments reversed, for the low group. As we see frequently, the pattern for the high group in partition 1 to a large extent drives the effect, suggesting a hyper-excitation response for those susceptible to discomfort when they are first presented the aggravating stimulus (the medium during partition 1), which relatively swiftly habituates during the course of the experiment.

There were no significant effects for factors by increase on the partitions with any factors.

#### 3.2.3 Onsets 2,3 vs 4,5 vs 6,7

The only significant cluster identified for this analysis is reported in the appendix (see 9.2.2.4, Table 13 and Figure 32). Although it is potentially of interest, we have presented this effect in the appendix because it is difficult to reconcile with our main findings. This is unless it represents a polarity reversal arising from observing the underlying electrical dipole from opposite sides. In which case, the pattern on the other side of the dipole would be consistent with our main findings. We leave this for further research.

### 3.3 DC Shift Offset Correlation

To link our findings in the DC shift period to the offset period, we correlated the highest amplitude in the DC shift period to the deepest negativity seen in the offset period (3.09 – 3.18s), for the electrode that was most significant for both of the mean/intercept analyses. For the DC shift period this was electrode A25 and for the offset this was electrode A26 (note that these electrodes are next to each other on the scalp with A26 being slightly more posterior). Simple correlation between these two points for each participant on the average of onsets 2-8 produced a correlation of 0.56 (p = 0.0002) for the PGI and 0.61 (p = 0.0001) for the medium stimulus.

**Figure 21:**
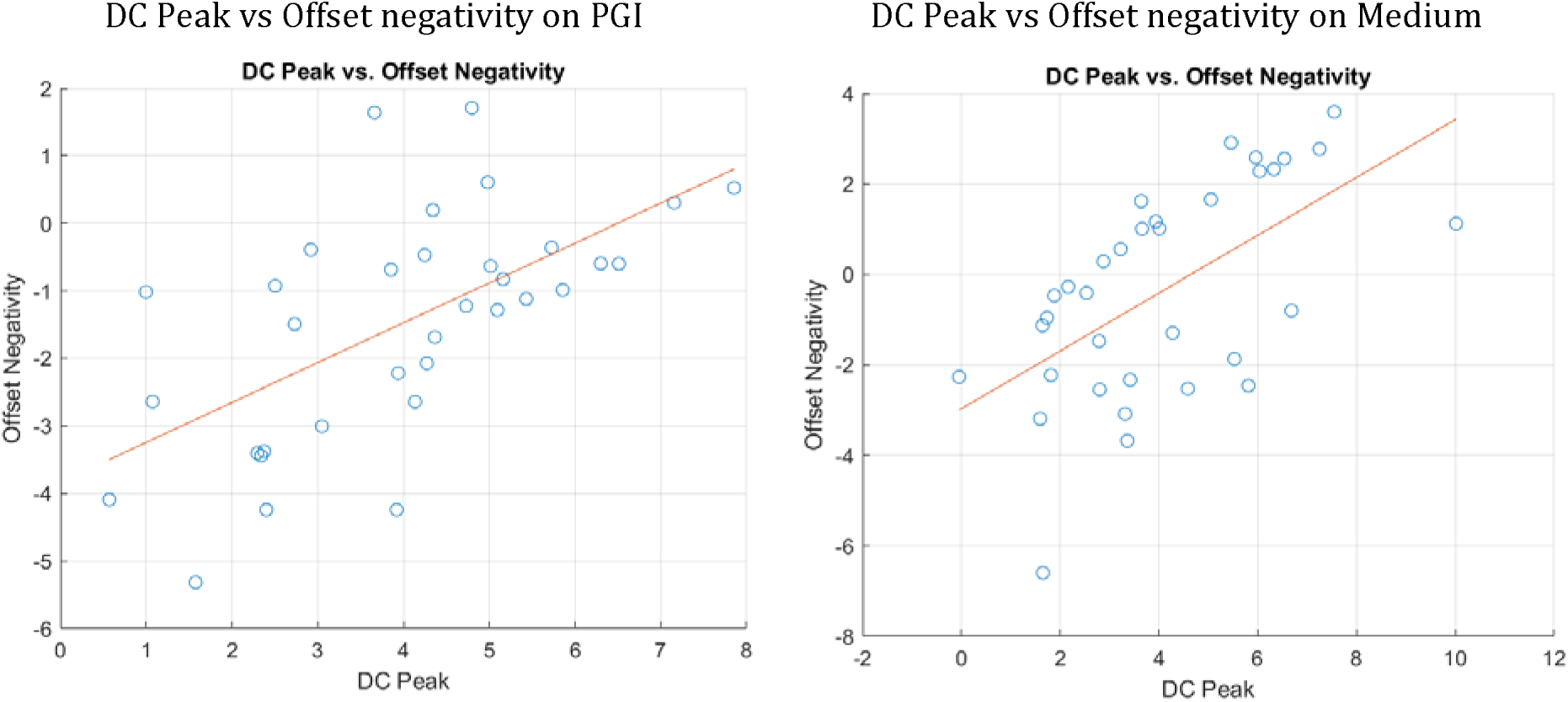
DC shift vs offset correlation. DC peak is plotted from electrode A25 and Offset negativity is plotted at A26. Both plots show the X axis as the peak of the DC shift period and the Y axis as the most negative point in the initial offset period.

## 4 Discussion

Through our analysis, we have discovered multiple features in the grand-averages that, in our data, correlate with the discomfort factor across several dependent variable types. We have also shown effects for the mean/ intercept, most notably in the offset period (see subsection 4.1). There may be more effects in the DC shift period not uncovered by this analysis. This is because statistical inference using FieldTrip favours effects that span a longer period sometimes not finding short powerful effects as these are washed out by the large analysis window. This can be seen with the offset analysis, where the period is split into four, with the largest window of analysis missing results that are short powerful effects like the one shown in Figure 17. We were unable to split the DC shift period in the same way, as it is smooth and does not have any change of stimulus presentation, which causes sharp changes in the participants’ grand-averages, like the ones seen in the offset.

Additionally, it is likely that expectation effects are present in our findings. Specifically, the length of time a stimulus is on for is fixed and it is highly likely that such predictability is detected by the brains of our participants. Indeed, such expectation effects are commonly observed in event related potentials, e.g. the contingent negative variation (Chennu et al., 2013). This does not though, in any way, invalidate our findings, which should be considered within this context. In particular, whether representing predictions or not, the electrical responses we report are all differential across spatial frequencies, exhibiting more extreme responses for the (clinically-relevant) medium stimulus.

### 4.1 Mean/Intercept Effects

The mean/intercept effects correspond to those investigated for the onset transients by (Tempesta et al., 2021). However, here we seek to build on the investigation in that study by looking at the mean/intercept throughout the rest of the grand-average, i.e. for the DC shift and offset periods.

In the DC shift period, there is a large effect that is present throughout the entire period; see Figure 10 and Figure 11. This effect on the medium stimulus follows a similar pattern to the control stimuli but at a higher amplitude, suggesting that the medium stimulus continues to drive the brain throughout the DC-shift period, this finding would fit with findings from Fong et al. who suggest that these striped patterns drive the brain to hyperexcitation (Fong et al., 2020).

We have observed multiple mean/intercept effects throughout the offset analysis windows. Firstly, in the window 3.09 – 3.18s, we see a large negativity for the medium stimulus at around 110ms after stimulus offset (see Figure 17). This effect is potentially notable since no such negativity is observed for the thick and thin stimuli. This sharp negativity may reflect the termination of an inhibitory mechanism that was engaged in response to hyper-excitation during the DC shift period.

The next effect we observe is in the large window 3.09 – 3.99s, which subsumes effects found in windows 3.18 – 3.45s and 3.45 – 3.83s. The effect is a deeper negativity for the medium stimulus compared to the control stimuli (see Figure 16), which likely reflects the greater distance from its higher DC-shift level that the medium stimulus has to fall to return to baseline after the stimulus is removed. Consistent with this interpretation, thick and thin exhibit a similar negativity, but of reduced amplitude compared to the medium stimulus.

### 4.2 Effects on discomfort factor

#### 4.2.1 Discomfort Effect in the offset period

The only significant pure-factor effect was discomfort during the offset (see Figure 19). This manifested as a higher amplitude response to the medium stimulus for the high group, with an accompanying earlier first transient. This may provide a signature for detecting those sensitive to hyper-excitation and would also support findings by Fong et al. who show similar responses to the pattern-glare stimuli for participants self-reporting sensitivity and discomfort to viewing these patterns. Fong et al also demonstrate that those who suffer from clinically diagnosed visually induced migraines show similar patterns (Fong et al., 2020).

#### 4.2.2 Discomfort habituating through partitions

For the partitions, both the Discomfort by Increase analysis in the DC shift period (see Figure 13) and Discomfort by Decrease in the offset period (see Figure 20) yielded similar patterns, where we see an habituation effect for the high group of participants^4^. (Although, the effect was not present for the Medium stimulus in the former of these; see Figure 13c, right column, 2^nd^ row.) These findings suggest that those participants who suffer from higher discomfort, habituate through the course of the whole experiment, whereas those who report lower discomfort seem to have little or no visible habituation effect (indeed, sometimes a reversed effect). This could indicate that those low on the discomfort factor do not experience the same hyperexcitation from the pattern glare stimuli than those who are high on the factor.

There is literature to support the hypothesis of habituation in the visual cortex when presented with visual stimuli (Obrig et al., 2002; Wang and Schoenen, 1998), however, there is conflicting evidence to support this habituation in migraineurs (Adjamian et al., 2004; Judit et al., 2000; Omland et al., 2013; Schoenen et al., 1995), with the majority of the literature suggesting that migraineurs have dysfunctional inhibitory mechanisms leading to no habituation or even sensitisation. Since our study was conducted on a healthy, rather than a clinical, group our results should be validated on migraine suffers, as those who fall in this group may not experience any habituation effect, in line with evidence found in (Adjamian et al., 2004; Schoenen et al., 1995).

Interestingly, focusing further on the DC-shift effects, Figure 12 and Table 2 also suggest an habituation effect through the partitions that is differential for the discomfort factor. However, this habituation is of a negative-going hyper-excitation pattern in partition 1. Thus, it involves a positive-going change through the partitions, back towards, what can be viewed as, a zero baseline, manifesting as a negative habituation effect. Notably, this, high group through the partitions, negative-to-positive pattern in Figure 12 is close on the scalp to the positive-to-negative pattern for Figure 13, with the latter more posterior and closer to the midline. This raises the possibility that there is a dipole reversal between these two effects, which would be consistent with the dipole reversal we observe for the medium stimulus in the mean/intercept (see Figure 10 and Figure 11).

Interestingly, the majority of our discomfort effects were observed in right posterior areas on the scalp (around electrode B12), with some of these at quite different time points: DC-shift versus offset transients. This may suggest a common electrical source. However, in other work, we have observed discomfort effects very posterior on the left side (Dogan et al., 2024). Although, those findings were made with a region of interest analysis, in which the area around B12 was not included. Accordingly, these two different findings may not be as inconsistent as they seem. More research is required to find the generator of these effects. Currently, there is some literature to support the location of these findings (Fischer et al., 2000), however, the findings are linked to habituation when viewing complex visual patterns and not discomfort or pattern-glare.

#### 4.2.3 Sensitisation through the onsets with the discomfort factor

We have found results to support the hypothesis of sensitisation through the fine time granularity in the DC-shift period. We see this effect in Figure 14 and Table 4. This contrast was formalised as a negative going decrease effect for the onsets (2,3 vs 4,5 vs 6,7) in which we see onsets 2,3 and 4,5 negative going for the high group and then onsets-pair 6,7 jumps to be positive going, which we tentatively (see footnote in subsection 3.1.3.1) interpret as a sensitisation effect. A corresponding sensitisation effect was not observed during the offset period.

Sensitisation effects through the onsets that are differentially observed across a factor, here discomfort, are potentially of considerable interest, since they could represent the electrophysiological correlates of the process by which those sensitive to visual stress become aggravated by visual stimuli. The particular pattern of aggravation we observe in Figure 14c, see right side, top two rows, may suggest that this aggravation can obtain in two ways: 1) by simply leaving an aggravating stimulus on (the effect is observed towards the end of the stimulus presentation period) and 2) by repeating that stimulus (sensitisation through the onset).

#### 4.2.4 Three-way interaction

The three-way interaction was setup with the strongest of each of the two-way interactions, increase across the partitions and decrease across the onsets. This is the opposite of our hypothesis in which we predicted that there would be habituation through the partitions and sensitisation through the onsets. The result for the three-way interaction comes out not significant (p = 0.0686), however, this effect looks consistent with our hypothesis for the change of condition through onsets and partitions as the close to significant effect is *negative* going. The grand-averages in Figure 15c show a short-term sensitisation (across the onsets) in the high group that diminishes over the three partitions.

Interestingly, there may be further evidence here of an even finer temporal grain sensitisation for the high group. That is, during the final Onset-pair (6,7), there seems to be a progressive increase in PGI and the response to the medium stimulus during the DC-shift time period. This suggests that for those sensitive to the stimulus, continuing to “drive” visual cortex with a sustained pattern-glare stimulus causes the brain’s response (indeed, its level of hyper-excitation) to continue to ramp upwards, with this obtaining over a 3 second period of time.

There may also be habituation during the first partition for those low on the factor. That is, for this group that are not sensitive to discomfort, the first Onset-pair (2,3) may be exhibiting a hyper-excitation response late in the DC-shift time period, which dissipates by the second Onset-pair. Thus, it may be that the well-functioning brain is initially aggravated by the pattern-glare stimulus (i.e. for early presentations of the medium stimulus at the start of the experiment), but is then able to habituate to that stimulus extremely rapidly? (The question that remains about this low group pattern is if it does reflect hyper-excitation, why does that particular pattern, i.e. an early Onset-pair, early in experiment, exhibiting an elevated response, not obtain for the high group? One might have thought that the high group would exhibit this pattern to a very marked extent.)

## 5 Conclusion

Our findings suggest that participants who reported greater discomfort to the pattern-glare stimulus displayed sensitisation at a fine time granularity and habituation at a longer granularity. This suggests the presence of sensory impairment, which is consistent with current literature on cortical hyperexcitability. Although, the particular pattern that we observed, short term sensitisation and long-term habituation, is new.

A fundamental question that remains is whether the effects we observe reflect the direct manifestation of hyper-excitation or alternatively, inhibitory processes initiated in response to that hyper-excitation. Investigations in the frequency domain, where particular frequencies have been associated with inhibition (Mathewson et al., 2011), may help to understand this question. Additionally, it needs to be recognised that the research presented here is fundamentally exploratory: we have performed a number of comparisons (see section 9.2.4 of the appendix) to identify our main findings (although many of these were correlated) and thus, our effects need to be replicated before they can be considered robust.

## 7 Author Contributions

Tom Jefferis wrote the analysis code, analysed the data, interpreted the results, wrote the first draft and contributed to later drafts. Cihan Dogan contributed to preprocessing code and analysis code, and contributed to later drafts. Claire E. Miller contributed to experiment and preprocessing code, contributed to later drafts, and collected the data. Maria Karathanou contributed to the analysis code, and found initial results for the DC shift period. Austyn J. Tempesta conceived the study, wrote experiment code, contributed to later drafts. Howard Bowman conceived the study, wrote experiment code, interpreted results, and contributed to all drafts.

## Footnotes

1 c/deg or cycles per degree is a measure of spatial frequency and is equal to the number of cycles of a grating (one dark and one light band) that subtends an angle of one degree at the eye (“Dictionary of Optometry and Visual Science, 7th edition,” 2009).

2 Due to the way Fieldtrip sets up statistical inference for a one-sample t-test, it is required to duplicate the data object, and to replace all the duplicate’s functional values with zeros. Then this zeroed data is assigned a different integer in the regressor (Figure 5). A two-sample t-test is then run over this regressor, which simulates a one-sample t-test over a standard (all-ones) intercept regressor.

3 The pattern observed in Figure 14c, right panels (high group), is difficult to definitively interpret. One could view the top right panel of Figure 14c as habituation through the onsets from a negative deflection (at early onsets) upwards or sensitization through the onsets towards a positive deflection (at late onsets). We marginally prefer the latter interpretation, because the response to medium (second row), which is more easily interpreted, since it is not calculated across many stimulus types (as the PGI is), ends above zero at Onset-pair 6:7.

4 In the DC-shift period, Figure 13, this comes out as a *negative*-going effect on an increase interaction, which due to its negative-direction can be interpreted as an habituation effect.

