## Appendix Material for "Beyond EEG Onset Transients: Sensitisation and Habituation of Hyper-excitation to Constant Presentation and Offset of Pattern-Glare Stimuli"

**Appendix / Supporting material**

**Methods**

**Factor Analysis**

An important point to consider is the reliability of the factor analysis we have performed. This section provides evidence for this reliability.

Firstly, we note Kaiser’s rule, which suggests having at least five times as many observations as you have variables, will ensure a reliable fit. In our case, we have seven variables, so we meet the threshold of 35 participants (Kaiser; 1974), since we have fitted to 39 participants. Although, there are conflicting opinions on this issue, e.g. Costello & Osborne (2005).

However, all such “rules of thumb” are limited in their applicability, since they do not reflect the characteristics of any particular data set. Indeed, a data set with less noise variability is going to give more reliable factor analyses than one with more.

**Background to Analysis**

In justifying the reliability of our factor analysis, our basic approach is to compare the decomposition into factors and the uncertainty re. this decomposition, with what we would observe if we performed the only real alternative approach to a data-driven decomposition, such as factor analysis. This alternative is what we call a *flat average*.

More specifically, in order to identify electrophysiological features that correlate with condition-relevant features (e.g. headache susceptibility), we need regressors that classify our participants according to such features. One could just include regressors that are single variables from those we have collected, e.g. Headache-intensity, but that would throw away a good deal of information collected. One would also expect that aggregating across multiple response variables, would provide a more robust measure. Additionally, one wants an aggregation procedure that provides orthogonal regressors, which when entered into a regression model explain non-overlapping variability in the data. Without orthogonality of regressors, interpretation of findings is challenging.

As previously discussed, to do this, one is really left with two options, especially if one wants to restrict to a linear association of (condition-relevant) variables to components:

1) flat averaging of relevant variables, based upon an intuition of the condition-relevant features of each variable; and
2) data-driven selection of weighting coefficients by a procedure such as PCA or factor analysis.

We contend that there are only really two plausible flat average patterns as depicted in Figure 22. These patterns differ only in the assignment of aura to the visual stress or headache factors respectively. Phenomenologically and perhaps, mechanistically, one might consider aura to fit with visual stress, where susceptibility to spurious visual percepts and consequences of hyper-excitation load. Intuitively, one might also think that aura would associate with the headache variables (H-duration, H-intensity, H-frequency), since it is an experience that occurs before and during migraine headaches. Effectively, our data-driven approach has provided a clear answer to this question; see subsection “Comparison to flat average”.

If one believed that our factor analysis was unreliable, it would generate a pattern of factor loadings that are substantially subject to noise. There are two obvious ways in which this noise might manifest:

1) It would cause the pattern of factor loadings to be quite different from the (a priori) intuitive flat average loadings.
2) It would lead to high variability/ uncertainty in the loadings that would be produced.

In the next two subsections, we explore whether there is any evidence of these manifestations of noise.

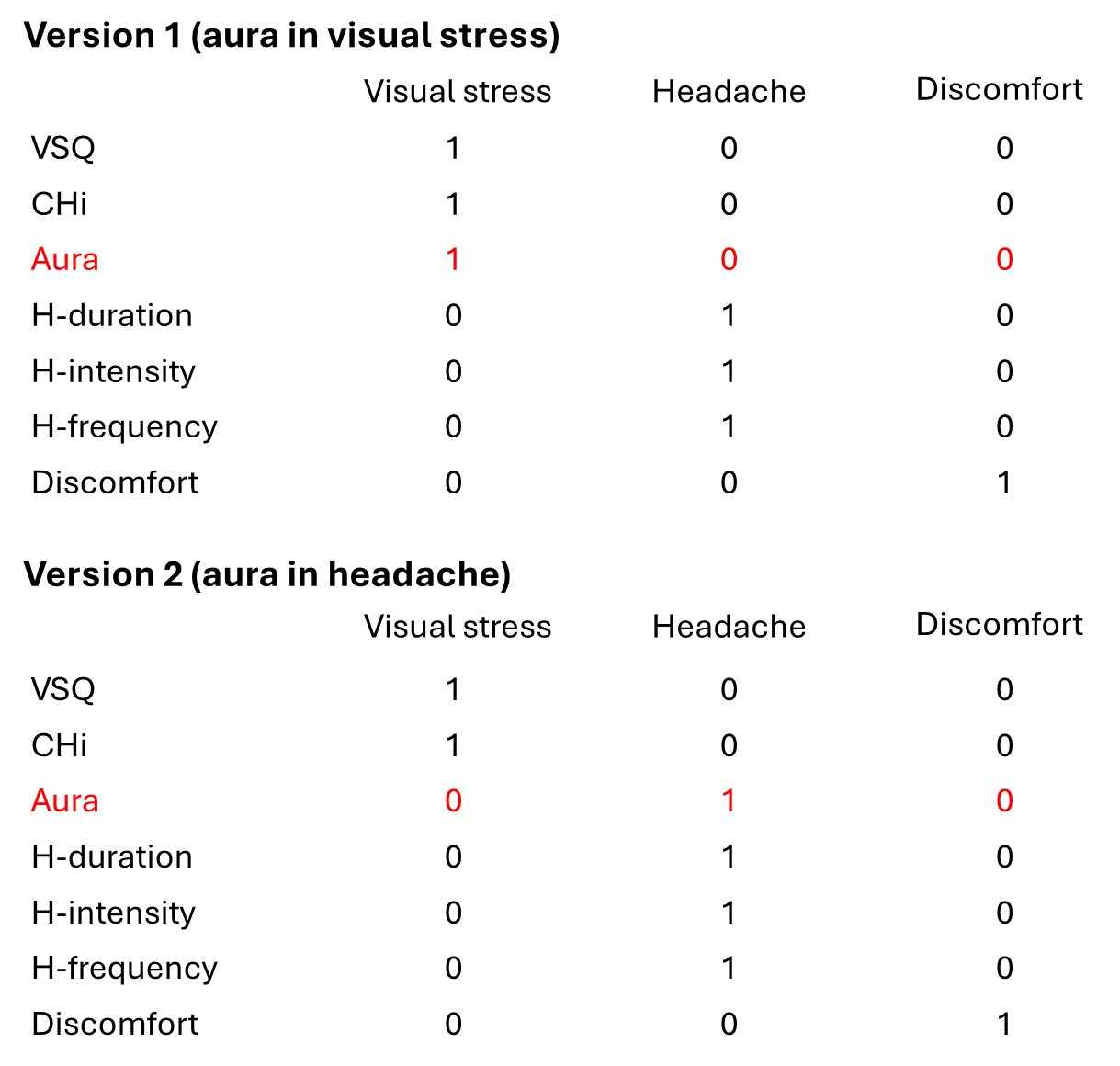

*Figure 22: The two (intuitively) plausible flat-average associations of (condition-related) variables to components. CHi is the cortical hyperexcitability index (CHi); VSQ is …..; H-duration is Headache duration and similarly for H-intensity and H-frequency. These two versions agree on the assignment of six out of seven (condition-related) variables, i.e. VSQ and CHi to visual stress; H-duration, H-intensity and H-frequency to Headache; and Discomfort to Discomfort. The assignment of Aura (see rows in red) is the point of contention: it could plausibly be associated with either visual stress or headache, because it exhibits symptoms that one might associate with the CHi and VSQ questionnaires, but occurs with headaches. An objective of our factor analysis was to provide a data-driven answer to the question of which component Aura should associate with.*

**Comparison to flat average**

Our application of factor analysis gave the weighting coefficients across our seven variables in Figure 23.

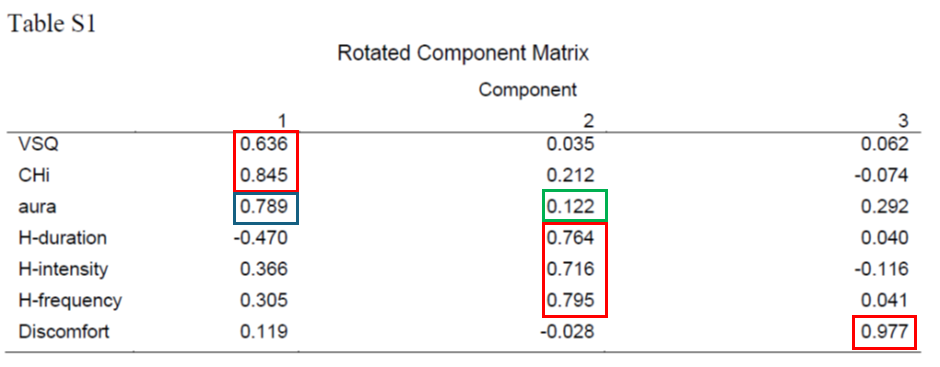

*Figure 23: factor loadings obtained through factor analysis on observed data.*

Importantly, these results are not substantially different to those we would obtain from flat averaging (see red rectangles). Although, as previously discussed, there is one condition-related variable (aura), for which there is uncertainty about its association (from an intuitive perspective) with components (see blue and green boxes).

Thus, our data-driven approach has provided loadings, which would correspond to the weighting matrix shown as version one (aura in visual stress), in Figure 22.

The correspondence to our derived weightings is especially the case if one notes that the positive and negative headache variables on the first factor/component cancel in our factor loadings. That is, the overall effect of the three headache variables on the first factor is going to be much closer to zero than might be suggested by the absolute values of each of the three coefficients. This is because one is negative and the other two are positive.

In fact, it should be reassuring that a data driven approach like factor analysis, identifies a factor structure so close to one’s intuitive expectations.

Effectively, the specific contribution of our data-driven approach is to provide a clear answer (loading in blue box is much higher than in green box) to the point of uncertainty we were faced with: which component should aura be associated with?

Aura correlating with visual stress is quite plausible with our population, who are sub-clinical. That is, it is likely that we have many participants who have headaches that are *neither* visually-induced nor associated with visual aura. For these participants, aura may be irrelevant to the frequency and features of their headaches. However, for others, the presence or absence of aura may be strongly correlated with the experience of visual stress-related percepts.

Thus, we would argue that the factor loadings we obtain are not speculative: the two possible approaches, data-independent or data-driven, are in the main very consistent in their outcomes, with the data-driven approach enabling us to determine the most appropriate component that aura should be allocated to in our participants.

**Variability investigation**

As previously discussed, if our factor analysis was especially subject to noise, we would expect this to manifest as instability in the factor loadings generated across replications. We seek to quantify this instability in this subsection and compare it to the instability found in the flat average approach, and we include a PCA decomposition for comparison.

*Methods*

To explore this instability, we bootstrapped our data, as one would do when quantifying error using confidence intervals. Thus, we started with our data represented as a 39 rows (participants) by 7 columns (variables) data structure. We then sampled each column with replacement to generate 50,000 (bootstrapped) surrogate data sets. On each such surrogate data set we ran three analyses, flat average, PCA and factor analysis, each of which projected the (39 x 7) surrogate data onto three components, giving a 39 x 3 projected surrogate data. More specifically, we performed the following procedures on each surrogate data:

1) *Flat average*: we projected onto three components, by applying the version 1 (flat average) weight matrix to the surrogate data, i.e. row *i* in the first column of the projected surrogate data was the average of the first three columns (VSQ, Chi and Aura) of the *i*th row of the surrogate data, and so on for the next two columns. This gave us the *flat average projected data*.

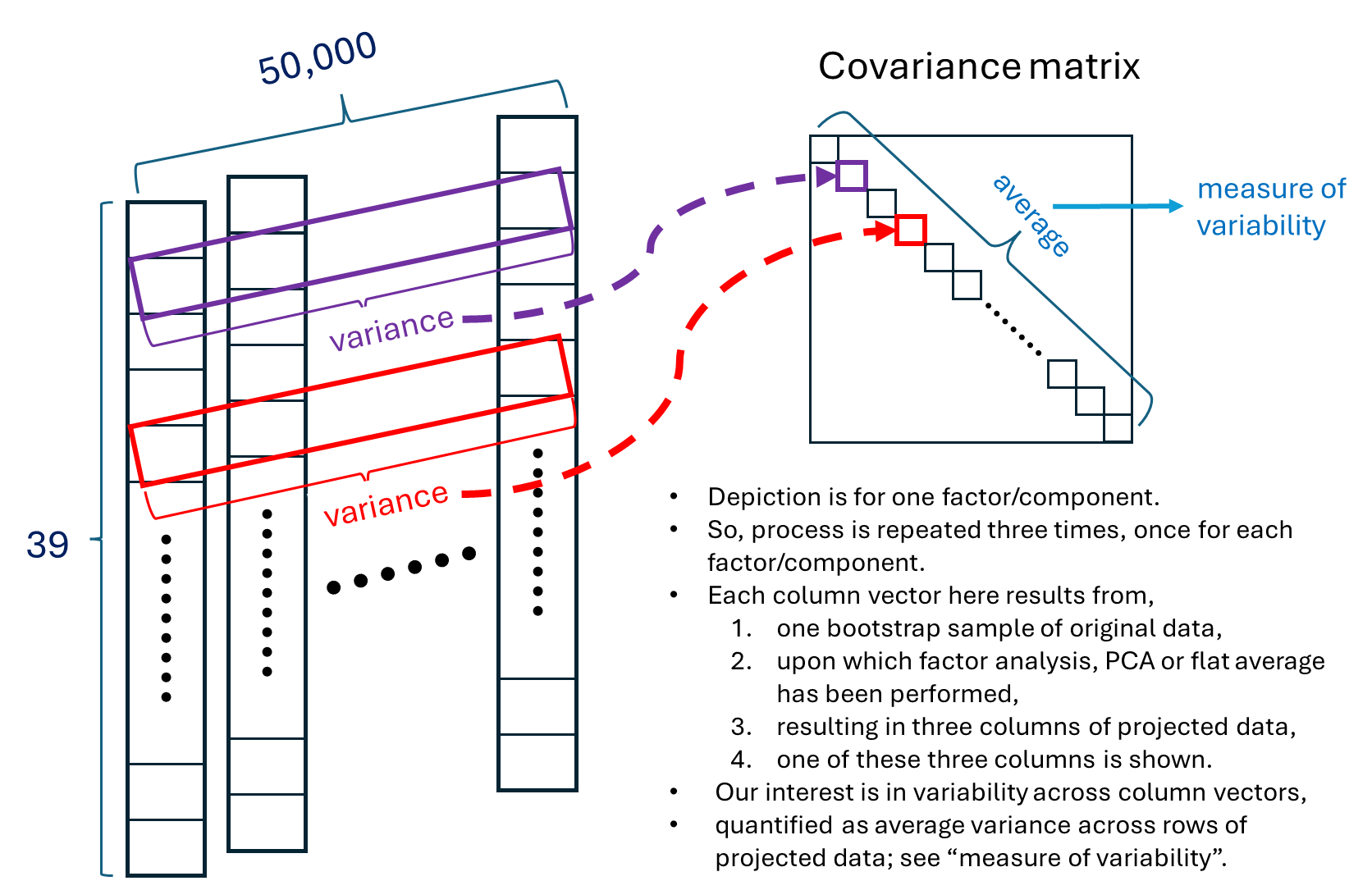

*Figure 24: method for calculating variance induced by a particular method on projected data. 39 is the number of participants, 50,000 is the number of bootstrap resamplings. The main diagonal of the covariance matrix contains the variances of each row of 39x50,000 data structure. Each row corresponds to a (surrogate/bootstrapped) participant. The average of the variances across these rows (i.e. the trace of the covariance matrix divided by the number of rows) is a measure of variability/uncertainty resulting from projecting our data through either factor analysis, PCA or flat average loading/projection matrix.*

2) *PCA*: we performed PCA decomposition on the surrogate data set, giving us a 7x3 matrix of component loadings/ weights. The surrogate data was then projected onto the three component dimensions using this matrix, giving us a 39x3 data set. This is the *PCA projected data*.
3) *Factor analysis*: we performed an exploratory factor analysis on the surrogate data set, giving us a 7x3 matrix of factor loadings/ weights. The surrogate data was then projected onto the three factor dimensions using this matrix, giving us a 39x3 data set. This is the *factor analysis projected data*.

So, for each of these procedures, we obtained 50,000 39 x 3 projected surrogate data sets. We are interested to quantify the variability across these 50,000. We do this separately for each of the three dimensions/components/factors projected onto, i.e. on 50,000 39 x 1 data structures. Thus, we are interested in the variability across these 50,000 column vectors, i.e. a 39 x 50,000 data structure; see Figure 24. We quantify this variability by constructing the 39 x 39 covariance matrix across this 39 x 50,000 matrix; see figure 24. The trace of this covariance matrix is the sum of the variability across each of the 39 rows. Note, off-diagonal elements in the covariance matrix are close to zero, since the 50,000 is big (because the bootstrap sampling is random). Consequently, there is effectively no covariance across rows in figure 24, to be concerned about.

This gives us a sum of variances for each of the three components/ factors, which we turn into an average. If these three averages of variances are substantially bigger for factor analysis than it is for the flat average, it would suggest that the factor analysis adds uncertainty/error.

*Results*

In this section, we present the average along the trace from each covariance matrix across all bootstrapped samples for each of the three procedures for projecting onto three components. These results are presented in Table 8.

*Table 8: Average along trace from each covariance matrix across all bootstrap samples for each of the three procedures for projecting onto three components/factors (indicated 1, 2 and 3 here): (flat) Averaging, principal component analysis (PCA) and exploratory factor analysis (EFA). Additionally, the total averaged trace across all three dimensions is presented. These dimensions would be called components for PCA and factors for EFA.*

| **# Bootstraps (50,000)** | **1** | **2** | **3** | **Total Average Trace** |
| --- | --- | --- | --- | --- |
| **Averaging** | 0.6 | 0.91 | 0.88 | **2.39** |
| **PCA** | 2.62 | 1.57 | 0.95 | **5.14** |
| **EFA** | 0.91 | 0.91 | 0.89 | **2.71** |

As expected, the variability reduces across the PCA components – mathematically, it is placing each component at the highest variance dimension of the data remaining after projecting out previous components. Additionally, although to a lesser extent, variability reduces across factors, since more explanatory variables are again earlier. The total of the three average variabilities along the trace (the last column) is smallest for flat averaging, slightly higher for EFA and then substantially higher for PCA. This total is an estimate of the error/uncertainty associated with each procedure, and importantly, based on this analysis, experimental factor analysis does not substantially increase the error over that observed from our baseline procedure, (flat) averaging. That is, the flat average is the plausible alternative to using factor analysis and one might believe that its simplicity (and the fact that loadings are fixed) would make it substantially less susceptible to error. However, this is not what we see – with our bootstrapping procedure, factor analysis exhibited only a little more error variance than flat averaging. This increase in variability almost certainly arises because factors are ordered by their explanatory value, which relates to variance explained. We are only looking at the first three factors, which will necessarily have higher explained variance.

However, our central point is clear, even though the factor analysis loading matrix is derived from the data and changes between data sets, it does not add substantially more variability to the flat average, which is a fixed loading matrix. This suggests that, with our data, factor analysis is not a great source of added uncertainty.

*Algorithm*

To estimate variability in these methods we employed a bootstrapping procedure, i.e. using sampling with replacement. The algorithm is listed below:

1. **Initialisation:**
   a. Set the number of bootstrapped datasets to generate k=50,000.
   b. Define the number of factors/components for EFA/PCA (3).
   c. Instantiate three vectors of size (number_of_bootstraps, number_of_participants, 3) to store results for the averaging, PCA and EFA procedures.
2. **Bootstrapping Procedure:**
   a. For i from 1 to k:
      i. Generate a bootstrapped sample of size 39x7 by sampling with replacement from the original dataset of 39 participants and 7 variables.
      ii. For each method (averaging, PCA, EFA):
          1. Average
             a. Calculate the following three factors:

i. Visual Stress: Mean of VSQ(z), Total Chi(z), and Aura(z)

ii. Headache: Mean of Headache Duration(z), Headache Intensity(z), and Headache Frequency(z)

iii. Discomfort: Discomfort Index(z)

b. Store each of the three factors in the averaging vector i.e.

i. averaging_results(i, :, 0) = Visual Stress

ii. averaging_results(i, :, 1) = Headache

iii. averaging_results(i, :, 2) = Discomfort

2. PCA:
   a. Apply Principal Component Analysis (PCA) to the bootstrapped dataset of size 39x7.
   b. Transform the dataset to a reduced dimensionality of size 39x3 using the first 3 principal components.
   c. Store each of the three components in the PCA vector i.e.

i. pca_results(i, :, 0) = PCA1

ii. pca_results(i, :, 1) = PCA2

iii. pca_results(i, :, 2) = PCA3

3. EFA:
   a. Apply Exploratory Factor Analysis (EFA) to the bootstrapped dataset of size 39x7.
   b. Extract 3 factors and transform the dataset to a reduced dimensionality of size 39x3.
   c. Store each of the three components in the EFA vector i.e.

i. efa_results(i, :, 0) = Factor 1

ii. efa_results(i, :, 1) = Factor 2

iii. efa_results(i, :, 2) = Factor 3

3. **Compute the covariance matrix of each one of the methods**
   i. For each method, calculate the covariance matrix for its own respective factor, component, score i.e.
      1. for Visual Stress: covariance_matrix(averaging_results(:,:,0))
      2. for Headache: covariance_matrix(averaging_results(:,:,1))
      3. for Discomfort: covariance_matrix(averaging_results(:,:,2))
4. **Given the covariance matrix compute the average trace and present the results.**

**Change through time regressors:**

We will now explain the steps to create the exponential change through time regressors. Continuous regressors representing the change through time were used as predictors. We considered two different profiles of change through time: decrease and increase. Decrease reflects an exponential decrease, whereas increase reflects an exponential increase, given in the following form:

$$Y_{i,inc}=e^{X_{i}} and$$

$$Y_{i,dec}=e^{(1-X_{i})},$$

$$where, i\mathbb{\in N (}1\leq i\leq3) s.t. X_{i}=\frac{\left( i-1 \right)}{2},$$

$$according to which X_{1}=0, X_{2}=0.5, X_{3}=1$$

Thus, we raise $e$ to the power of a linear decrease in the decrease regressors, and a linear increase in the increase regressors. The resulting exponential decrease or increase was selected as neural responses are typically better described by exponential changes, reflecting the non-linearity of neuron firing. The regressors, after mean-centring, are visualised in Figure 25, where $time\_point i=Y_{i,dec}$ on the left-hand side and $time\_point i=Y_{i,inc}$ on the right-hand side.

| 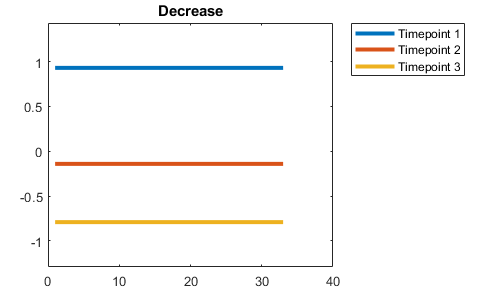 | 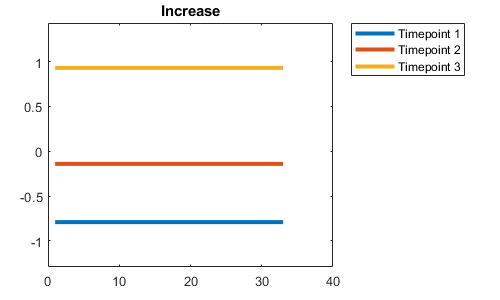 |
| --- | --- |

*Figure 25: Change through time regressors. (Left) A regressor representing decrease through time, (Right) a regressor representing increase through time. The x-axis represents the participants and the y-axis the design matrix value.*

**Results**

For the results in this section, we will present unorthogonalized factors that the MUA found to have significant or close to significant clusters.

**DC Shift Period**

***Average of Onsets 2-8***

**Discomfort Factor:** There were two effects that were close to significance in the DC shift period that came out of the MUA for the average of onsets 2-8 for the discomfort factor. The first (significant) cluster is small, only occupying 15% of the volume at its maximum point (Figure 26a). The effect does last around 500ms, but when visualised with grand averages, it is not very compelling, not showing much difference between the low and high groups, without substantial separation of plotted confidence intervals (see left column of plots row 2 and 3 of Figure 26c). The effect looks particularly weak when the medium stimulus is plotted alone (row 3 of Figure 26c), suggesting an effect that is somewhat carried by thick and thin. However, interestingly, the effect discovered by the MUA is in a similar location to other discomfort effects discussed (see Table 9).

The second of these clusters happens just before the first cluster, starting at around 1.75s and lasting around 250ms. The cluster is weak, with a p value of 0.0862. When looking at the topographic plots in Figure 26b, this cluster is the dark blue area in the 4^th^ map. It is biggest at the same location as the later more prominent cluster (B5), suggesting it is part of the same effect as that first cluster, and that the time-series in Figure 26c are representative of it. All of this indicates that the effect is not particularly notable.

*Table 9: MUA results for discomfort on the average of onsets 2-8 regressor. Only results for clusters containing significant effects (FWE-corrected) or borderline effects smaller than a p-value of 0.1 are shown, both positive and negative tails.*

| Effect | Tail (+1) | Tail (-1) |
| --- | --- | --- |
| Discomfort 0.5-3.0s | No significant Cluster | 1^st^ cluster p-value: 0.0584  1^st^ electrode: B5  1^st^ peak time: 2.3379  1^st^ r correlation effect size: -0.64  1^st^ Cohens d effect size: -1.68  2^nd^ cluster p-value: 0.0862  2^nd^ electrode: B5  2^nd^ peak time: 1.9122  2^nd^ r correlation effect size: -0.54  2^nd^ Cohens d effect size: -1.283 |

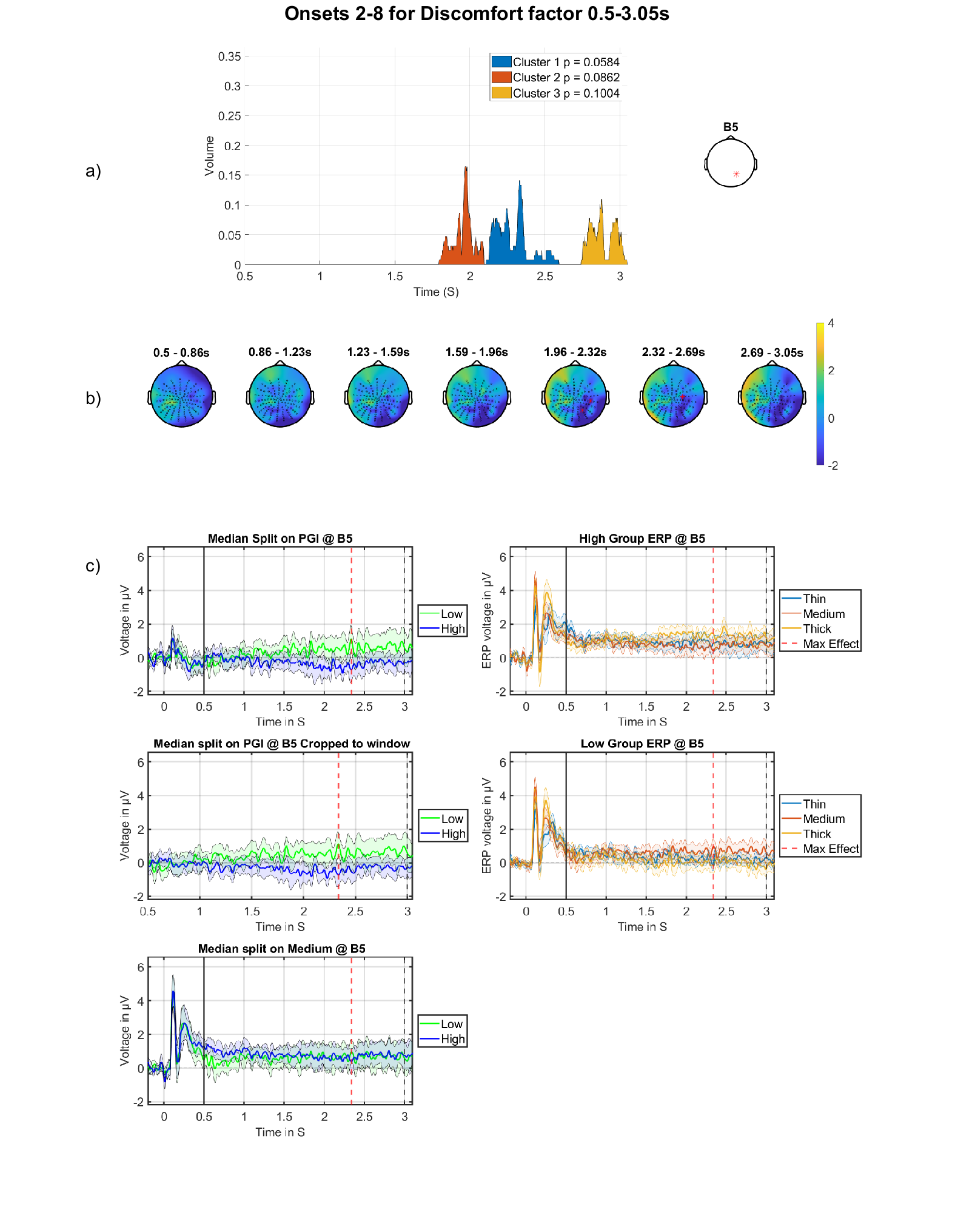

*Figure 26: DC shift effect for the discomfort factor on the average of the onsets. a) Cluster volume as a percentage of the entire scalp. The electrode used for plotting is displayed on the right. b) Topographic maps through time for the whole period, with red crosses indicating the significant cluster, which corresponds to the blue region in a). c) grand-averages at electrode indicated on right in panel (a), with maximum effect marked with the red dashed vertical line and window start with a black solid line. Top left is the high vs low discomfort group for the PGI; middle left are the grand-averages for high vs low discomfort group for the PGI with window showing only the period of analysis; bottom left are the high vs low group for the medium stimulus; top right are the grand-averages for the high group showing thick, medium and thin; bottom left are the grand-averages for the low group showing thick, medium and thin.*

***Onsets 2,3 vs 4,5 vs 6,7 Unorthogonalized results***

**Discomfort-by-decrease:** The cluster found in the MUA for this regressor was comparable to the orthogonalized regressor for the same contrast shown in Table 4 and Figure 14, however, with a less significant p value (0.007 vs 0.001). This could be because of the unorthogonalized regressor. The cluster is almost identical to that described in Table 4 and shown in Figure 14.

*Table 10: MUA results for the discomfort by decrease through the onsets with unorthogonalized regressor. Only results for analysis windows containing significant effects are shown, both positive and negative tails.*

| Effect | Tail (+1) | Tail (-1) |
| --- | --- | --- |
| Discomfort by decrease 0.5-3.0s | No significant Cluster | 1^st^ cluster p-value: 0.007  1^st^ electrode: B16  1^st^ peak time: 2.8008  1^st^ r correlation effect size: -0.42  1^st^ Cohens d effect size: -0.92 |

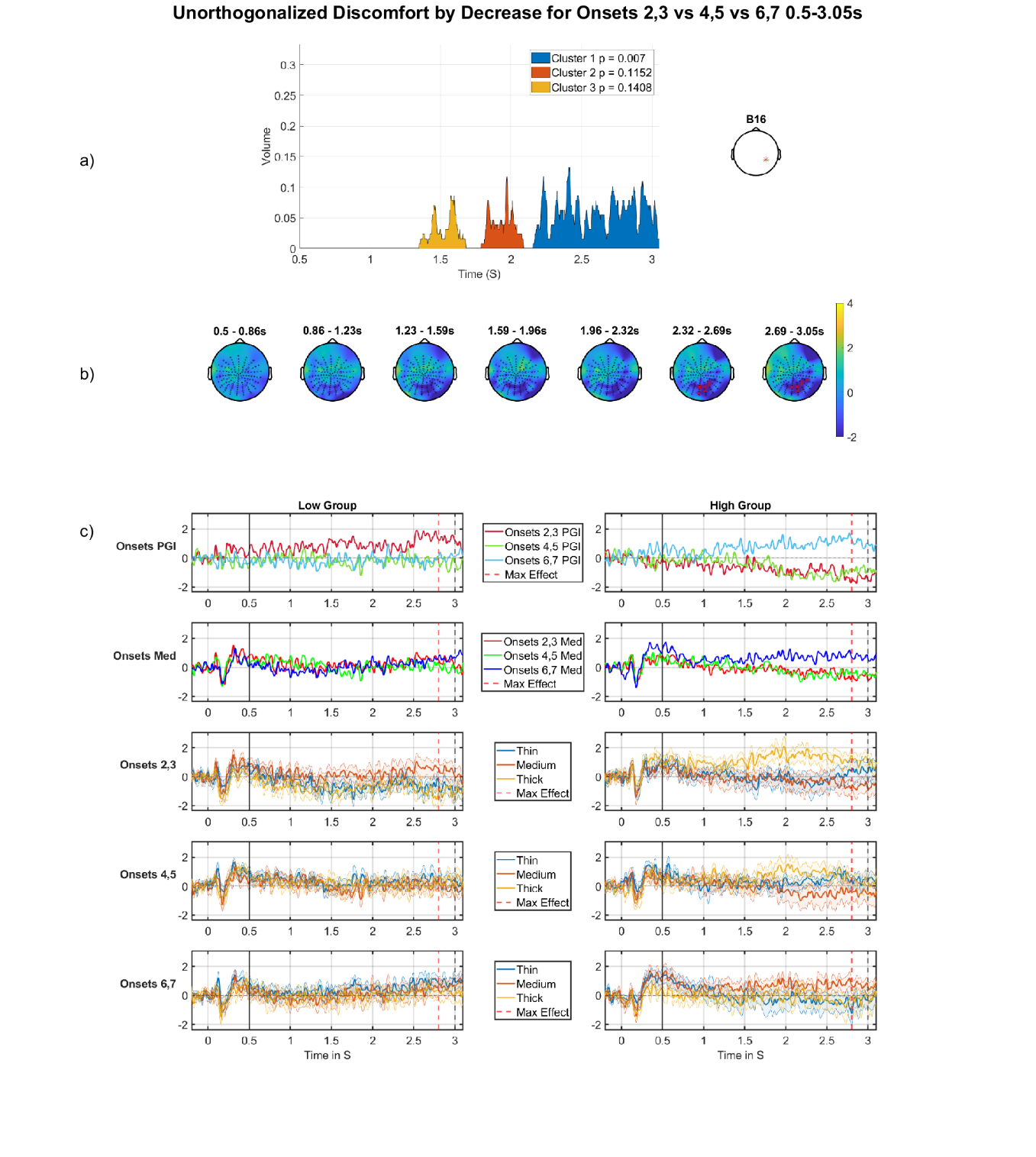

*Figure 27: Discomfort with a decrease across Onsets, negative cluster, DC shift period. A) Cluster volume as a percentage of the entire scalp, the electrode used for plotting displayed on the right. B) Topographic maps through time for the whole period, with red crosses indicating the significant cluster, which corresponds to the blue region in a). c) Median split (on Discomfort) grand-averages at electrode indicated on right in panel (a), the left column of grand-averages is for the low group, right column of grand-averages is for the high group. Top is the grand-average for the PGI for each partition, red (partition 1), green (partition 2), blue (partition 3) with maximum effect marked with a red vertical line, window start marked with a black solid line and stimulus off marked with a black dashed line; second row is the grand-average for the medium stimulus for each partition, red (partition 1), green (partition 2), blue (partition 3); third, fourth and fifth rows present grand-average for partitions 1, 2 and 3 (respectively), each showing thin, medium and thick. For the high group.*

**Offset Effects**

***Mean/intercept effects on the average of onsets***

The table below (Table 11) shows the effects in section 3.2.1.1 of the main body, summarising all mean/intercept effects in the offset period, only three of which are presented in section 3.2.1.1.

*Table 11: MUA results for mean/intercept effect for the average across all onsets in the offset period. Only results for clusters containing significant effects (FWE-corrected) or borderline effects smaller than a p-value of 0.1 are shown, both positive and negative tails.*

| Effect | Tail (+1) | Tail (-1) |
| --- | --- | --- |
| Mean/intercept 3.08-3.99s | No significant Cluster | 1st cluster p-value: 0.0004  1st electrode: A24  1st peak time: 3.5215  1st r correlation effect size: -0.71  1st Cohens d effect size: -2.01 |
| Mean/intercept 3.09-3.18s | 1st cluster p-value: 0.0384  1st electrode: B30  1st peak time: 3.1152  1st r correlation effect size: 0.66  1st Cohens d effect size: 1.75 | 1st cluster p-value: 0.0076  1st electrode: A26  1st peak time: 3.1191  1st r correlation effect size: -0.83  1st Cohens d effect size: -2.98 |
| Mean/intercept 3.18-3.45s | No significant Cluster | 1st cluster p-value: 0.0032  1st electrode: A25  1st peak time: 3.2949  1st r correlation effect size: -0.69  1st Cohens d effect size: -1.9 |
| Mean/intercept 3.45-3.83s | No significant Cluster | 1st cluster p-value: 0.0006  1st electrode: A24  1st peak time: 3.5215  1st r correlation effect size: -0.71  1st Cohens d effect size: -2.01 |

The figures presented below are for the mean intercept effect in windows 3.18-3.45s and 3.45-3.83s, which were parts of the same cluster as found in the 3.09-3.99s analysis window. We can see this by firstly comparing the cluster volume plots (a panels) in Figure 16, with those of Figure 28 and Figure 29, where you can see almost the exact same shape, but “zoomed in”. This is also supported by the topographic maps displaying the clusters identified being at the same location on the scalp and occurring at corresponding time periods.

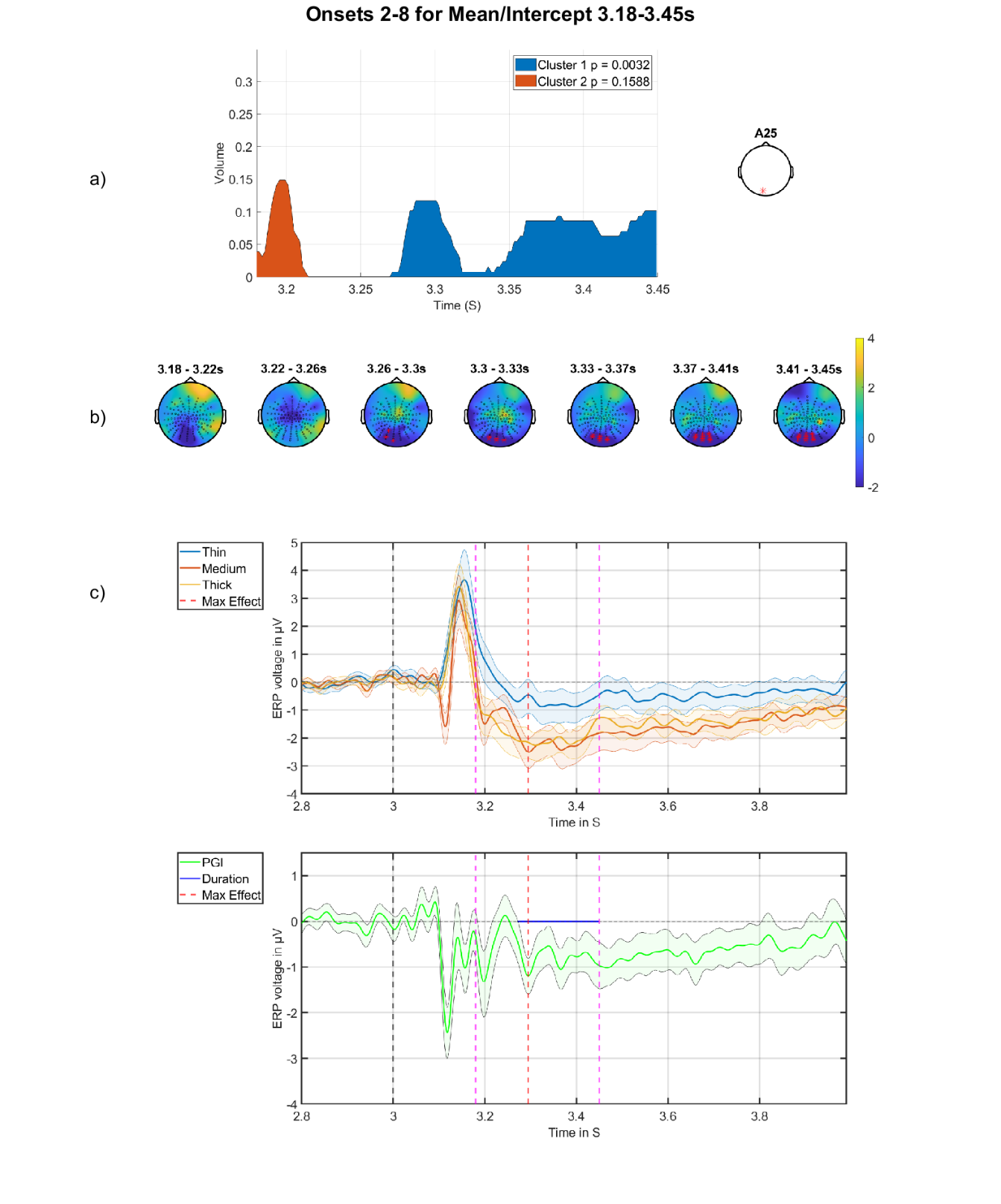

*Figure 28: Mean/intercept of offset, in period 3.18-3.45s. a) Cluster volume as a percentage of the entire scalp,* *with the electrode used for plotting displayed on the right. B) Topographic maps through time for the whole period, with red crosses indicating the significant cluster, which corresponds to the blue region in a). c) grand-averages at electrode indicated on right in panel (a), with time of maximum effect marked with red vertical dashed line and window of analysis marked with pinked dashed lines. Top are the grand-averages for thick, medium and thin, with stimulus offset marked at three seconds with a black dashed line; bottom is the grand-average of the PGI, with the blue horizontal line indicating the period of statistical significance.*

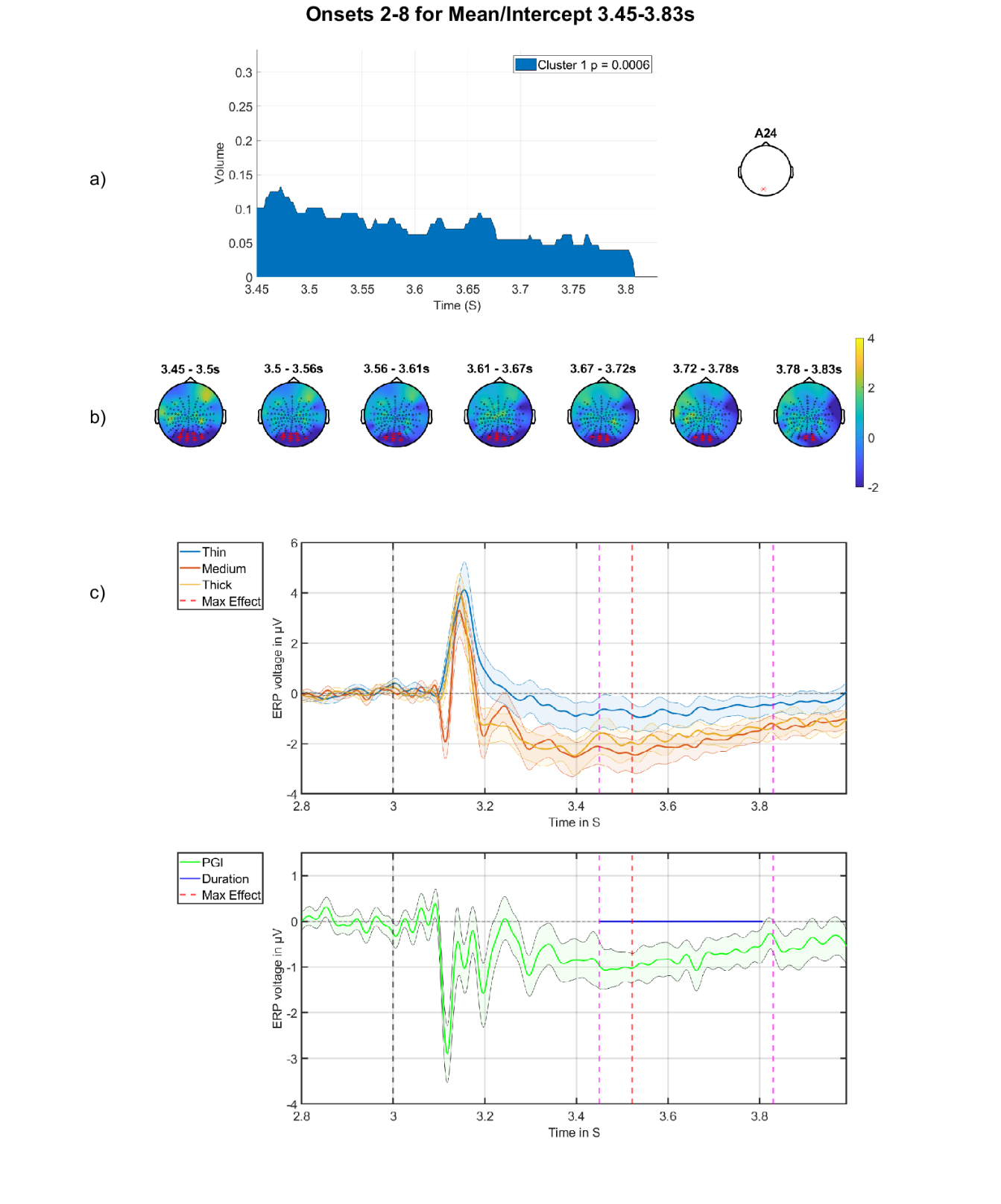

*Figure 29: Mean/intercept of offset, in period 3.45-3.83s. a) Cluster volume as a percentage of the entire scalp, with the electrode used for plotting displayed on the right. B) Topographic maps through time for the whole period, with red crosses indicating the significant cluster, which corresponds to the blue region in a). c) grand-averages at electrode indicated on right in panel (a), with time of maximum effect marked with red vertical dashed line and analysis window marked with pink dashed lines. Top are the grand-averages for thick, medium and thin, with stimulus offset marked at three seconds with a black dashed line; bottom is the grand-average of the PGI, with the blue horizontal line indicating the period of statistical significance.*

***Discomfort over average of Onsets 2-8 results***

Figure 30 is a continuation from Figure 19, which shows the median split for discomfort plotted at the most significant electrode for the mean/intercept effect in the offset found in Figure 17.

Figure 30 is included to determine if there was any effect of the discomfort factor on the N120. Although not significant, a difference between the two groups is shown, with the high group having a shallower negativity than the low group, the opposite pattern to what would be naturally expected of a hyper-excitation explanation.

d)

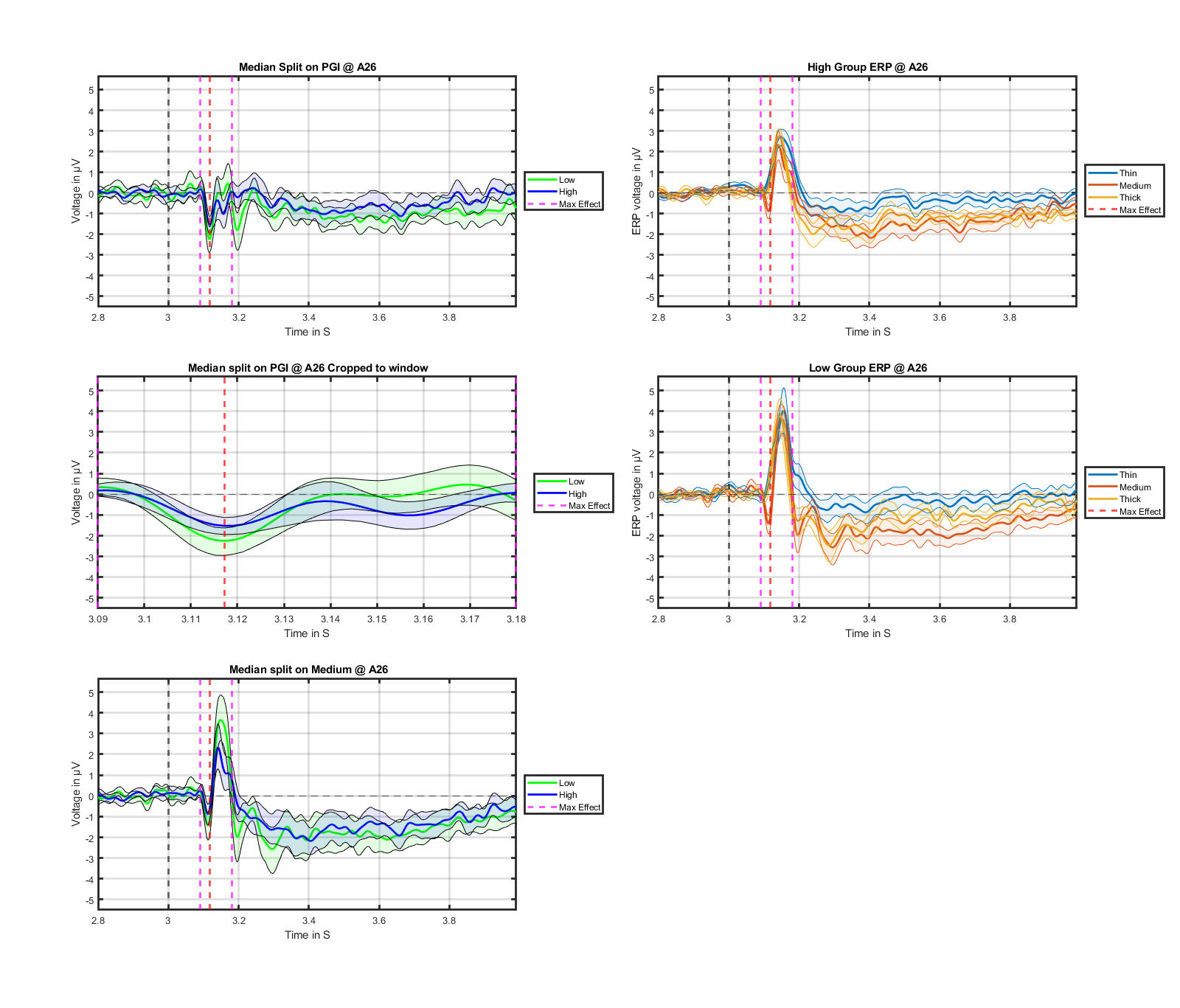

*Figure 30: Continued figure from Figure 19. d) grand-averages at electrode A26, with maximum effect marked with the red vertical line. Top left is the high vs low discomfort group for the PGI; middle left are the grand-averages for high vs low discomfort group for the PGI, with window showing only the period of analysis; bottom left are the high vs low group for the medium stimulus; top right the grand-averages for the high group showing thick, medium and thin; bottom right are the grand-averages for the low group showing thick, medium and thin.*

***Pure change through time***

Next, we present the MUA results for the pure change through time effect. The only significant result found during MUA is for the window 3.45s-3.83s shown in Table 12.

*Table 12: MUA results for the pure decrease through time across the partitions for the time period 3.45 – 3.83s. Only results for clusters containing significant effects (FWE-corrected) or borderline effects smaller than a p-value of 0.1 are shown, both positive and negative tails.*

| Effect | Tail (+1) | Tail (-1) |
| --- | --- | --- |
| Pure decrease in partitions 3.45-3.83s | 1^st^ cluster p-value: 0.0316  1^st^ electrode: D22  1^st^ peak time: 3.5835  1^st^ r correlation effect size: 0.36  1^st^ Cohens d effect size: 0.78 | No significant Cluster |

The effect shown in Table 12 and Figure 31 clearly shows a decrease in amplitude through the partitions. This is most obvious in the grand-average (top left) that shows the PGI however, the effect is also present in the medium stimulus grand-average (bottom left). (Additionally, this decrease through the partitions interacted with Discomfort, but again only unorthogonalized; see section 9.2.2.5 in the appendix.)

This effect (and the corresponding interaction in the appendix) is curious since there is no prior precedence to support an effect of the pattern glare stimulus in this region on the scalp. Additionally, these pure change through time regressors are not orthogonalized with respect to the visual stress factor.

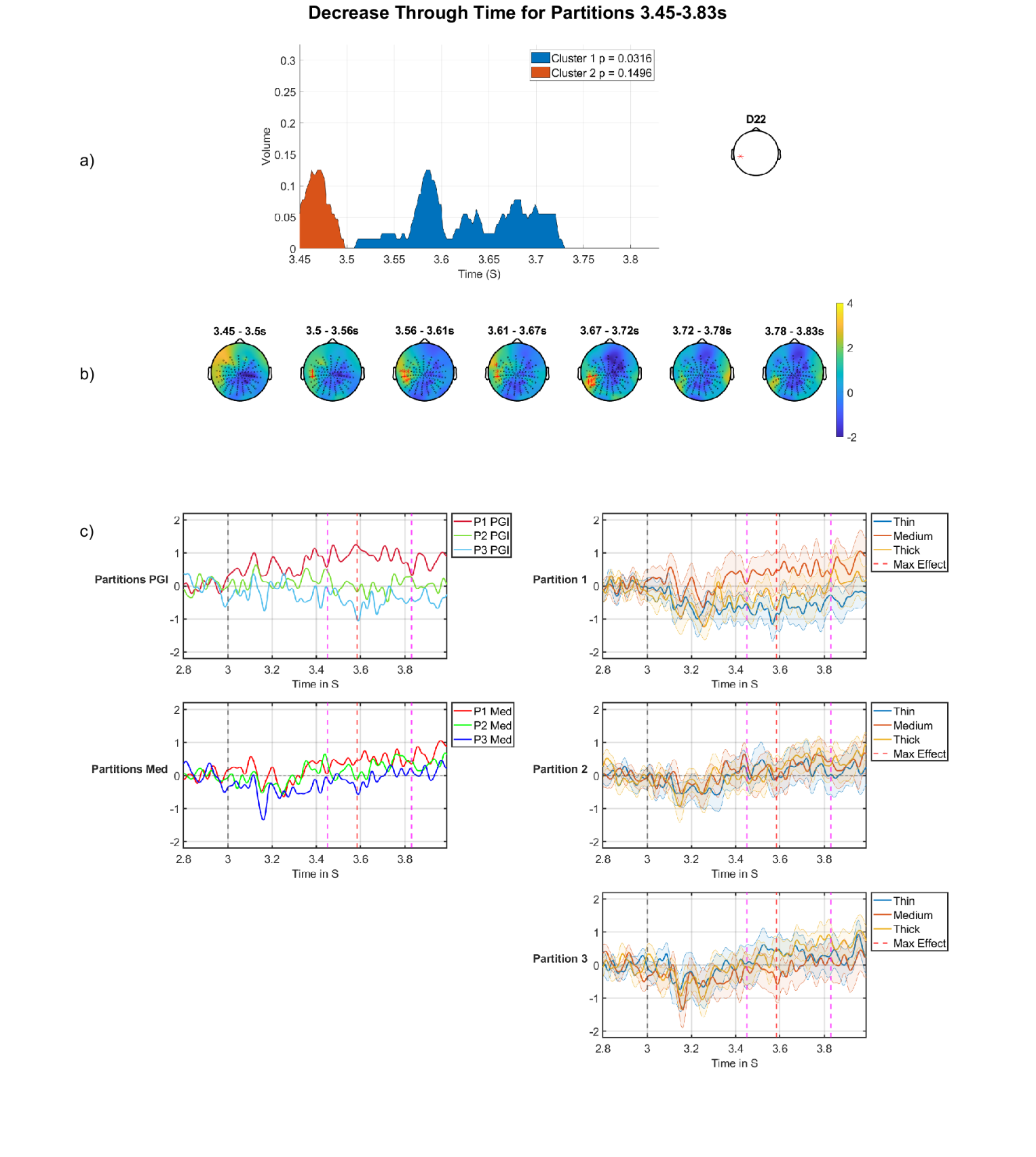

*Figure 31: pure decrease regressor across the partitions on the offset period. a) Cluster volume as a percentage of the entire scalp; the electrode used for plotting displayed on the right. b) Topographic maps through time for the whole period, with red crosses indicating the significant cluster, which corresponds to the blue region in a). c) grand-averages plotted at indicated electrode (D22), top left is the PGI through partitions, bottom left is medium stimulus through partitions, top right shows partition 1, middle right shows partition 2 and bottom right shows partition 3. Maximum effect is marked with a red dashed line, stimulus offset marked with a black dashed line and analysis windows are marked with dashed pink lines.*

***Orthogonalized Discomfort in onsets 2,3 vs 4,5 vs 6,7***

In this section, we look at the results of the MUA on the fine time granularity (onsets 2,3 vs 4,5 vs 6,7) to see if there is a habituation or sensitisation pattern through the onsets with block/partitions collapsed out. As before, we expect an habituation or sensitisation in three bins, so the same regressors are used as in the partitions analysis.

**Discomfort-by-decrease:** Now we look at discomfort by decrease on onsets 2,3 vs 4,5 vs 6,7. In Table 13 we can see the significant cluster. There is only one factor that came out as significant; this factor is discomfort and is found in the first window of analysis (3.09 – 3.18s).

*Table 13: MUA results for the discomfort by decrease regressor on the onsets 2,3 vs 4,5 vs 6,7. Only results for clusters containing significant effects (FWE-corrected) or borderline effects smaller than a p-value of 0.1 are shown, both positive and negative tails.*

| Effect | Tail (+1) | Tail (-1) |
| --- | --- | --- |
| Discomfort by decrease 3.09-3.18 | 1^st^ cluster p-value: 0.0056  1^st^ electrode: B15  1^st^ peak time: 3.1230  1^st^ r correlation effect size: 0.38  1^st^ Cohens d effect size: 0.83 | No significant Cluster |

The positive cluster shown in Table 13 and Figure 32 shows a positive going effect with p value of 0.0056 (FWE-corrected at cluster-level). The effect is in the right posterior region of the scalp, we have seen effects associated with discomfort throughout this paper, and at its maximum point it occupies just over 15% of the volume, as seen in Figure 32a. Figure 32b shows the clusters location through time with the cluster starting in a relatively anterior region before filling out and becoming more posterior, before moving more anteriorly again as it wanes. Figure 32c shows the median split (on discomfort) grand-averages. At the point of maximum effect, the low group shows a sensitisation effect, with onsets 2,3 having the least positive going PGI (top row, left) and the high group having almost the opposite with onsets 2,3 being the most positive going PGI (top row, right). The plots for medium in the second row show a similar pattern, but with a less extreme effect. Additionally, this effect closely tracks the pure discomfort factor effect in both space and time, compare panels a, b & c) in Figure 19 and Figure 32.

This effect actually exhibits an opposite pattern to the corresponding contrast during the DC-shift period, with Figure 14 showing habituation for the low group and sensitisation for the high group. Although it is potentially of interest, we have presented this effect in the appendix because it is difficult to reconcile with our main findings. This is unless it represents a polarity reversal arising from observing the underlying electrical dipole from opposite sides. In which case, the pattern on the other side of the dipole would be consistent with our main findings. We leave this for further research.
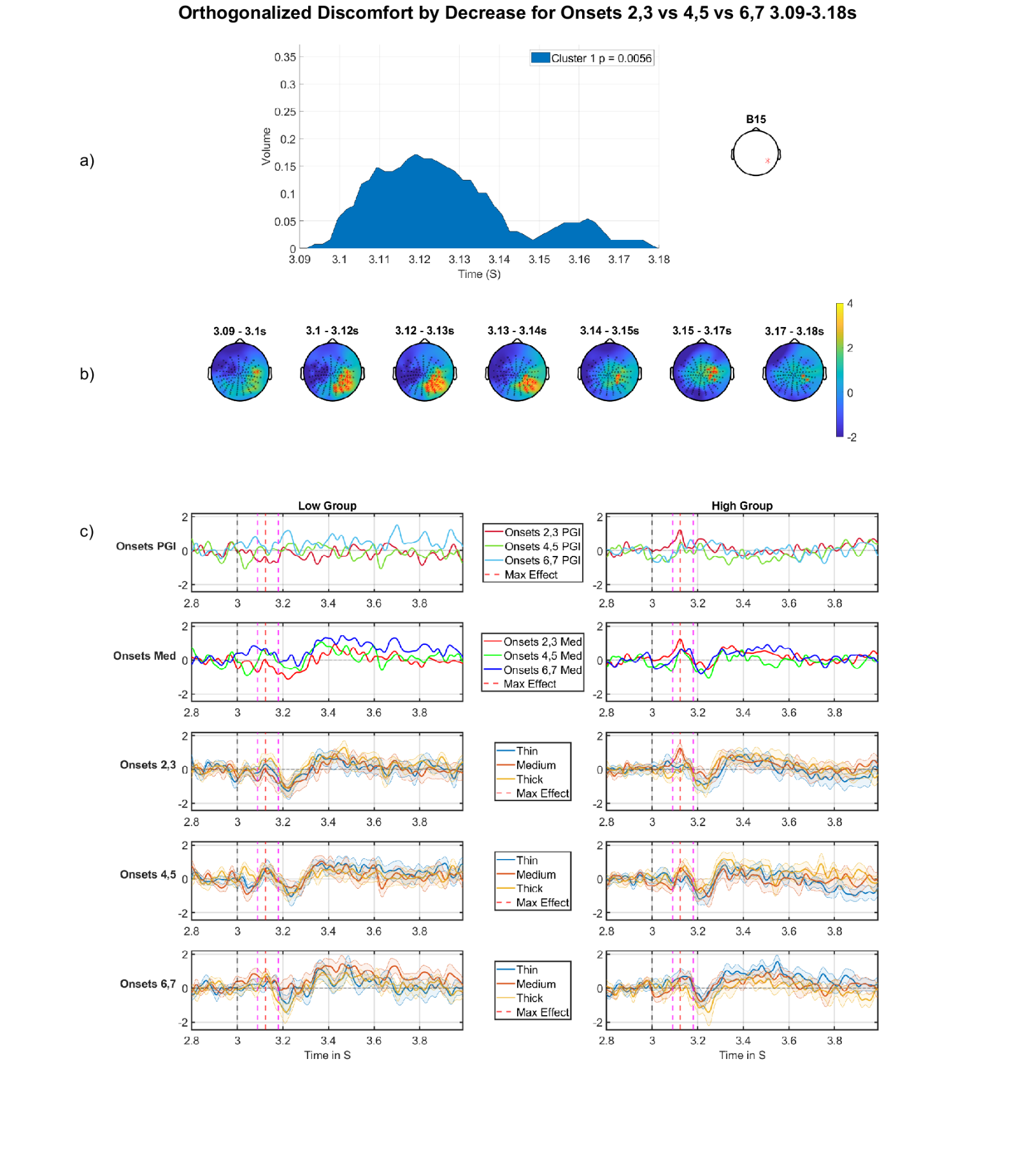

*Figure 32: Discomfort by decrease regressor on the offset period for onsets 2,3 vs 4,5 vs 6,7. a) Cluster volume as a percentage of the entire scalp. The electrode used for plotting is displayed on the right. b) Topographic maps through time for the whole period, with red crosses indicating the significant cluster, which corresponds to the blue region in a). c) Median split (on discomfort) grand-averages at electrode indicated on right in panel (a), the left panel of grand-averages is for the low group, right panel of grand-averages is for the high group. Top are the grand-averages for the PGI for each onset-pair, red (onsets 2,3), green (onsets 4,5), blue (onsets 6,7), with maximum effect marked with a red dashed vertical line, stimulus offset marked with a black dashed line and analysis window marked with pink dashed lines; second row is the grand-average for the medium stimulus for each onset-pair, red (onsets 2,3), green (onsets 4,5), blue (onsets 6,7); third, fourth and fifth rows present grand-averages for onsets 2,3, 4,5, and 6,7 (respectively), showing in each plot thin, medium and thick. The pattern of brain responses, which underlies this effect, can most easily be seen in the top row of panel c), where, at the point of maximum effect, we can see a sensitisation pattern for the low group and an decrease pattern for the high group. However, this decrease pattern in the high group is brief in time and substantially weaker when just looking at the medium stimulus in the 2nd row of panel c). This suggests that the effect is at least partially carried by changes in thin and thick through the onsets, indicating that this may not be as interesting an effect as some others in this paper.*

***Partitions Unorthogonalized results***

Discomfort-by-decrease across partitions for the offset period: the effect displayed in Table 14 shows a significant cluster found in the MUA for a discomfort by decrease effect. The effect here just crosses the threshold for significance (p=0.0462); however, it is in a region of the scalp where we have not seen any other effects and this effect is on the edge of the volume, which can be more susceptible to noise and artifacts.

But, this effect does suggest an habituation through the partitions, which is present for both PGI (top row of Figure 33c) and the medium stimulus (second row of Figure 33c). Furthermore, this reduction in amplitude is substantially greater for those in the high group.

*Table 14: MUA results for the discomfort factor by decrease across the partitions for the time period 3.45 – 3.83s. Only results for significant clusters are shown, both positive and negative tails.*

| Effect | Tail (+1) | Tail (-1) |
| --- | --- | --- |
| Discomfort by Decrease 3.45-3.83s | 1^st^ cluster p-value: 0.0462  1^st^ electrode: D22  1^st^ peak time: 3.7070  1^st^ r correlation effect size: 0.41  1^st^ Cohens d effect size: 0.90 | No significant Cluster |

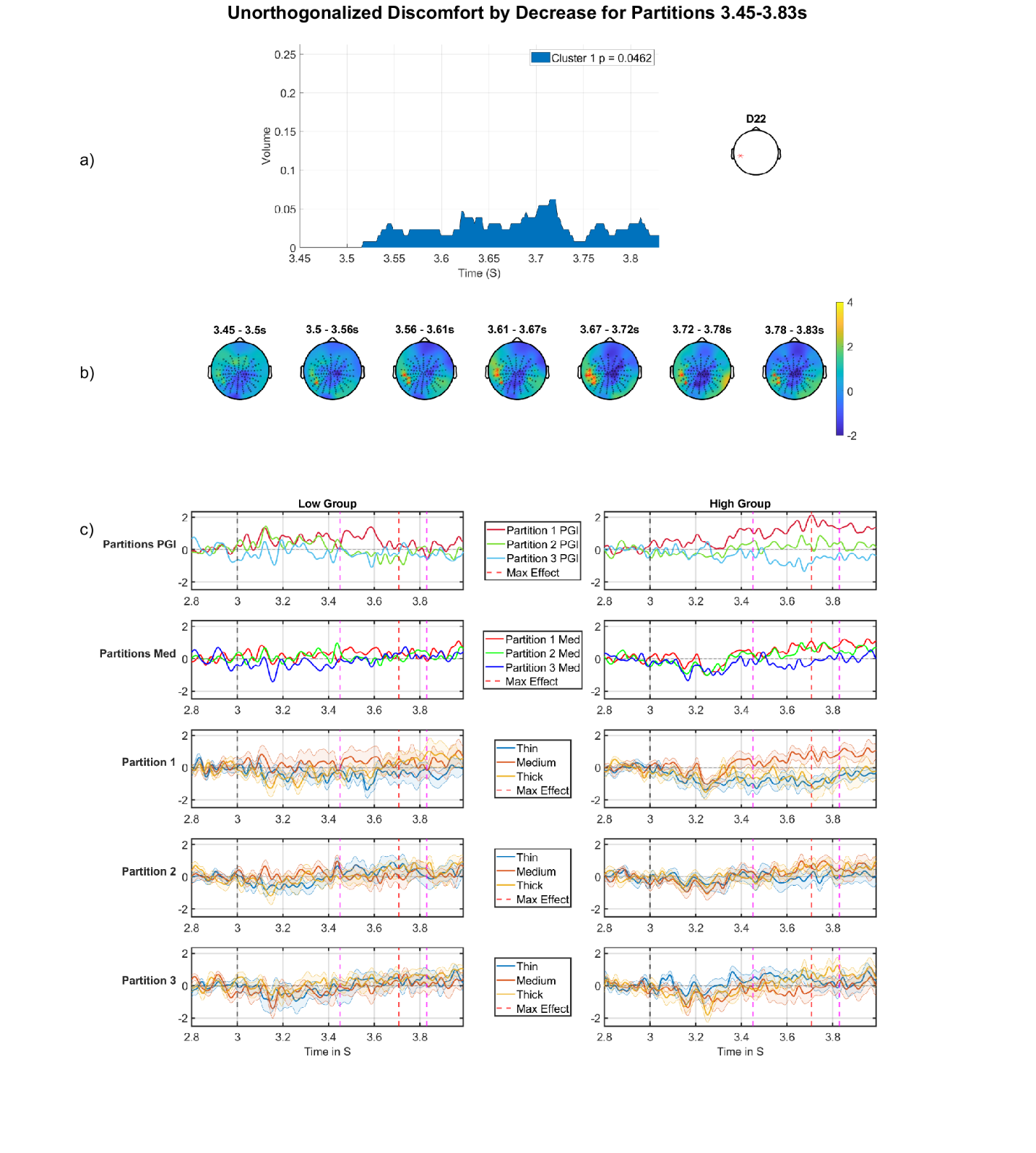

*Figure 33: Discomfort by decrease regressor across the partitions on the offset period. a) Cluster volume as a percentage of the entire scalp; the electrode used for plotting displayed on the right. b) Topographic maps through time for the whole period, with red crosses indicating the significant cluster, which corresponds to the blue region in a). c) Median split (on Discomfort) grand-averages at electrode indicated on right in panel (a). The left panel of grand-averages is for the low group, right panel of grand-averages is for the high group. Top are the grand-averages for the PGI for each partition, red (partition 1), green (partition 2), blue (partition 3), with maximum effect marked with a red vertical dashed line, window of analysis marked with pink dashed lines and stimulus offset marked with a black dashed line; second row are the grand-averages for the medium stimulus for each partition, red (partition 1), green (partition 2), blue (partition 3); third row are the grand-averages for partition 1, fourth row for partition 2, fifth row for partition 3, in each case, showing thin, medium and thick.*

**Effects for Visual stress and headache factors**

There were not many significant effects for the other two factors, however, in this section we will present those effects that were significant or tended towards significance.

***Onsets 2-8 headache factor in the DC shift period***

The MUA for the headache factor in the DC shift period revealed a close to significant result outlined in Table 15. This effect (p=0.0798) sits in a similar spatial location to the effects we have discovered for the discomfort factor; see Figure 34b. The effect shown occupies just over 10% of the volume at its maximum point. This effect, though not significant, does display a more positive PGI in the grand-average shown in Figure 34c top two panels on the left. This effect is also present for the medium stimulus shown in the third panel of Figure 34c. The effect starts towards the end of the period and is relatively small spatially, as shown in Figure 34a. Throughout the whole analysis period, the PGI and Medium stimulus for the high group are more positive than the low group, which could indicate that the high group’s response to the initial presentation does not return to a resting state in the way that it does for the low group does.

*Table 15: MUA results for headache on the average of onsets 2-8 orthogonalized regressor. Only results for clusters containing significant effects (FWE-corrected) or borderline effects smaller than a p-value of 0.1 are shown, both positive and negative tails.*

| Effect | Tail (+1) | Tail (-1) |
| --- | --- | --- |
| Headache 0.5 – 3.0s | 1^st^ cluster p-value: 0.0798  1^st^ electrode: B18  1^st^ peak time: 2.435  1^st^ r correlation effect size: 0.65  1^st^ Cohens d effect size: 1.73 | No significant Cluster |

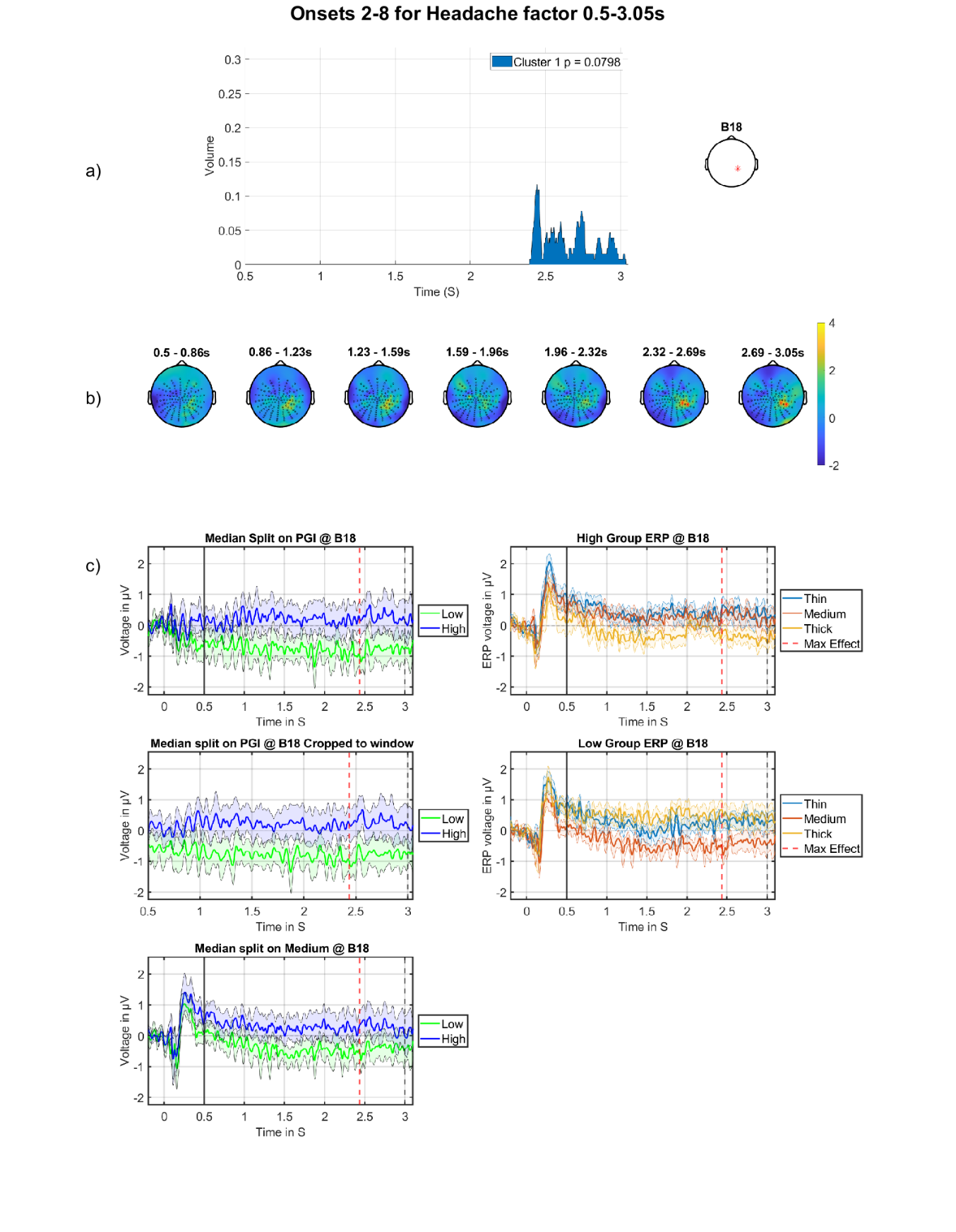

*Figure 34: Offset effect for the headache factor on the average of the onsets. a) Cluster volume as a percentage of the entire scalp. The electrode used for plotting is displayed on the right. b) Topographic maps through time for the whole period, with red crosses indicating the significant cluster, which corresponds to the blue region in a). c) grand-averages at electrode indicated on right in panel (a), with maximum effect marked with the red dashed vertical line, stimulus onset marked with a solid black line and stimulus offset marked with a black dashed line. Top left is the high vs low discomfort group for the PGI; middle left are the grand-averages for high vs low discomfort group for the PGI with window showing only the period of analysis; bottom left are the high vs low group for the medium stimulus; top right are the grand-averages for the high group showing thick, medium and thin; bottom right are the grand-averages for the low group showing thick, medium and thin.*

***Three-way, headache factor in the DC shift period***

We now look at the three-way interaction for the headache factor with a decrease in the partitions and an increase in the onsets. This shows a significant positive going effect with p=0.0326; see Table 16. As shown in Figure 35b, the effect sits in the same spatial location as those seen in Figure 34b and is also temporally similar, with the cluster occurring at the end of the DC shift period; see Figure 35a. The grand-averages are not as convincing for this effect; see Figure 35c, which seem to be mostly carried by the last partition in which we see a reversal in polarity between low and high groups.

*Table 16: MUA results for the three-way interaction headache by decrease in the partitions by increase in the onsets. Only results for clusters containing significant effects (FWE-corrected) or borderline effects smaller than a p-value of 0.1 are shown, both positive and negative tails.*

| Effect | Tail (+1) | Tail (-1) |
| --- | --- | --- |
| Headache by decrease in the partitions by increase in the onsets 0.5 – 3.0 | 1^st^ cluster p-value: 0.0326  1^st^ electrode: B6  1^st^ peak time: 2.871  1^st^ r correlation effect size: 0.25  1^st^ Cohens d effect size: 0.52  2^nd^ cluster p-value: 0.0812  2^nd^ electrode: B19  2^nd^ peak time: 0.966  2^nd^ r correlation effect size: 0.27  2^nd^ Cohens d effect size: 0.55 | No significant Cluster |

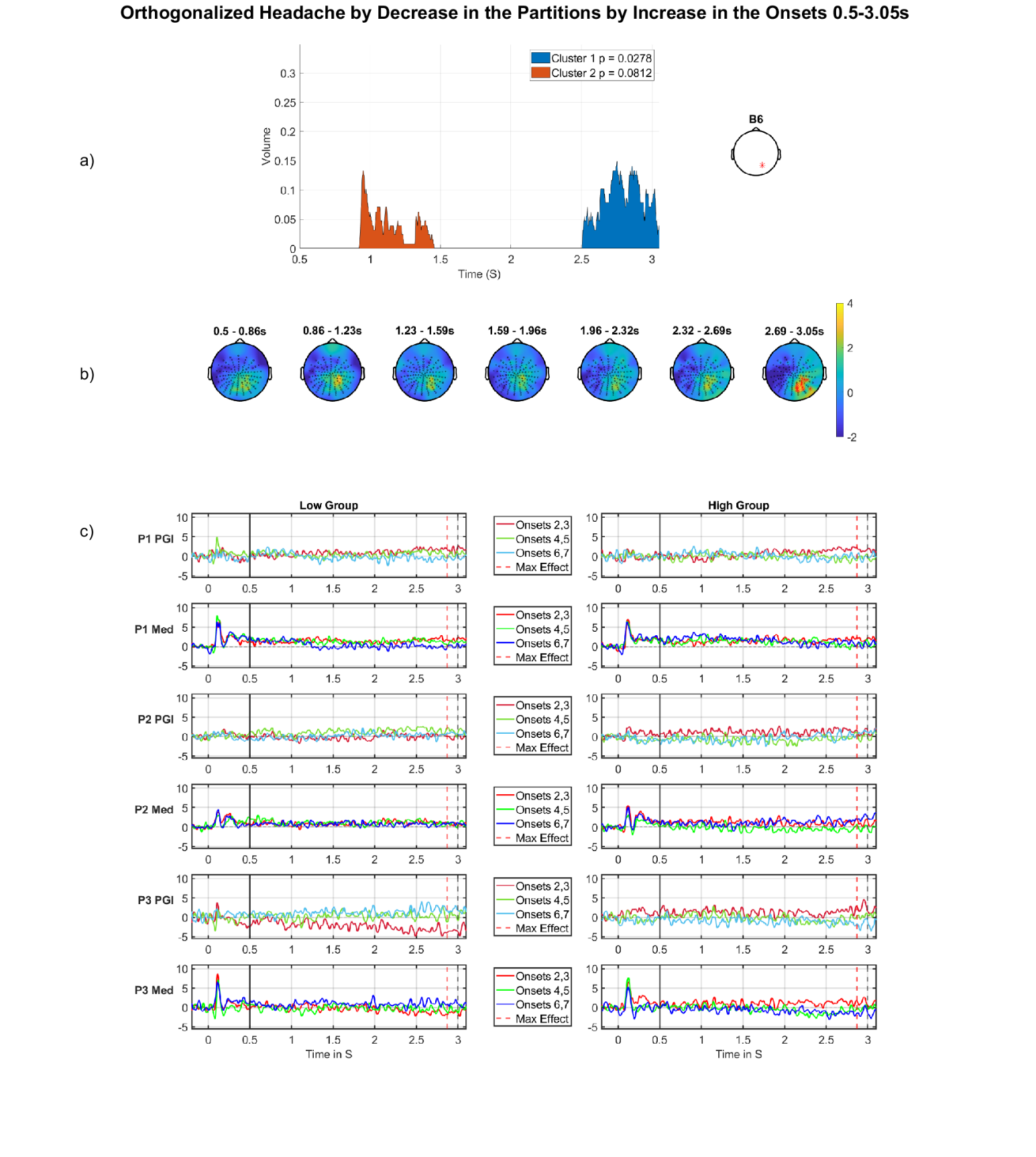

*Figure 35: Headache by decrease (through the partitions) by increase (through the onsets), DC shift period. a) Cluster volume as a percentage of the entire scalp, the electrode used for plotting displayed on the right. b) Topographic maps through time for the whole period, with red crosses indicating the significant cluster, which corresponds to the blue region in a). c) Median split (on Headache) grand-averages at electrode indicated on right in panel (a); the left panel of grand-averages is for the low Headache group, right panel of grand-averages is for the high group. Top row (partition 1), third row (partition 2) and fifth row (partition 3) show grand-averages for the PGI for each onset, red (onsets 2,3), green (onsets 4,5), blue (onsets 6,7) with maximum effect marked with a red vertical line, window start marked with a black solid line and stimulus offset marked with a black dashed line. Second row (partition 1), fourth row (partition 2) and sixth row (partition 3) are the grand-averages for just the medium stimulus for each onset, red (onsets 2,3), green (onsets 4,5), blue (onsets 6,7).*

***Onsets 2-8 Visual Stress factor in the Offset (3.45 – 3.83s)***

We now look at the average of onsets 2-8 for the visual stress factor. This effect occurs in the final window of the offset period (3.45 – 3.83s) shown in Table 17 and has a p value of 0.053, which is above the significance threshold of 0.05. This effect sits in the later part of the window, lasting over 100ms and occupying just over 10% of the volume shown in Figure 36a. The effect is in a similar spatial location to the effects seen for discomfort in the offset period: compare Figure 19b and Figure 36b. A relatively sustained higher amplitude for the high group is evident throughout the time windows of the top two panels of Figure 36c. This pattern is also somewhat present for the medium stimulus; see third panel of Figure 36c. This combination of features suggest a potential hyper excitation for those susceptible to visual stress, although in this data set it was not strong enough to reach significance.

*Table 17: MUA results for* *visual stress on the average of onsets 2-8 orthogonalized regressor. Only results for clusters containing significant effects (FWE-corrected) or borderline effects smaller than a p-value of 0.1 are shown, both positive and negative tails.*

| Effect | Tail (+1) | Tail (-1) |
| --- | --- | --- |
| Visual Stress 3.45 – 3.83s | 1^st^ cluster p-value: 0.053  1^st^ electrode: B10  1^st^ peak time: 3.7012  1^st^ r correlation effect size: 0.72  1^st^ Cohens d effect size: 2.05 | No significant Cluster |

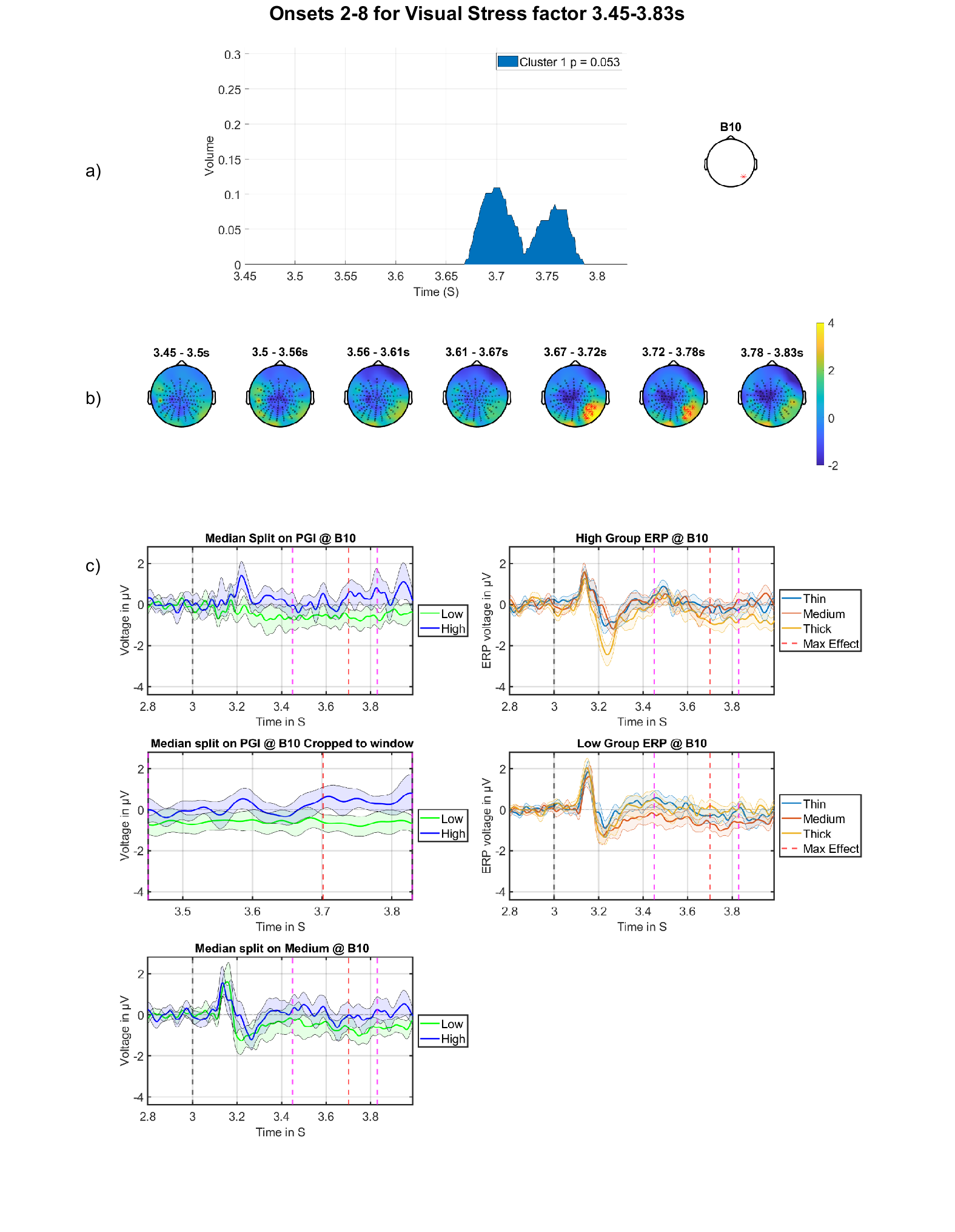

*Figure 36: Offset effect for the visual stress factor on the average of the onsets. a) Cluster volume as a percentage of the entire scalp. The electrode used for plotting is displayed on the right. b) Topographic maps through time for the whole period, with red crosses indicating the significant cluster, which corresponds to the blue region in a). c) grand-averages at electrode indicated on right in panel (a), with maximum effect is marked with a red dashed line, stimulus offset marked with a black dashed line and analysis windows are marked with dashed pink lines. Top left is the high vs low visual stress group for the PGI; middle left are the grand-averages for high vs low visual stress group for the PGI with window showing only the period of analysis; bottom left are the high vs low group for the medium stimulus; top right are the grand-averages for the high group showing thick, medium and thin; bottom right are the grand-averages for the low group showing thick, medium and thin.*

***Discussion for Visual stress and headache factors***

**Headache:** Of the three effects presented for the non-discomfort factors (visual stress and headache) only the three-way interaction for headache crossed the threshold for significance at the second level. This effect is not very compelling when considering the grand average plots (Figure 35c) and looks like it is mostly driven by the order reversal of the onsets in the third partition, where the low group displays an increasing voltage across the onsets and the high group displays the reverse pattern. The two-way interaction did not find any clusters and therefore is not presented. The headache effect for the average of onsets approached significance (p value of 0.0798). This effect does however, span a similar time period to that of the three-way cluster and at a similar location in the volume. For the effect for the average of onsets, the low group’s medium grand average (green line Figure 34c, third row) is more negative-going than the high group’s. This is consistent with the effect for the three-way interaction, which could suggest a common underlying effect on our headache factor, however, more analysis is required to confirm this.

**Visual Stress:** The effect shown for visual stress occurs at the end of the offset period; see grand average plots in Figure 36c. The PGI plots for this figure show the high group having a heightened PGI to the stimulus offset, however, for the grand average plots for the three stimuli (right side of panel), the salient feature is a more negative going medium during the duration of the cluster. This can also be seen in the median split on medium; see bottom panel (on left). This may be an interesting finding, but since it was not significant and no other clusters for visual stress were identified during the DC shift or offset period, we have not interpreted this further. Another study is required to see if the finding replicates.

**Full MUA results**

In this section we will summarise all the MUA tests and if the results yielded any clusters.

The tables are split into sections, the first section contains the results for the DC shift period. The first table is for the orthogonalized MUA and second is the unorthogonalized MUA results. The next section is the Offset period; this is split into subsections for each of the time windows used in the MUA, following the same pattern as those in the DC shift period.

The Table contains three different possibilities for results. (Note, the possibility that no sample/time-space point crossed the first level statistical threshold, did not happen for any of our contrasts; thus, for all of these contrasts clusters were formed.)

1. Significant cluster reported in the paper, with section and table presenting results.
2. **NSC**: Clusters found but did not pass 2^nd^ level threshold for significance.
3. **N/A**: No MUA was run for this combination.

***DC-shift period***

**Orthogonalized Results**

*Table 18: DC shift MUA summary for orthogonalized regressors*

| Type/Factor | Mean/Intercept | Pure Change through time | Discomfort | Visual Stress | Headache |
| --- | --- | --- | --- | --- | --- |
| Onsets 2-8 | See 3.1.1.1, Table 1 | N/A | See 9.2.1.1, Table 9 | NSC | See 9.2.3.1, Table 15 |
| Orthogonalized partitions, Increase | N/A | NSC | See 3.1.2.1, Table 3 | NSC | NSC |
| Orthogonalized partitions, Decrease | N/A | NSC | See 3.1.2.1, Table 2 | NSC | NSC |
| Orthogonalized  Onsets 2,3 vs 4,5 vs 6,7, Increase | N/A | NSC | NSC | NSC | NSC |
| Orthogonalized  Onsets 2,3 vs 4,5 vs 6,7, Decrease | N/A | NSC | See 3.1.3.1, Table 4 | NSC | NSC |
| Three-way Interaction, Increase in Partitions, Decrease in onsets | N/A | N/A | See 3.1.4, Table 5 | NSC | NSC |
| Three-way Interaction, Increase in Partitions, Decrease in onsets | N/A | N/A | NSC | NSC | See 9.2.3.2, Table 16 |

**Unorthogonalized Results**

*Table 19: DC Shift MUA summary for unorthogonalized regresors*

| Type/Factor | Mean/Intercept | Pure Change through time | Discomfort | Visual Stress | Headache |
| --- | --- | --- | --- | --- | --- |
| Unorthogonalized partitions, Increase | N/A | N/A | NSC | N/A | NSC |
| Unorthogonalized partitions, Decrease | N/A | N/A | NSC | N/A | NSC |
| Unorthogonalized Onsets 2,3 vs 4,5 vs 6,7, Increase | N/A | N/A | NSC | N/A | NSC |
| Unorthogonalized Onsets 2,3 vs 4,5 vs 6,7, Decrease | N/A | N/A | See 9.2.1.2, Table 10 | N/A | NSC |

***Offset Period***

**3.09-3.99s Orthogonalized Results**

*Table 20: Offset period 3.09-3.99s MUA summary for othogonalized regressors*

| Type/Factor | Mean/Intercept | Pure Change through time | Discomfort | Visual Stress | Headache |
| --- | --- | --- | --- | --- | --- |
| Onsets 2-8 | See 9.2.2.1, Table 11 | N/A | NSC | NSC | NSC |
| Orthogonalized partitions, Increase | N/A | NSC | NSC | NSC | NSC |
| Orthogonalized partitions, Decrease | N/A | NSC | NSC | NSC | NSC |
| Orthogonalized  Onsets 2,3 vs 4,5 vs 6,7, Increase | N/A | NSC | NSC | NSC | NSC |
| Orthogonalized  Onsets 2,3 vs 4,5 vs 6,7, Decrease | N/A | NSC | NSC | NSC | NSC |
| Three-way Interaction, Increase in Partitions, Decrease in onsets | N/A | N/A | NSC | NSC | NSC |
| Three-way Interaction, Increase in Partitions, Decrease in onsets | N/A | N/A | NSC | NSC | NSC |

**3.09-3.99s Unorthogonalized Results**

*Table 21: Offset period 3.09-3.99s MUA summary for unothogonalized regressors*

| Type/Factor | Mean/Intercept | Pure Change through time | Discomfort | Visual Stress | Headache |
| --- | --- | --- | --- | --- | --- |
| Unorthogonalized partitions, Increase | N/A | N/A | NSC | N/A | NSC |
| Unorthogonalized partitions, Decrease | N/A | N/A | NSC | N/A | NSC |
| Unorthogonalized Onsets 2,3 vs 4,5 vs 6,7, Increase | N/A | N/A | NSC | N/A | NSC |
| Unorthogonalized Onsets 2,3 vs 4,5 vs 6,7, Decrease | N/A | N/A | NSC | N/A | NSC |

**3.09-3.18s Orthogonalized Results**

*Table 22: Offset period 3.09-3.18s MUA summary for othogonalized regressors*

| Type/Factor | Mean/Intercept | Pure Change through time | Discomfort | Visual Stress | Headache |
| --- | --- | --- | --- | --- | --- |
| Onsets 2-8 | See 9.2.2.1, Table 11 | N/A | See 3.2.1.2, Table 6 | NSC | NSC |
| Orthogonalized partitions, Increase | N/A | NSC | NSC | NSC | NSC |
| Orthogonalized partitions, Decrease | N/A | NSC | See 3.2.2.2, Table 7 | NSC | NSC |
| Orthogonalized  Onsets 2,3 vs 4,5 vs 6,7, Increase | N/A | NSC | NSC | NSC | NSC |
| Orthogonalized  Onsets 2,3 vs 4,5 vs 6,7, Decrease | N/A | NSC | See 3.2.2.2, Table 13 | NSC | NSC |
| Three-way Interaction, Increase in Partitions, Decrease in onsets | N/A | NSC | NSC | NSC | NSC |
| Three-way Interaction, Increase in Partitions, Decrease in onsets | N/A | NSC | NSC | NSC | NSC |

**3.09-3.18s Unorthogonalized Results**

*Table 23: Offset period 3.09-3.18s MUA summary for unothogonalized regressors*

| Type/Factor | Mean/Intercept | Pure Change through time | Discomfort | Visual Stress | Headache |
| --- | --- | --- | --- | --- | --- |
| Unorthogonalized partitions, Increase | N/A | N/A | NSC | N/A | NSC |
| Unorthogonalized partitions, Decrease | N/A | N/A | NSC | N/A | NSC |
| Unorthogonalized Onsets 2,3 vs 4,5 vs 6,7, Increase | N/A | N/A | NSC | N/A | NSC |
| Unorthogonalized Onsets 2,3 vs 4,5 vs 6,7, Decrease | N/A | N/A | NSC | N/A | NSC |

**3.18-3.45s Orthogonalized Results**

*Table 24: Offset period 3.18-45s MUA summary for othogonalized regressors*

| Type/Factor | Mean/ Intercept | Pure Change through time | Discomfort | Visual Stress | Headache |
| --- | --- | --- | --- | --- | --- |
| Onsets 2-8 | See 9.2.2.1, Table 11 | N/A | NSC | NSC | NSC |
| Orthogonalized partitions, Increase | N/A | NSC | NSC | NSC | NSC |
| Orthogonalized partitions, Decrease | N/A | NSC | NSC | NSC | NSC |
| Orthogonalized  Onsets 2,3 vs 4,5 vs 6,7, Increase | N/A | NSC | NSC | Significant Cluster found however, borderline, and not in a region of interest, thus not reported | NSC |
| Orthogonalized  Onsets 2,3 vs 4,5 vs 6,7, Decrease | N/A | NSC | NSC | NSC | NSC |
| Three-way Interaction, Increase in Partitions, Decrease in onsets | N/A | NSC | NSC | NSC | NSC |
| Three-way Interaction, Increase in Partitions, Decrease in onsets | N/A | NSC | NSC | NSC | NSC |

**3.18-3.45s Unorthogonalized Results**

*Table 25: Offset period 3.18-3.45s MUA summary for unothogonalized regressors*

| Type/Factor | Mean/Intercept | Pure Change through time | Discomfort | Visual Stress | Headache |
| --- | --- | --- | --- | --- | --- |
| Unorthogonalized partitions, Increase | N/A | N/A | NSC | N/A | NSC |
| Unorthogonalized partitions, Decrease | N/A | N/A | NSC | N/A | NSC |
| Unorthogonalized Onsets 2,3 vs 4,5 vs 6,7, Increase | N/A | N/A | NSC | N/A | NSC |
| Unorthogonalized Onsets 2,3 vs 4,5 vs 6,7, Decrease | N/A | N/A | NSC | N/A | NSC |

**3.45-3.83 Orthogonalized Results**

*Table 26: Offset period 3.45-3.83s MUA summary for othogonalized regressors*

| Type/Factor | Mean/Intercept | Pure Change through time | Discomfort | Visual Stress | Headache |
| --- | --- | --- | --- | --- | --- |
| Onsets 2-8 | See 9.2.2.1, Table 11 | N/A | NSC | See 9.2.3.3, Table 17 | NSC |
| Orthogonalized partitions, Increase | N/A | NSC | NSC | NSC | NSC |
| Orthogonalized partitions, Decrease | N/A | See 9.2.2.3, Table 12 | NSC | NSC | NSC |
| Orthogonalized  Onsets 2,3 vs 4,5 vs 6,7, Increase | N/A | NSC | NSC | NSC | NSC |
| Orthogonalized  Onsets 2,3 vs 4,5 vs 6,7, Decrease | N/A | NSC | NSC | NSC | NSC |
| Three-way Interaction, Increase in Partitions, Decrease in onsets | N/A | NSC | NSC | NSC | NSC |
| Three-way Interaction, Increase in Partitions, Decrease in onsets | N/A | NSC | NSC | NSC | NSC |

**3.45-3.83s Unorthogonalized Results**

*Table 27: Offset period 3.45-3.83s MUA summary for unothogonalized regressors*

| Type/Factor | Mean/Intercept | Pure Change through time | Discomfort | Visual Stress | Headache |
| --- | --- | --- | --- | --- | --- |
| Unorthogonalized partitions, Increase | N/A | N/A | NSC | N/A | NSC |
| Unorthogonalized partitions, Decrease | N/A | N/A | See 9.2.2.5, Table 14 | N/A | NSC |
| Unorthogonalized Onsets 2,3 vs 4,5 vs 6,7, Increase | N/A | N/A | NSC | N/A | NSC |
| Unorthogonalized Onsets 2,3 vs 4,5 vs 6,7, Decrease | N/A | N/A | NSC | N/A | NSC |
